# *Lhx2* in germ cells suppresses endothelial cell migration in the developing ovary

**DOI:** 10.1101/2022.03.07.483280

**Authors:** Neha Singh, Domdatt Singh, Anshul Bhide, Richa Sharma, Sarthak Sahoo, Mohit Kumar Jolly, Deepak Modi

**Affiliations:** Molecular and Cellular Biology Laboratory, ICMR-National Institute for Research in Reproductive and Child Health, Indian Council of Medical Research (ICMR), JM Street, Parel, Mumbai 400012, India; Center for BioSystems Science and Engineering, Indian Institute of Science, CV Raman Rd, Bangalore, 560012, India

**Author notes:** Corresponding author Dr. Deepak Modi, Scientist F and Head, Molecular and Cellular Biology Laboratory, ICMR-National Institute for Research in Reproductive and Child Health, Indian Council of Medical Research (ICMR), J.M. Street, Parel’, Mumbai 400 012 India.

**Keywords:** Homeobox, LIM gene, LIM-HD, Sex determination, Gonad development, Testis, Endothelial cell migration, Angiogenesis, Vasculature

## Abstract

LIM-homeobox genes play multiple roles in developmental processes, but their roles in gonad development are not completely understood. Herein, we report that *Lhx2, Ils2, Lmx1a*, and *Lmx1b* are expressed in a sexually dimorphic manner in mouse, rat, and human gonads during sex determination. Amongst these, *Lhx2* has female biased expression in the developing gonads of species with environmental and genetic modes of sex determination. Single-cell RNAseq analysis revealed that *Lhx2* is exclusively expressed in the germ cells of the developing mouse ovaries. To elucidate the roles of *Lhx2* in the germ cells, we analyzed the phenotypes of *Lhx2* knockout XX gonads. While the gonads developed appropriately in *Lhx2* knockout mice and the somatic cells were correctly specified in the developing ovaries, transcriptome analysis revealed enrichment of genes in the angiogenesis pathway. There was an elevated expression of several pro-angiogenic factors in the *Lhx2* knockout ovaries. The elevated expression of pro-angiogenic factors was associated with an increase in numbers of endothelial cells in the *Lhx2*-/-ovaries at E13.5. Gonad recombination assays revealed that the increased numbers of endothelial cells in the XX gonads in absence of *Lhx2* was due to ectopic migration of endothelial cells in a cell non-autonomous manner. We also found that, there was increased expression of several endothelial cell-enriched male-biased genes in *Lhx2* knockout ovaries. Also, in absence of *Lhx2*, the migrated endothelial cells formed an angiogenic network similar to that of the wild type testis, although the coelomic blood vessel did not form. Together, our results suggest that *Lhx2* in the germ cells is required to suppress vascularization in the developing ovary. These results suggest a need to explore the roles of germ cells in the control of vascularization in developing gonads.

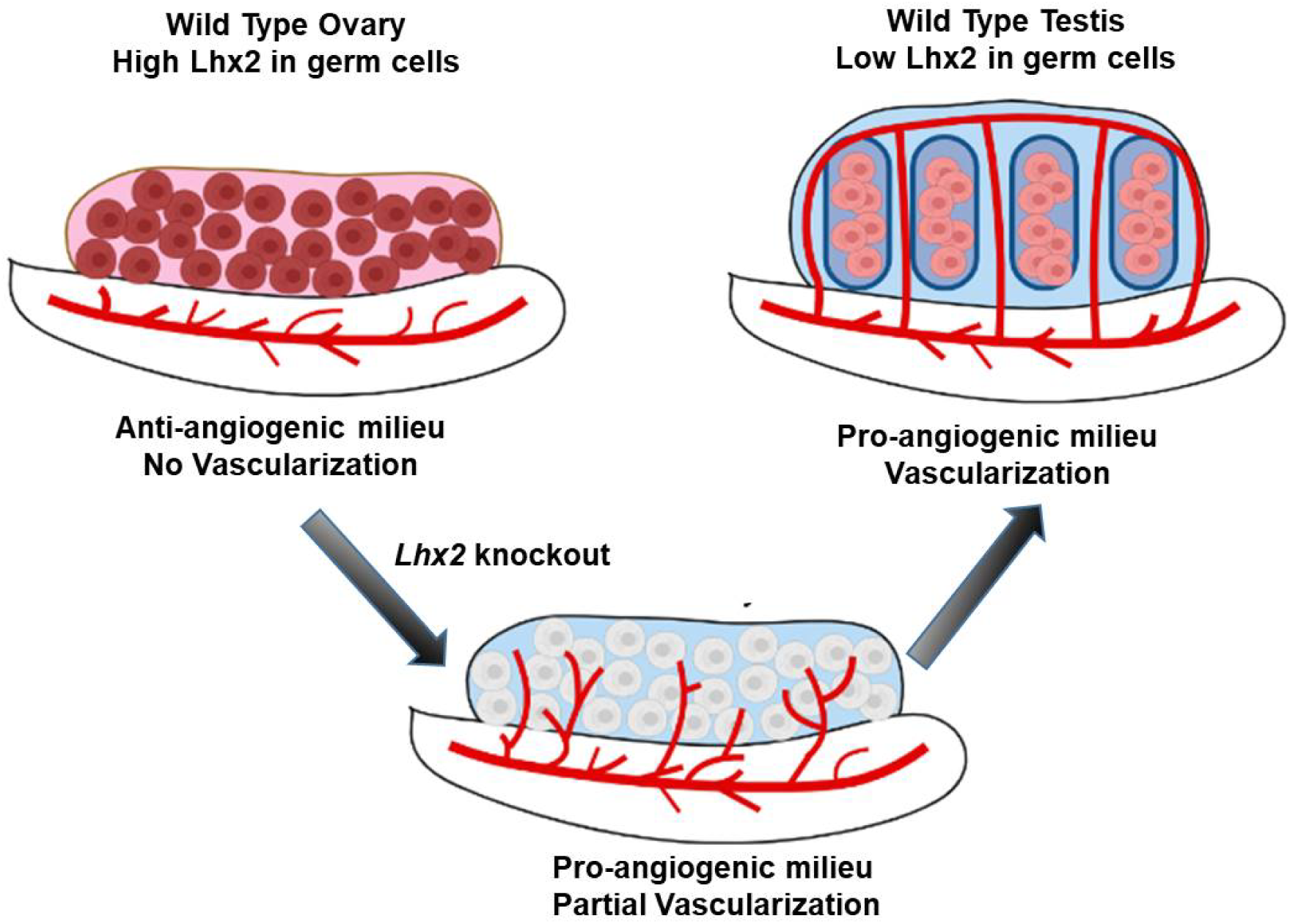

**Highlight:**

- Multiple LIM-HD genes are expressed in developing gonads during the window of sex determination with *Lhx2* having female dominating expression in an evolutionarily conserved manner
- *Lhx2* is expressed in the germ cells of developing mouse ovaries
- Loss of *Lhx2* in the developing ovaries alters the expression of genes involved in various pathways including angiogenesis
- *Lhx2* in germ cells suppress endothelial cell migration in the developing ovaries

## Introduction

In most sexually dimorphic species, the gonads develop as a bipotential urogenital ridge that differentiates either into the testis or an ovary depending on genetic and/or environmental cues. The coelomic epithelium from the ventral surface of mesonephros proliferates and provides the somatic niche; the primordial germ cells migrate and colonize to form the gonad. In the mouse, *Sry* gene is induced in the XY gonads around E10.5 that leading to the differentiation of somatic cells to Sertoli cells (Ortega et al., 2019; Rotgers et al., 2018). Under a well-defined genetic network, the Sertoli cells organize to form seminiferous tubules which is the hallmark of testicular differentiation (Nef et al., 2019; Rotgers et al., 2018; Singh and Modi, 2020; Yildirim et al., 2020). Along with somatic cell differentiation, there is sex-specific vascularization. In the XY gonads at E12.5, the mesonephric vascular plexus breaks down releasing the mesonephric endothelial cells which migrate into the testis establishing a vascular network and forming a coelomic blood vessel (Coveney et al., 2008; Svingen and Koopman, 2013; Ungewitter and Yao, 2013). Signalling factors like *Fgfs, Pdgf*, and *Vegf* aid in endothelial migration and male-specific blood vascular patterning (Cool et al., 2011; Gu et al., 2021; Svingen and Koopman, 2013). Vascularization is crucial for the delivery of exogenous factors in the testicular compartments, in the initial organization of the testis cords, and also in the regulation of testis morphogenesis (Gu et al., 2021; Singh and Modi, 2020; Svingen and Koopman, 2013). It is also required for maintaining the interstitial progenitor population of the fetal Leydig cells (Gu et al., 2021; Kumar and DeFalco, 2018; Li et al., 2021).

During development, angiogenesis is specific to the XY gonads as endothelial cell migration and vascularization do not occur in the XX gonads. Although vascular endothelial cells are detected in the fetal ovary, these do not arise from the mesonephros and neither forms the vascular network, rather the vasculature grows by pre-existing branches and its reorganization (Coveney et al., 2008). However, ectopic migration of endothelial cells from the mesonephros is detected in the fetal ovaries of mice knockout for *Rspo1, Wnt4, Ctnnb1*, or *Fst*, and the endothelial cells also form a coelomic blood vessel similar to that observed in the testis (Chassot et al., 2014; Nicol and Yao, 2015; Tang et al., 2020; Yao et al., 2006). These results indicate that although a genetic program is required to initiate vascularization in the XY gonads, there also exists a genetic program to suppress vascularization in the developing XX gonads. While the biological need to suppress vascularization in the developing ovary is not clear, ectopic vascularization is accompanied by differentiation of Sertoli-like cells and fetal Leydig cells in the developing XX gonads (Chassot et al., 2014; Tang et al., 2020). In the testis too, disrupted vascularization promotes premature differentiation of fetal Leydig cells (Kumar and DeFalco, 2018). These observations imply that suppression of vascularization may be required to prevent masculinization of the fetal ovary. However, beyond *Wnt4, Rspo1, Ctnnb1*, and *Fst*; factors (if any) that suppress vascularization in the fetal ovary are unknown.

LIM homeobox (Lhx) genes encode for transcription factors that contain two tandem N terminal LIM domains and centrally located a DNA binding homeodomain (HD). The LIM homeodomain (LIM-HD) proteins execute their functions through protein-protein interactions mediated by LIM domains and the homeodomain controls gene expression (Singh et al., 2021; Yasuoka and Taira, 2021). During embryonic development, LIM-HD genes are crucial regulators of cell proliferation, cell specification, migration, and body axis patterning (Chou and Tole, 2019; Hunter and Rhodes, 2005; Singh et al., 2021; Yasuoka and Taira, 2021). Amongst the LIM-HD genes, *Lhx1* is known for its role in the development of the Mullerian tract (Huang et al., 2014), *Lhx8* is required for oocyte development after the germ cells enter meiosis (Choi et al., 2008) and *Lhx9* is essential for the development of bipotential gonads (Birk et al., 2000). We have also shown that multiple members of the LIM-HD gene family and their co-regulators are dynamically expressed in the developing mouse gonads specifically during the window of sex determination (Singh et al., 2021). However, the functional role of most LIM-HD family genes in gonad development and differentiation has not been explored.

In the present study, we investigated the expression profiles of *LIM-HD* genes in the gonads of multiple species and determined the role of one of the *LIM-HD* genes, *Lhx2* in gonad development. Our results revealed that amongst the *LIM-HD* genes, expression of *Lhx2* is sexually dimorphic with higher levels in the female gonads during sex determination in multiple species. We further observed that loss of *Lhx2* in the XX mouse gonads leads to ectopic migration of the endothelial cells without causing somatic cell sex reversal.

## Materials and Methods

### Expression profiling of *LIM-HD* genes in the developing gonads

To investigate the expression pattern of LIM homeobox genes in developing gonads of different species, publicly available bulk RNA seq datasets of mice, rats and humans were analyzed (Cardoso-Moreira et al., 2019). To determine the expression profiles of *Lhx2* in species with different modes of sex determination, we analyzed bulk RNAseq datasets of lizard (Whiteley et al., 2021), turtle (Czerwinski et al., 2016), chicken, and opossum (Cardoso-Moreira et al., 2019). Levels of *Lhx2* in mice and humans were also analyzed in two additional datasets (Lecluze et al., 2020; Zhao et al., 2018). The accession numbers of these datasets are given in Supplementary Table 1.

### Determining the cell types that express *Lhx2*

To determine the cell types in the mouse gonads that express *Lhx2*, we analyzed its mRNA expression single-cell RNAseq dataset (GSE128553) of mouse gonads at E13.5 (Ge et al., 2021). This dataset contains information on 4,842 cells from the XX gonads at E13.5. We performed t-distributed Stochastic Neighbor Embedding (tSNE) and clustering on all the cells in the dataset to identify various cell types. After the tSNE projection, we used *Kdr* and *Flt1* as specific markers to identify the endothelial cell population, the germ cell population was visualized using the classical germ cell markers *Ddx4* and *Dazl*. To visualize the expression of selected genes, we coloured the tSNE plots by imputed gene expression as computed by the MAGIC algorithm (van Dijk et al., 2018).

### *Lhx2* knockout mice

The study was approved by the institutional animal ethics committee (IAEC Project No., 27/14, 2/16). Heterozygous male and female mice with a targeted disruption of *Lhx2* (Porter et al., 1997) and those with conditional *Lhx2* allele (Mangale et al., 2008) were used in this study. The *Lhx2*^flox/flox^ mice crossed with mice expressing Cre under tamoxifen responsive estrogen receptor promoter (Supplementary Fig 1) were used to conditionally delete the *Lhx2* allele. All these strains were kindly provided by Prof. Shubha Tole, TIFR, Mumbai, India. The strains were bred at the experimental animal facility of ICMR-NIRRCH. Animals were mated in a 2:1 ratio and the day of vaginal plug observed was considered as embryonic day (E) 0.5.

The strategy of obtaining *Lhx2*^flox/flox^ pups is shown in Supplementary Fig 1. The *Lhx2*^flox/flox^ pregnant females were fed orally with a single dose of 10mg/ml of tamoxifen (Sigma Aldrich; MO, USA) micronized in corn oil (Sigma Aldrich) on E9.5. This dose and time were chosen as it is known to flox *Lhx2* allele within 24h (Mangale et al., 2008). Preliminary experiments conducted in the gonadal tissue also showed adequate floxing and down regulation of *Lhx2* mRNA at this dose and time (Supplementary Fig 1). As controls, *Lhx2*^flox/flox^ mice without the tamoxifen-inducible CreER were fed orally with equivalent amounts of tamoxifen.

In the present study, mice with tamoxifen-inducible CreER and the *Lhx2* flox alleles treated with tamoxifen are referred to as *Lhx2*^flox/flox^ while those without the CreER and treated with tamoxifen are labelled as wild type (WT) controls. In the case of standard knockouts, the homozygous knockout pups are referred to as *Lhx2*-/- and the homozygous wild types as WT. Gonads were collected between E11.5 to E15.5 from the *Lhx2-/-* or the *Lhx2*^flox/flox^ strain.

### Allelic genotyping and embryonic sex genotyping

To determine the *Lhx2* genotype and the fetal sex, PCR was done using the Terra™ PCR Direct PCR mix (Takara Bio, Kusatsu, Shiga Japan). Genotyping for *Lhx2* was performed as described earlier (Mangale et al., 2008). The PCR reactions with *Lhx2* primer pairs give a 385 bp band for the WT allele and a 550 bp band for the mutant allele. The sex of the embryos was determined using the primer pairs for the *Jarid* gene as described earlier (Clapcote and Roder, 2005). The primer sequences are given in Supplementary Table 2.

### RNA extraction, cDNA synthesis, and qPCR

Gonads were dissected and the mesonephros was carefully separated. The gonads were stored in RNA later (ThermoFisher Scientific, MA, USA). RNA was extracted using the RNeasy mini kit (Qiagen, Germany) on column-based DNase I (ThermoFisher Scientific) digestion was performed following the manufacturer’s protocol. cDNA was synthesized using High Capacity cDNA synthesis kit (Applied Biosystems, ThermoFisher Scientific, MA, USA) and quantitative PCR was done using Eva Green chemistry (BioRad Laboratories Inc., CA, USA) in the CFX96 Real-Time PCR System (BioRad) as detailed previously (Godbole and Modi, 2010; Mishra et al., 2020a). Each PCR was run for 35 cycles which involved initial denaturation of 95°C for 30 sec, primer annealing at optimized temperature for 30 sec, and extension at 72°C for 30 sec with a final extension step at 72°C for 5 min. PCR amplifications were carried out in duplicates, for all biological replicates. Primer efficiency value for each primer pair was calculated from standard curve analysis of serially diluted cDNA. The primer sequences, their annealing temperatures, and expected product sizes are given in Supplementary Table 2. Data were analyzed using the Pfaffl method as described earlier (Godbole and Modi, 2010).

### RNA Sequencing

The strategy of gonad collection for RNAseq is shown in Supplementary Fig 1. Pregnant females were fed with tamoxifen (as described above) and the pups were collected on E12.5. The gonads were separated from the mesonephros and preserved in RNA later (ThermoFisher Scientific). A minimum of 30 pairs of gonads each from the WT and *Lhx2*^flox/flox^ embryos were pooled and outsourced for RNA sequencing (Sandor Biosciences, Hyderabad, India) using Illumina NextSeq 500 platform. Illumina TruSeq adapters were used; the read length was 76 nucleotides and was paired at both ends. An average of 45 million raw reads was generated per sample. Alignment was done using Tophat-2.0.13 using mm84 sequence as reference (ftp://ftp.ensembl.org/pub/release-84/fasta/mus_musculus/dna/).

### Differential gene expression analysis

Fold change in FPKM values in *Lhx2*^flox/flox^ ovaries was calculated with respect to WT ovaries. Genes with a fold change of 0.5 or less were taken as downregulated and fold change of ≥ 1.5 is upregulated. The differentially expressed genes were used for Gene Set Enrichment Analysis (GSEA) using the WEB-based GEne SeT AnaLysis Toolkit (http://www.webgestalt.org/) at the default settings.

### Histology and Immunohistochemistry

To study histology, mesonephros gonad complex was fixed in 4% paraformaldehyde (Sigma Aldrich) and 5um thick paraffin sections were stained with Hematoxylin-Eosin as detailed earlier (Mishra et al., 2020b). Immunofluorescence was performed as described previously (Godbole et al., 2017; Tiwari et al., 2021). In brief, sections were deparaffinized and epitopes were exposed in antigen retrieval buffer (Tris-EDTA pH-6.0). Blocking was done in 1% Bovine Serum Albumin (MP Biomedicals; Maharashtra, India) and the sections were probed overnight in 1:10 diluted anti-VE-Cadherin antibody (mouse monoclonal 8E5 clone, ELK Biotech, Wuhan China, catalogue # EM1322). Negative controls were incubated either with isotype controls or Phosphate Buffered Saline (PBS) pH 7.2 instead of the primary antibody. The next day, slides were washed thrice in PBS, and detection was done using the Alexa 594 conjugated donkey anti-mouse antibody (Invitrogen, ThermoFisher Scientific, MA, USA Cat # A-21203). The sections were visualized and images were captured using a fluorescence microscope equipped with a sCMOS camera (Leica Microsystems DMi8, Mannheim, Germany). The hematoxylin-eosin stained sections were viewed under a bright-field microscope (Olympus; Tokyo, Japan) and representative areas were imaged using a digital camera (Olympus). From the images, the desired areas were extracted using ImageJ and digitally enhanced. All quantifications were done before image enhancements.

### *In Vitro* Gonad Recombination Assay and estimation of branching pattern

The scheme of the gonad recombination assay is described in Supplementary Fig 2. Breeding pairs of FVB mice constitutively expressing Green fluorescent protein (GFP) were originally obtained from the National Institute of Immunology (NII), New Delhi, India. Mice were bred and maintained at the Experimental Animal Facility of ICMR-NIRRCH. For gonad recombination assays, the pups from FVB-GFP and *Lhx2* heterozygous mothers were collected on E11.5. The GFP mesonephros and the gonads were recombined in agarose blocks. The dimensions of agar mould and the protocol are detailed elsewhere (Capel and Batchvarov, 2008). Briefly, the sterile moulds were poured with sterile 1.5% agar (prepared in DMEM-F12) and allowed to cool. The agar blocks were removed and equilibrated in DMEM-F12 with 10% fetal bovine serum and antibiotic-antimycotic mixture (all from Gibco, ThermoFisher Scientific, MA, USA). The gonads and mesonephros were carefully dissected and the GFP mesonephros (of either sex) was carefully overlayed with the gonad from WT or *Lhx2*-/-ovaries in the ridges of the agar mould. XY gonads from WT pups were used as positive controls. The gonadal assembly was cultured at 37^0^C with 5% CO_2_ and migration of the GFP positive cells was monitored at 24, 48, and 72 hours post culture. Recombinants were imaged on a stereomicroscope equipped with fluorescent imaging (Olympus).

### Quantification of angiogenesis

For quantification of angiogenesis patterns, the angiogenesis analyser macros in ImageJ software (https://imagej.nih.gov/ij/macros/toolsets/AngiogenesisAnalyzer.txt) was used (Carpentier et al., 2020). The desired areas were selected, the background was corrected and the images were subjected to the angiogenesis analyser that quantified various branching parameters. The details of the parameters and their interpretation are provided in (Supplementary Table 3). All the images of the recombinant gonads were blinded for their sex and genotype and then analyzed. Data from 3 independent experiments (n=3 gonads) each from WT XX, WT XY, and *Lhx2*-/-XX at 72h of recombination was analyzed. Mean and standard deviation was computed and statistical analysis was applied.

### Assessment of endothelial cell transcriptome

We used the previously curated dataset of cell-type-specific transcriptome where the genes enriched in the endothelial cells of both the sexes are enlisted (Jameson et al., 2012). This data set is generated out of microarray analysis of flow-sorted *Flk1*-mCherry positive endothelial cells from XX and XY mouse gonads at E11.5, E12.5, and E13.5. The dataset contains 473 genes at E12.5 whose expression is significantly higher in endothelial cells as compared to other cell types (p-value cutoff of 0.05, and a fold change cutoff of 1.5). Information for 445/473 endothelial cell-enriched genes was available in our RNAseq dataset. Their FPKM levels were compared between WT XX vs WT XY vs *Lhx2*^flox/flox^ XX gonads at E12.5. Those genes whose expression in the XY gonads was higher than those of XX were considered male-biased and those with expression lower in XY gonads as compared to XX were considered female-biased (fold change >1.25 or <0.5 respectively).

### Data analysis and presentation

All the data were analyzed statistically using ANOVA and graphs were plotted using GraphPad Prism (Version 8, California). Heatmaps were plotted using Morpheus (https://software.broadinstitute.org/morpheus). The GSEA results were visualized using the R studio version 3.6.2 with ggplot2 packages (RStudio Team, Boston, MA http://www.rstudio.com/) as described previously (Ashary et al., 2020; Colaco et al., 2021). The graphical abstract and experimental schematic in supplymentery figures were created with BioRender.com

## Results

### Expression profiles of *LIM-HD* genes in developing mouse, rat, and human gonads

Fig 1A shows expression profiles of twelve *LIM-HD* genes that are present in the mammalian genome. As evident, *Lhx5* is not expressed in the mouse, rat, and human gonads at the time of sex determination. The others are expressed at varying levels with some being sexually dimorphic while others are equally expressed in both the sexes. In general, most *LIM-HD* genes were expressed in a female dominant manner in the mouse and human gonads, while in rat gonads some of them were male dominant. For example, *Lmx1b* is female dominant in XX mouse and human gonads while it is XY dominant in rat gonads. Some genes were sexually dimorphic only in one species but not others. For example, *Isl1* and *Isl2* are largely female dominant in the human gonads but not in the mouse and rat gonads; *Lhx4* is female dominant only in the rat gonads. Intriguing, irrespective of the species, *Lhx2* was XX dominant during sex determination in all three species. In the mouse, as early as E10.5 *Lhx2* was XX dominant and remained high throughout the window of sex determination. In the rat, *Lhx2* became sexually dimorphic slightly later (just after the onset of sex determination, E15), while in the humans, it was XX dominant right from 7 weeks and remained high in the female gonads throughout (until 11 weeks).

**Fig 1:**
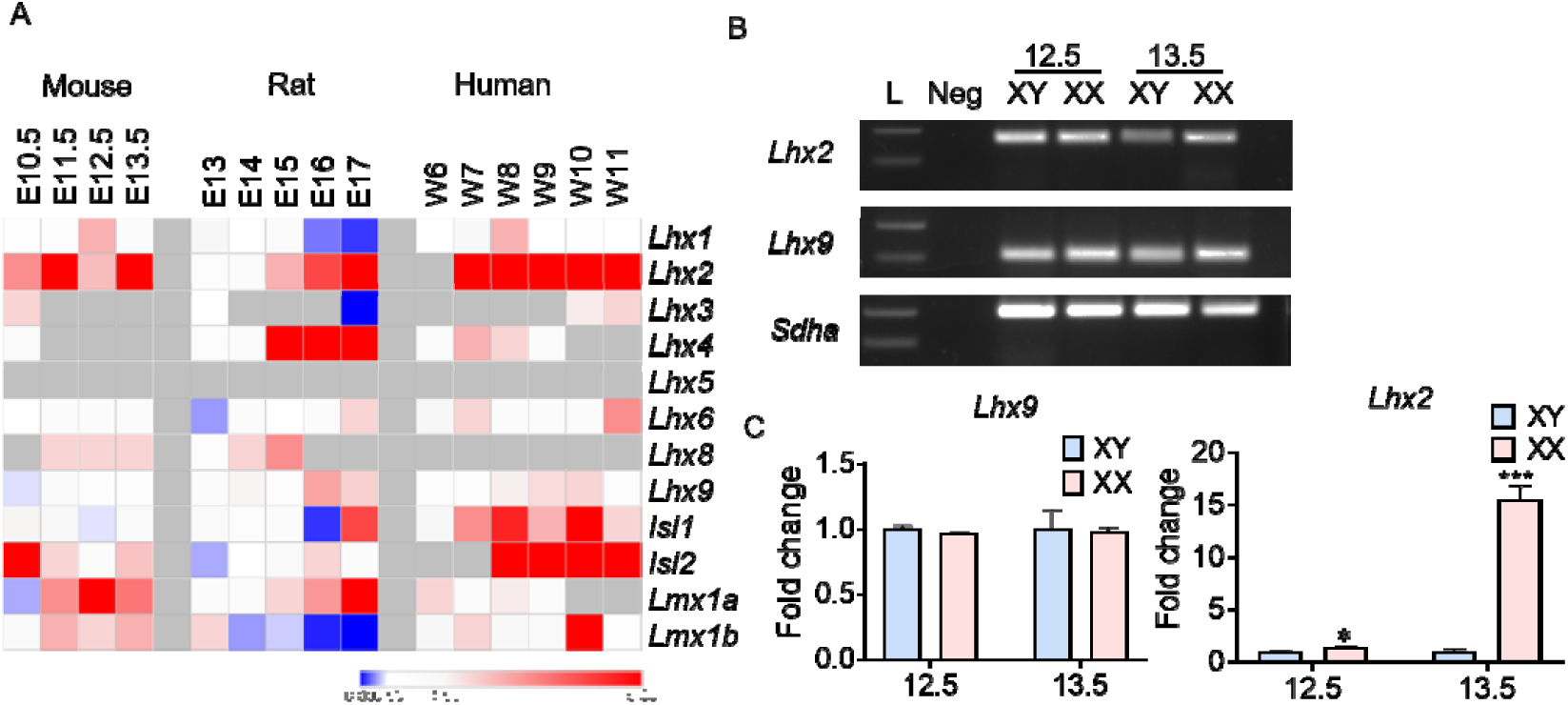
Expression profile of LIM homeobox genes in mammalian developing gonads. (A) Heatmap demonstrating the ratio of FPKM values (XX/XY) for LIM homeobox. genes (*Lhx*1-6, *Lhx8-9, Isl*1*-2, Lmx1a-lb)* in gonads. Bach row is the gene and each column is the time point in embryonic (E) development m days for the mouse and rat and weeks (W) in humans. FPKM values were extracted from RNA-seq dataset (Cardoso et aL,2019). Boxed in red shades indicate female dominance while those in blue shades indicate male dominance. White boxes indicate the levels are not sexually dimorphic (XX/XY ratio >0.5 or <l.5). Grey are missing values. (B) Gel image showing expression of *Lhx2* and *Lhx9* by RT-PCR in developing gonads at El2.5 and E13.5. *Sdha* is housekeeping gene and L is molecular marker of l00bp. Neg lane is loaded with no DNA template control (C) Quantification of *Lhx2* and *Lhx9* mRNA levels by qPCR in XX and XY gonads at E12.5 8 and E13.5 (X axis). values were normalized to *Sdha* and the data is expressed in fold change of the mean value obtained for XY gonads. Values On. Y axis are mean + SD (n=3 biological replicates/time point). *p<0.05, ***p<0.00 l

We performed RT-PCR for *Lhx2* and *Lhx9* in the XX and XY mouse gonads at E12.5 and E13.5. As expected, both these genes were expressed in XX and XY gonads (Fig 1B). Quantitative analysis showed that at both E12.5 & E13.5, *Lhx2* was significantly higher in the XX as compared to XY mouse gonads (Fig 1C). As predicted from the RNAseq data, *Lhx9* was not sexually dimorphic at E12.5 and E13.5 (Fig 1C).

### Sexual dimorphic expression of *Lhx2* is evolutionarily conserved

Since *Lhx2* was sexually dimorphic with higher expression in XX when compare to XY gonads in mice, rats, and humans, we sought to compare its expression in the gonads of other species with environmental or genetic (XX/XY and ZW/ZZ) modes of sex determination during gonad development. The red-eared slider turtle (*Trachemys scripta elegans*) has a temperature-dependent sex determination mechanism (Czerwinski et al., 2016). In this species, eggs incubated at female promoting temperature (FPT, 31°C) had higher expression of *Lhx2* as compared to those incubated at male promoting temperature (MPT, 26°C) from the onset of sex determination. One of the species of lizards (*Pogona vitticeps)* uses the genetic and temperature-dependent sex determination mechanisms, and higher temperature >32^°^C can override the genetic mode of sex determination (Whiteley et al., 2021). In this species, throughout sex determination, *Lhx2* was 3-6 folds higher in the eggs incubated at FPT as compared to MPT irrespective of their genetic sex (Fig 2). Chicken has ZW/ZZ mechanism of sex determination and the female is heterogametic (Capel, 2017). In the chicken (Gallus gallus), *Lhx2* expression was 2-4 folds higher in the gonads of ZW (female) embryos as compared to ZZ (male) during the period of sex determination (Fig 2). Opossum (*Monodelphis domestica*) has XX/XY mechanism of sex determination and the SRY gene has first evolved (Katsura et al., 2018). In this species too, irrespective of the time point, XX gonads had 7-20 folds higher expression of *Lhx2* as compared to XY gonads (Fig 2). In mice and humans, both of which utilize the XX/XY mode of sex determination, *Lhx2* was female dominant in both species. In mice, *Lhx2* was at least 2-4 folds higher in XX gonads as compared to XY. However, in humans, *LHX2* expression was 15-18 folds higher in the XX gonads between W9-12.

**Fig 2:**
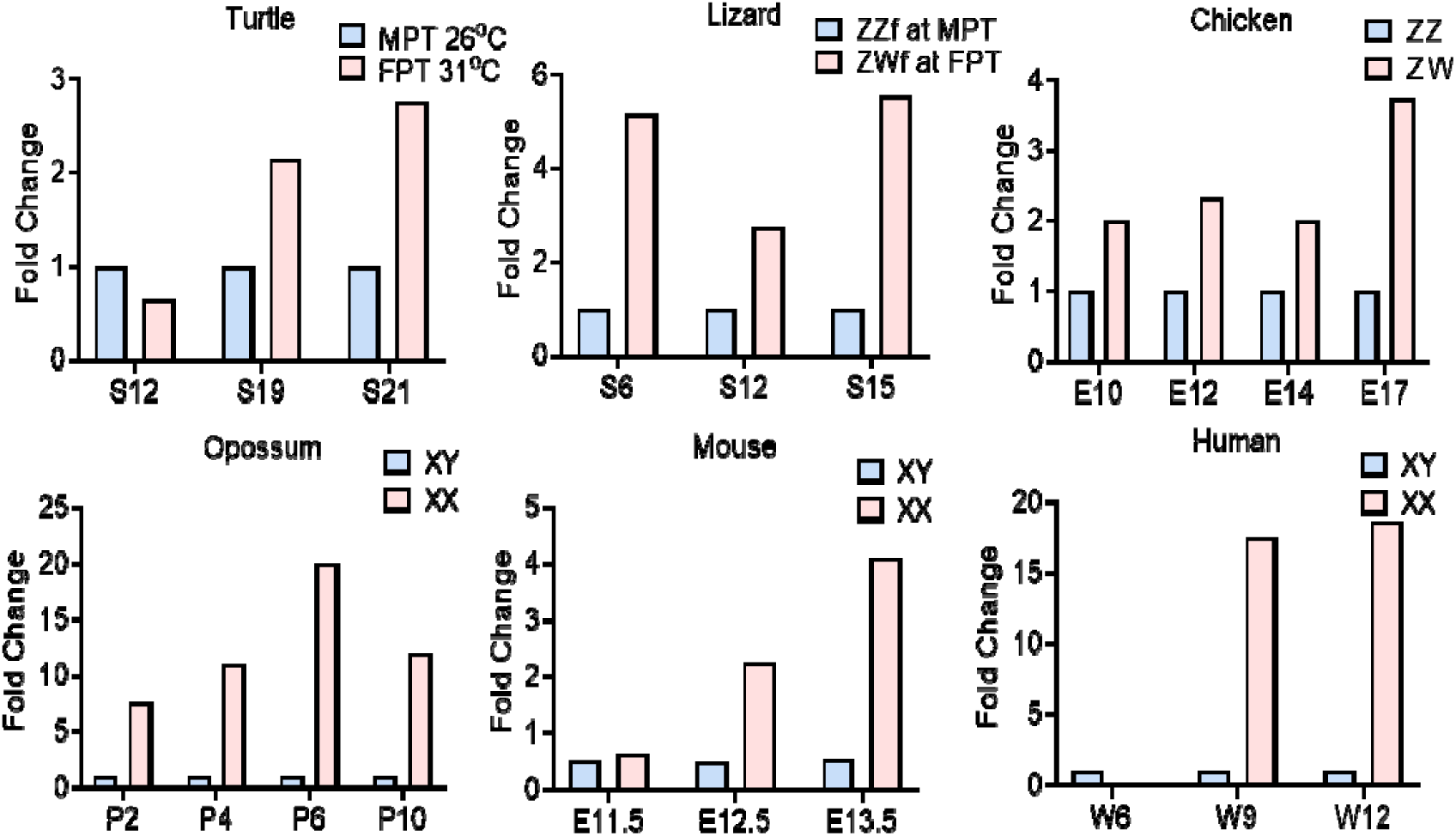
Sexually dimorphic expression of *Lhx2* in developing gonads of species with different mechanisms of sex determination. Bulk RNA seq datasets of turtle (SRP079664; Czerwinski et al., 2016). lizard (PRJNA699086; Whiteley et al., 2021). chicken (E-MTAB-6769; Cardoso-Moreira et at., 2019), opossum (E-MTAB-6833; Cardoso-Moreira et al., 2019), mouse (SRP076584; Zhao et at., 2018) and human (GSE116278; Lecluze et al., 2020) were extracted and analysed. Expression values in each species were converted to fold change with respect to XY values at one and plotted on Y axis. X axis shows the time point of embryonic development.

### *Lhx2* is dispensable for gonad development and somatic cell sex determination

Gonads from the *Lhx2*-/-embryos were well developed with distinct mesonephros and gonad proper. Histologically, *Lhx2-/-* XX gonads were indistinguishable from that of the WT controls at both E13.5 and E15.5 (Fig 3A). Although in some instances, the XX gonads of *Lhx2-/-* embryos at E15.5 appeared rounded and attached to the mesonephros (data not shown). There was no evidence of tubular organization in the *Lhx2*-/-XX gonads like that seen for the XY gonads at E13.5 and E15.5 (Fig 3A)

**Fig 3:**
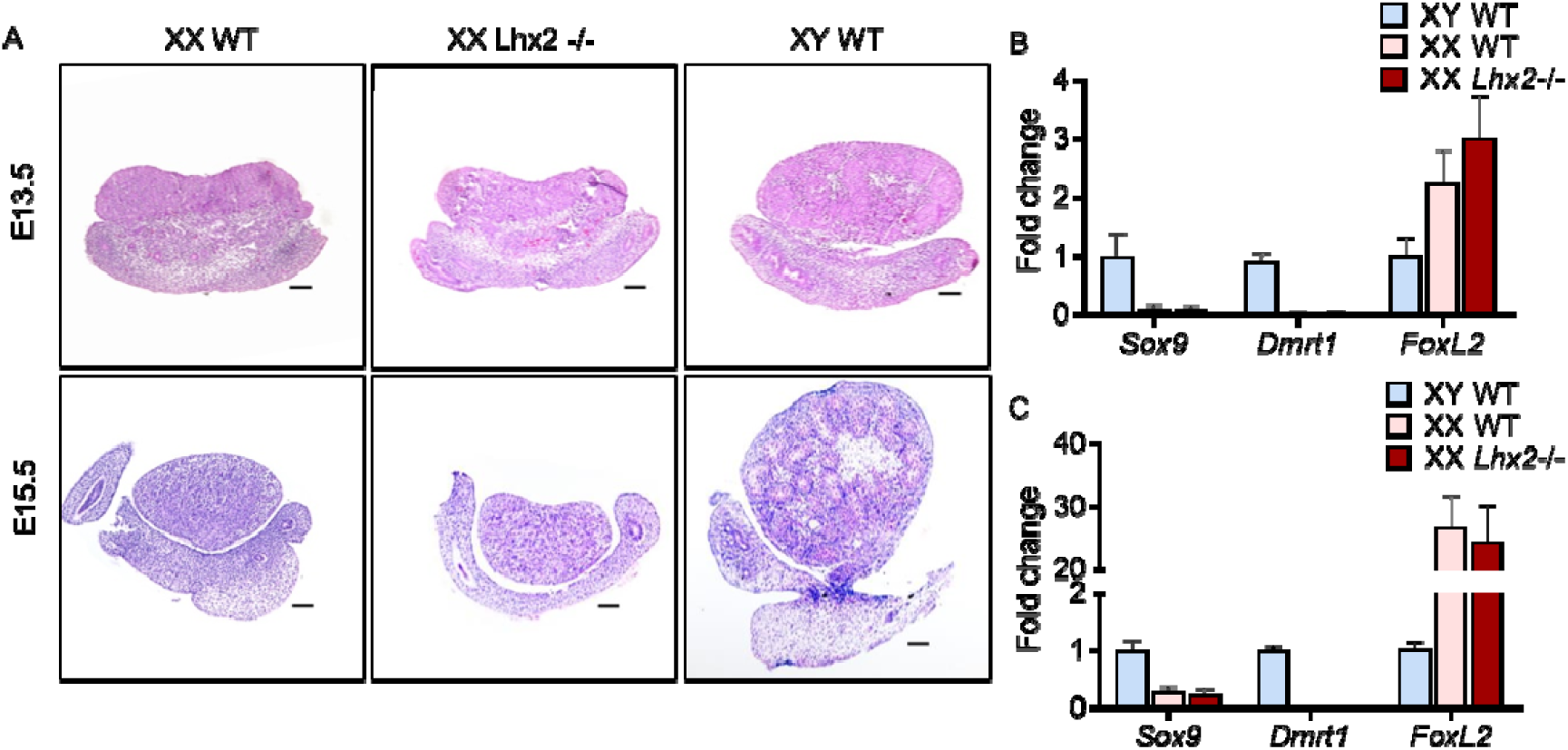
Somatic sex is not reversed in absence of *Lhx2*. (A) Histology of wild type (WT) XY, XX and XX *Lhx2-/-* gonads at E13.5 and. El5.5. Scale bar is of 50μm. (B) Quantification of mRNA levels of *Sox9* and *Dmrt1* (Sertoli cell marker) and *Fox12* (Granulosa cell marker) by qPCR. in the WT XY, XX and *Lhx2-/-* XX gonads at El3.5. (C) Quantification of mRNA levels of *Sox9* and *Dmrt1* (Sertoli cell marker) and *Foxl2* (Granulosa cell marker) by qPCR in WT XY, XX *and Lhx2-/-* XX gonads at El5.5. Values for *Sdha* were used for normalization. Data is presented as a mean+SD of fold change of the WT XY gonads for 6 biological replicates/time point/sex/genotype.

We analyzed the levels of key genes involved in sex determination in the developing *Lhx2*^flox/flox^ gonads at E12.5 and observed that most of these genes (except *Sf1*) were expressed similarly at identical levels in *Lhx2*^flox/flox^ gonads as compared to their WT counterparts (Supplementary Fig 1). We also analyzed the mRNA levels of male somatic cell-specific genes *Dmrt1* and *Sox9* at E13.5 and E15.5 by qRT-PCR. As expected, both these genes were sexually dimorphic with higher expression in XY gonads as compared to XX gonads at E13.5 and E15.5. The levels of these genes did not significantly differ in ovaries of WT and *Lhx2-/-* embryos (Fig 3B and 3C respectively). As expected, the mRNA level of *Foxl2* (female somatic cell marker) was higher in the WT XX gonads as compared to the WT XY gonads. The level of *Foxl2* was identical in the WT and *Lhx2-/-* XX gonads at E13.5 and E15.5 (Fig 3B and 3C respectively).

### Identification of genes regulated by *Lhx2* in the XX gonads

To address the roles of *Lhx2* in the developing XX gonads, we carried out RNA seq of WT XX, WT XY, and *Lhx2*^flox/flox^ XX gonads at E12.5. A total of 4076 genes were found to be differentially expressed in the *Lhx2*^flox/flox^ XX gonads as compared to WT. Amongst these, 2566 genes were upregulated while 1510 were downregulated in the *Lhx2*^flox/flox^ ovaries as compared to the wild type ovaries (Supplementary Table 4). Of the down-regulated genes, 33 genes were exclusively expressed in the WT XX gonads but not in the *Lhx2*^flox/flox^ XX gonads (Supplementary Table 4). *Isg15* was the most upregulated gene, while *Lgals3* was the most down-regulated gene in the *Lhx2*^flox/flox^ ovaries (Supplementary Table 4).

Differentially expressed genes were analyzed using GSEA (Fig 4). The results revealed that biological processes like organ or tissue immune responses, protein-DNA complex subunit organization, and hepatobiliary development were overrepresented, while synaptic transmission, sensory perception to taste, cilium development were under-represented (Fig 4A). Amongst the molecular functions, semaphorin receptor binding, double-stranded RNA binding, and beta-catenin binding were overrepresented while peptidase activity, structural constituents of the ribosome, electron transfer activity, and antioxidant activity were significantly underrepresented (Fig 4B). The panther pathways associated with vascular functions such as angiogenesis, PDGF and WNT signalling pathway, insulin/IGF1 protein kinase B signalling cascade, and cadherin signalling pathways were significantly overrepresented (Fig 4C). The pathway associated with DNA replication were underrepresented (Fig 4C). The pathway enrichment plots and expression profiles of a subset of genes contributing significantly to the enrichment of angiogenesis and PDGF signalling pathways are shown in Fig 4D. The enrichment plots of other selected biological processes and pathways are shown in Supplementary Fig 3.

**Fig 4:**
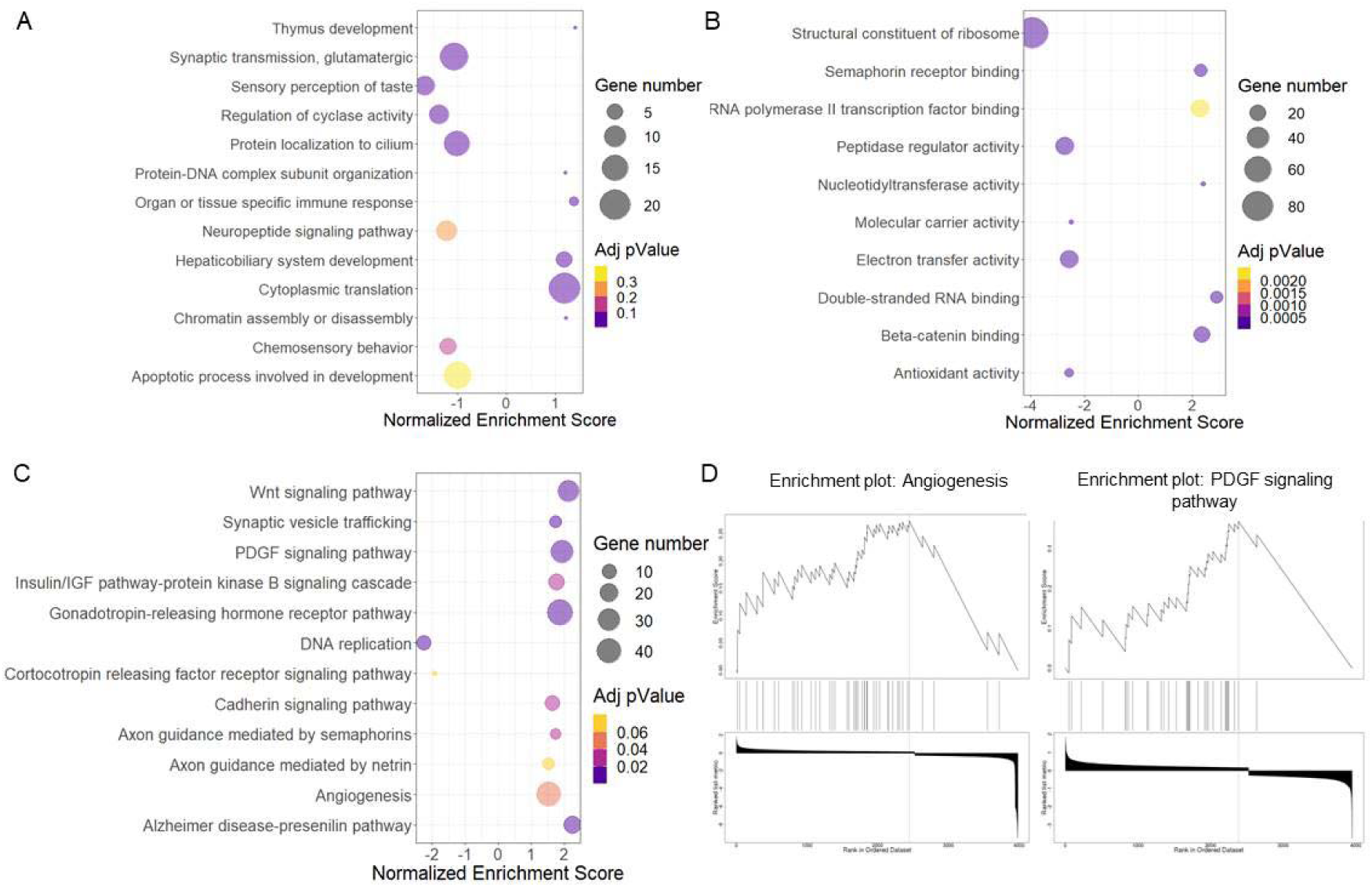
Gene Set Enrichment Analysis (GSEA) for the differentially expressed genes in *Lhx2-/-* XX gonads. The differentially expressed genes between wild type and *Lhx2*^flox/flox^ XX gonads at El2.5 were subjected to GSEA using the WEB-based GEne SeT AnaLysis Toolkit (http://www.webgestalt.org/). (A) The enriched biological processes. (B) molecular Functions and (C) panther pathways are shown as bubble plots. X axis is normalized enrichment score where the highly enriched processes are skewed on the right. The numbers of genes and the adjusted p values for each process is shown. (D) Enrichment plots for the angiogenesis pathway and PDGF signaling pathway. Y axis represents enrichment score which shows the degree of overrepresentation of gene sets and vertical bars shows where the member of the gene set appeared in ranked list The bottom part of each graph shows ranking matrix that measures the gene’s correlation with their phenotype.

We performed immunofluorescence for VE-Cadherin and observed a higher number of endothelial cells in the WT XY gonads as compared to XX WT gonads (Fig 5C). However, many VE-Cadherin-positive cells were detected in the ovarian stroma of E13.5 in the *Lhx2-/-* XX gonads (Fig 5C). The negative control sections incubated with an isotype IgG did not show any reactivity (Fig 5C inset).

**Fig 5:**
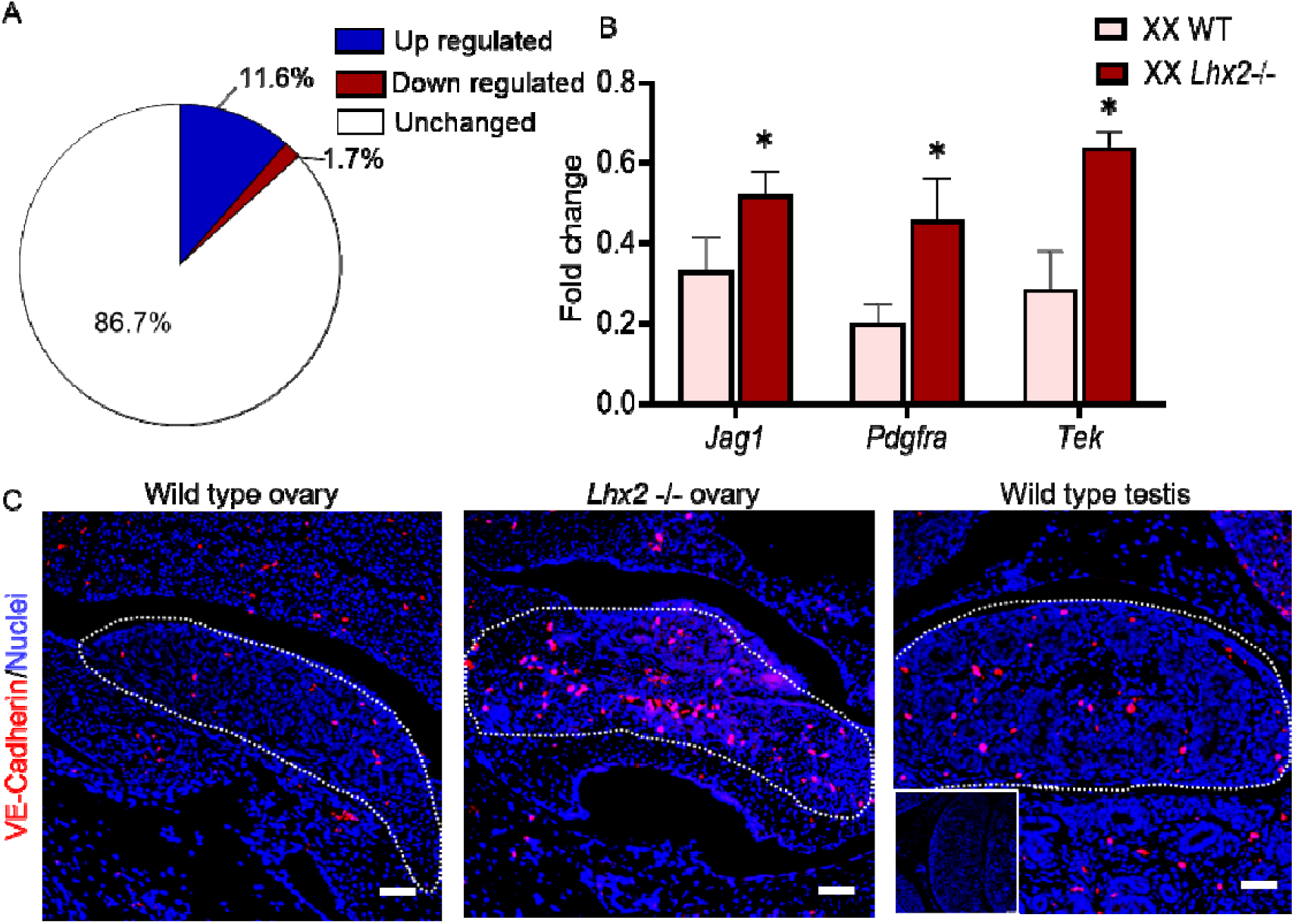
Endothelial genes are enriched in the ovaries lacking *Lhx2*. **(A)** Percentage of endothelial cell specific genes altered in the ovaries of *Lhx2*^flox/flox^ embryos as compared to wild type (WT) XX at El2.5. **(B)** qPCR for endothelial enriched genes at E13.5 in WT and *Lhx2-/-* XX gonads. Data is presented as fold change where mean values of was taken as l. Values are mean+SD of three biological replicates. ** p<0.01, *** p<0.001 **(C)** Immunofluorescence for VE-Cadherin (red channel) in wild type wild type and *Lhx2-/-* ovaries and wild type testis at E13.5. Nuclei are stained with DAPI (blue channel). Negative control is shown in inset Scale bar is 50 μm. The dotted area marks the gonad.

### *Lhx2* is expressed in the germ cells and not in endothelial cells of XX gonads at E13.5

After filtering for low-quality cells (cells with less than 200 genes and those expressed in less than 3 cells were removed). we obtained a total of 4,842 cells at E13.5. After performing tSNE, based on gene expression diversity we could identify a total of 17 independent clusters (Fig 6A). Of these *Lhx2* was detected in the cells in three clusters (Fig 6B). However, none of these clusters expressed endothelium-specific genes *Kdr* and *Flt1*. Instead, the endothelium-specific genes were detected in a separate cluster of the tSNE plot (Fig 6B). The cluster of cells that expressed *Lhx2*, expressed the germ cell-specific marker *Ddx4* (Fig 6B) and *Dazl* (Fig 6F) confirming that *Lhx2* is expressed by the germ cells. Other cell types (if any) barely expressed *Lhx2* and these were not endothelial cells (not shown).

**Fig 6:**
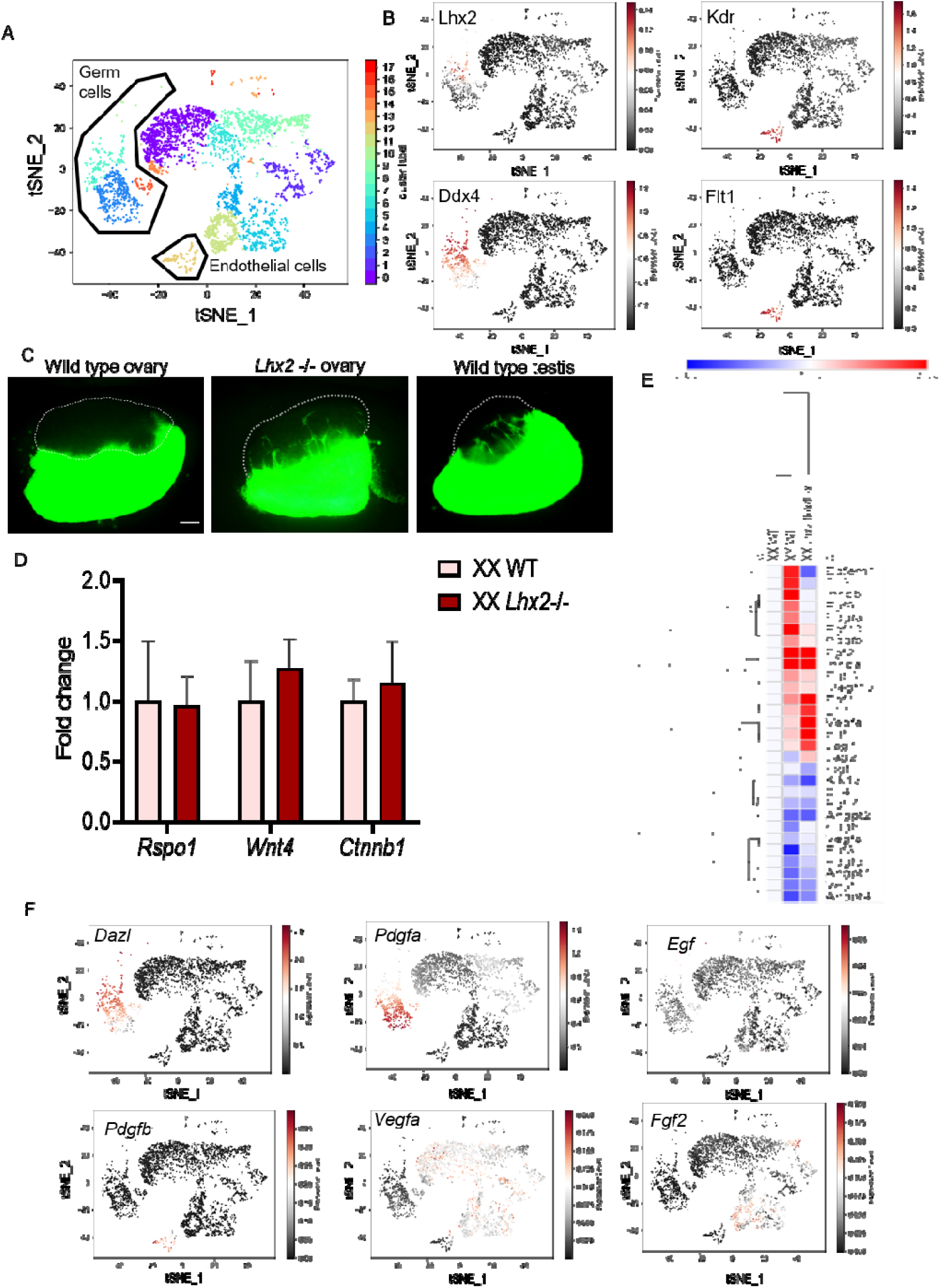
*Lhx2* is expressed in the germ cells and its loss leads to excessive mesonephric endothelial cell migration in a cell non-autonomous manner. (A) tSNE plot of single cell RNA seq dataset (GSE128553) of mouse ovaries at E13.5 showing clustering of different cell types. (B) Imputed expression of *Lhx2* and marker genes in the tSNE plot. The germ cell marker is *Ddx4* end endothelial cell markers are *Kdr* and *Flt*. (C) Gonad recombination assay where mesonephros from wild type (WT) XX or XY animals constitutively expressing GFP were overlaid with gonads from WT XY, WT XX and XX Lhx2-/-embryos. The co-cultures were monitored for mesonephric cell migration, at 48h. The region of the gonad is outlined. The scale bar is 100 μm (D) qPCR of *Rspol, Wnt4* and *Ctnnb1* in *Lhx2-/-* XX gonads at E13.5. Data is presented as fold change where mean values of WT was taken as 1. Values are mean+SD of three biological replicates. (D) Heatmap of secretory factors involved in endothelial cell migration in the WT XX, WT XY and *Lhx2*^flox/flox^ XX gonads at E12.5. The relative scale is shown. (E) Imputed expression of genes involved in endothelial cell migration projected on the tSNE plot shown in A. *DazL* is a germ cell marker.

### Loss of *Lhx2* leads to excessive endothelial cell migration from the mesonephros in a cell non-autonomous manner

Since *Lhx2* is expressed by the germ cells and not the endothelial cells, we hypothesized that the large numbers of endothelial cells in the *Lhx2-/-* XX gonads could be due to their excessive migration in a cell non-autonomous manner. To test, if the angiogenesis observed in the *Lhx2-/-* XX gonads is cell-autonomously regulated, we performed a gonad recombination assay. As expected, irrespective of the genetic sex (XX/XY) of the mesonephros, by 48 hrs of incubation, GFP-positive cells were seen streaming into the XY gonads, but not in the XX gonads (Fig 6C). In the experiments where *Lhx2*-/-XX gonads were co-cultured with GFP expressing mesonephros, several GFP-positive cells could be detected in the gonads at various regions (Fig 6C). In the WT XX gonads, GFP-positive cells could not be detected even at 72h of co-incubation (not shown) indicating the specificity of the assays.

Previous studies have shown that mice knockout for *Wnt4, Rspo1*, or *Ctnnb1* have ectopic blood vessel development in XX gonads (Chassot et al., 2014). To test if the elevated endothelial cells migration in the *Lhx2-/-* XX gonads is due to loss of *Wnt4, Rspo1*, or *Ctnnb1*, we performed qPCR for these three genes in XX gonads of WT and *Lhx2-/-* embryos at E13.5 (Fig 6D). The results revealed that the levels of *Wnt4, Rspo1*, or *Ctnnb1* were comparable in both groups. We also analyzed the level of *Lhx2* from the RNAseq dataset of *Wnt4* KO (Naillat et al., 2015). The results revealed that the mRNA levels of *Lhx2* are identical in *Wnt4* KO and WT embryos (Supplementary Fig 3F).

We next studied the mRNA levels of 29 secretory factors that are known to aid in endothelial cell migration and angiogenesis in WT XY gonads. As evident from Fig.6E, the mRNA levels of most of the angiogenic factors involved in endothelial cell migration are elevated in WT XY gonads and were also elevated in *Lhx2*^flox/flox^ ovaries and those that are down-regulated in WT XY gonads were also down-regulated in the *Lhx2*^flox/flox^ ovaries at E12.5 (Fig 6E).

To test if any of these factors are secreted by the germ cells we analyzed their expression in the single-cell RNAseq dataset. The tSNE plots (Fig 6F) of the selected genes revealed that *Pdgfa* is abundantly transcribed by most germ cells, a small subset of germ cells also transcribed *Egf*. The others were expressed by the supporting cells including the endothelial cells (Fig 6F).

### Male-like behavior of the endothelial cells in the XX gonads in absence of *Lhx2*

To characterize the endothelial cells in the XX gonads in absence of *Lhx2*, we analyzed the expression profiles of 445 genes that are known to be enriched in the endothelial cells (Supplementary Table 5). The results revealed that 107/445 genes were male-biased (>1.25 folds in the XY gonads with respect to XX gonads) while 52/445 genes were female-biased (<0.5 folds in the XY gonads with respect to XX gonads) at E12.5. Interestingly, the expression of 62/107 (∼58%) male-biased endothelial cells was elevated in the *Lhx2*^flox/flox^ ovaries (Fig 7A, Supplementary Table 5).

**Fig 7:**
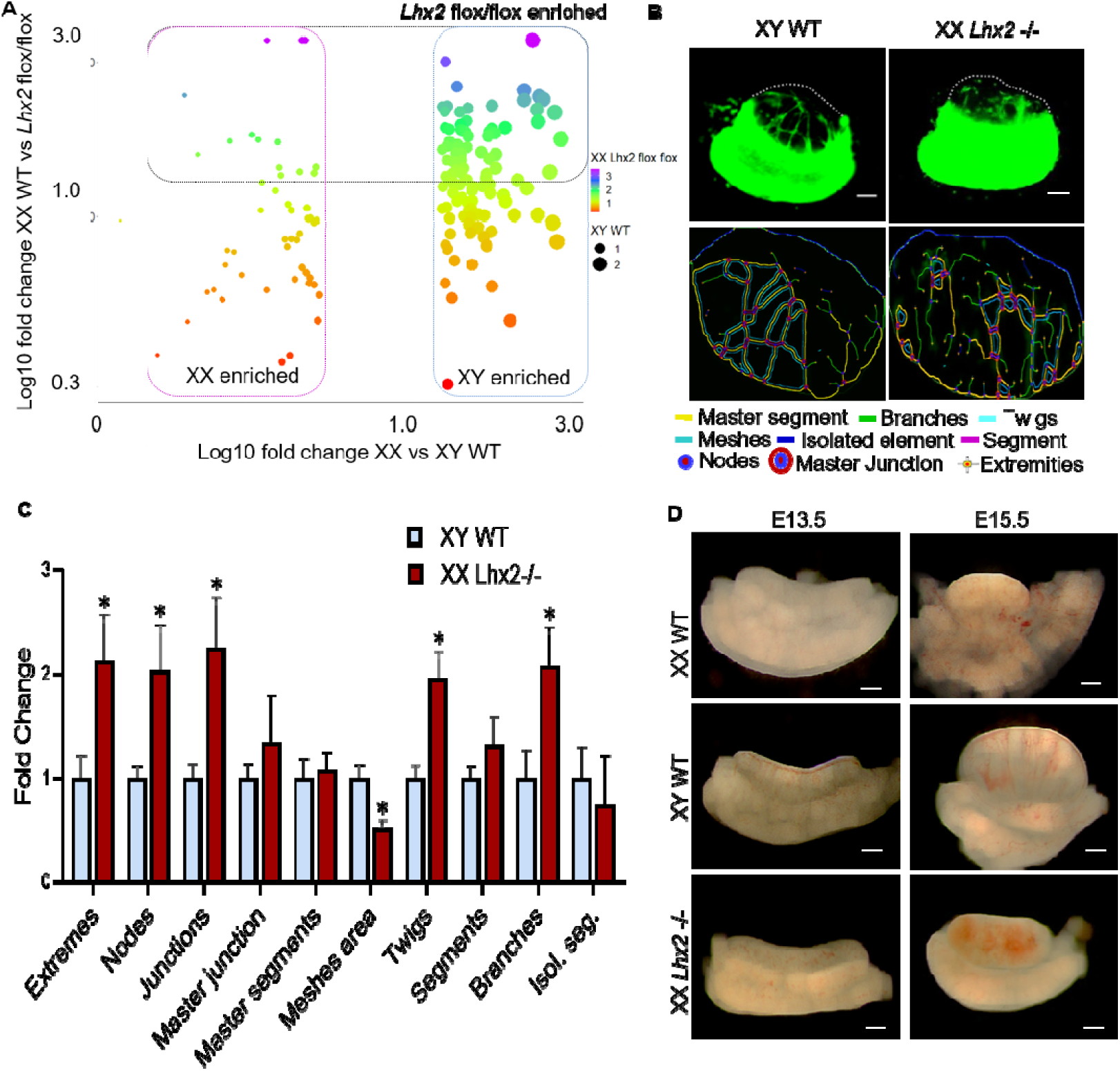
Masculinization of the endothelial cells in the XX gonads In absence of *Lhx2*. **(A)** Bubble plot showing expression levels of endothelial enriched genes (al E12.5) in wild type (WT) XY and in XX *Lhx2*^flox/flox^ gonads normalized to the values observed in WT XX gonads. **(B)** Gonad recombination assay at 72h of co-culture. XY WT, and XX *Lhx2*-/-gonads were used for gonadal recombination assay and observed after 72h of co-culture. Recombinant gonads were analyzed using ImageJ software with angiogenesis analyzer plugin. The different patterns of branching are shown. Scale bar is of 100 μm **(C)** Graph were plotted using fold change of different parameters of angiogenesis. Values are mean-SD for 3 biological replicates and * indicates p <0.05. **(D)** Brightfield images of XX WT, XY WT and XX *Lhx2*-/-gonads at E13.5 and E15.5. Scale bar b of 100μm.

To test if the endothelial cells in the *Lhx2-/-* XX gonads are capable of organizing into a network like that in the testis, we compared the branching patterns of the GFP positive cells in the gonad recombination assay of WT XY and *Lhx2-/-* XX gonads at 72h of co-culture. As expected, WT testis had distinct master segments, branches, and twigs with an intricate meshwork (Fig 7B). Almost a similar meshwork was also seen in *Lhx2*-/-ovaries (Fig 7B). Supplementary Fig 4 gives the representative images for the nodes, extremities, and mesh pattern in the WT testis and *Lhx2-/-* ovaries.

We quantified the pattern of branching in both the conditions (n=3 each) and observed that the numbers of extremes, nodes, junctions, twigs, and branches were significantly higher in the *Lhx2-/-* ovaries as compared to WT testis (Fig 7C). However, the number of segments, master segments, isolated segments, and master junctions were comparable in WT testis and *Lhx2*-/-ovaries (Fig 7C). The mesh area was significantly lower in *Lhx2-/-* XX gonads as compared to WT testis (Fig 7C).

We next asked if the endothelial cells that migrated in the *Lhx2-/-* ovaries form a coelomic blood vessel (Fig 7D). In WT testis (n-10) at E13.5, a distinct coelomic blood vessel was detected which was not observed in WT XX gonads (Fig 7D). Although some vascularization was observed in *Lhx2-/-* XX gonads (n=10), there was no distinct coelomic blood vessel akin to WT testis. We also analyzed the gonads at E15.5 where a distinct vascular network was seen in the XY gonads (n=10) but this was not evident in the XX gonads (Fig 7D). In most *Lhx2*-/-XX embryos, the gonads (n=15) had hemorrhagic areas, a distinct coelomic blood vessel was not observed (Fig 7D).

## Discussion

In the present study, we demonstrate that multiple *LIM-HD* genes are expressed in the developing gonads during the window of sex determination. Amongst these, *Lhx2* is predominantly expressed in the developing ovaries in most species irrespective of the mode of sex determination. In the mouse ovaries, *Lhx2* is exclusively expressed in the germ cells and its loss leads to an increase in endothelial cell migration in a cell-non autonomous manner.

*LIM-HD* genes are transcription factors that play a fundamental role in cell fate determination and patterning of somatic tissues. In the context of gonad development, three *LIM-HD* genes are known to play crucial roles. *Lhx1* has a role in Mullerian duct development; *Lhx9* is required for the development of bipotential gonad and *Lhx8* is essential for oocyte survival in the developing ovaries (Singh et al., 2021). However, along with these three, other members of the *LIM-HD* gene family are also dynamically expressed in the developing mouse gonads with many having a sexually dimorphic expression during the window of sex determination (Singh et al., 2021). Extending these findings, herein we show that except for *Lhx5*, mRNA for most members of *LIM-HD* genes are detected in the mouse, rat, and human gonads during the window of sex determination. Interestingly, many of these, have female biased expression during sex determination. These include *Lhx2, Lhx4, Ils2, Lmx1a*, and *Lmx1b*; all of which are female biased in human gonads and also in one of the two rodent species. These results imply that *LIM-HD* genes might have a role in ovarian development. Indeed, *LIM-HD* binding elements are enriched in several meiosis-specific genes in the germ cells of developing ovaries (Li et al., 2013).

In the vertebrate gonads, sex-determining mechanisms are highly divergent where environmental cues and genes, individually or in combination operate to determine the fate of the bipotential gonads (Bhattacharya and Modi, 2021; Capel, 2017). Our analysis revealed that in the mouse, rat, and humans which have a genetic (XX/XY) mode of sex determination, *Lhx2* is expressed in a sexually dimorphic manner with higher expression in females as compared to male gonads. To determine if the sexually dimorphic expression of *Lhx2* is influenced by the mode of sex determination, we compared its mRNA levels in developing gonads of species with environmental (mainly temperature) and genetic (ZZ/ZW or XX/XY) mode of sex determination. The red-eared slider turtle has a temperature-dependent sex determination mechanism (Czerwinski et al., 2016) while *Pogona vitticeps* (a species of lizard) has both genetic and a temperature-dependent sex determination mechanism where temperature can override the genetic mode of sex determination (Whiteley et al., 2021). In both these species, levels of *Lhx2* were higher in the developing ovaries as compared to testis. In the chicken which has exclusively ZW/ZZ mechanism of sex determination (Capel, 2017; Chue and Smith, 2011), *Lhx2* mRNA was female dominant. The opossum is a marsupial (metatherian) species where the SRY gene has first evolved and has become operative during evolution (Capel, 2017; Katsura et al., 2018). In this species too, *Lhx2* was over abundant in the female gonads as compared to males during the period of sex determination. Thus, our analysis indicates that female dominating expression of *Lhx2* in the gonads is evolutionarily conserved and independent of the different mechanisms of sex determination. A very limited set of genes are known to be expressed in a sexually dimorphic manner in an evolutionarily conserved fashion during the period of sex determination and gonad differentiation (Bhattacharya and Modi, 2021; Capel, 2017). Our study adds *Lhx2* as another gene whose expression is evolutionarily female biased. Based on its conserved expression profiles in the gonads across species, we hypothesized that *Lhx2* might have a key role in ovarian development.

There are two mammalian orthologs of the drosophila apterous gene, *Lhx2*, and *Lhx9*, both of which have high sequence homology and have overlapping patterns of expression in the developing nervous system (Bulchand et al., 2003; Chou and Tole, 2019). *Lhx9* has an indispensable role in gonad development as mice lacking *Lhx9* do develop the genital ridges, but the somatic cells fail to proliferate and the gonad is arrested at the bipotential stage (Birk et al., 2000). However unlike *Lhx9, Lhx2* appears to be dispensable for gonad development as in *Lhx2*-/-embryos, the gonads developed and differentiated correctly into ovaries. At E13.5 and E15.5, the ovaries were histologically indistinguishable from the WT counterparts. Together this data indicated that, unlike *Lhx9* which is essential at the early steps of genital ridges, *Lhx2* is not involved in bipotential gonad development.

Sex determination in the developing gonads involves the differentiation of somatic cells into either Sertoli cells or pre-granulosa cells. A plethora of molecular players in a mutually opposing manner regulates the fate of somatic cells in the developing gonads. There are pro-male factors that suppress the expression of pro-female genes allowing Sertoli cell specification; in parallel, there are pro-female factors that suppress the expression of pro-testis factors allowing the specification of the pre-granulosa cells (Capel, 2017; Ortega et al., 2019; Rotgers et al., 2018; Ungewitter and Yao, 2013; Yildirim et al., 2020). Since the expression of *Lhx2* was female biased, we hypothesized that it might have a role in female sex determination. However, contrary to our expectation, at E12.5, E13.5 and E15.5, *Lhx2* knockout ovaries appropriately expressed the pre-granulosa cell markers, *Foxl2, Wnt4, Rspo1*, and *Ctnnb1*. Also, there was no evidence of sex reversal as the levels of the Sertoli cell markers; *Sox9* and *Dmrt1* were identical between the WT and mutant XX gonads while their levels were significantly high in the WT XY gonads. These results imply that although *Lhx2* is expressed in a female dominant manner, is not involved in somatic cell sex determination (or pre-granulosa cell specification) in the developing gonads.

To determine the roles of *Lhx2*, we next carried out RNAseq of the WT and *Lhx2*^flox/flox^ ovaries at E12.5 and identified a large number of genes that were differentially expressed. GSEA revealed that the biological processes associated with synaptic transmission, sensory perception, and protein localization to cilium were enriched in *Lhx2*^flox/flox^ ovaries. In the brain, *Lhx2* is involved in the development and patterning of the neurons involved in sensory perception (Pal et al., 2021; Zembrzycki et al., 2015). In hypothalamic tanycytes, *Lhx2* regulates the numbers of motile cilia by controlling the expression of *FoxJ1* (Salvatierra et al., 2014). One of the enriched biological processes in *Lhx2*^flox/flox^ ovaries was that of hepatobiliary development. Interestingly, *Lhx2* regulates the differentiation and activation of hepatic stellate cells, and loss of *Lhx2* causes hepatic cell fibrosis (Miyoshi et al., 2019; Wandzioch et al., 2004). Amongst the enriched molecular function in the *Lhx2* knockout ovaries was that of β-Catenin binding. In the developing brain (Hsu et al., 2015; Shetty et al., 2013), and nasopharyngeal carcinoma (Liang et al., 2019) *Lhx2* cooperates with β-Catenin to regulate the expression of its downstream genes. These results imply that *Lhx2* has a role in diverse biological processes in multiple tissues, some of which also overlap in the developing ovaries. It appears that some of the functions of *Lhx2* may be conserved and these commonalities could aid in dissecting its downstream targets or mechanisms that may also be shared between these systems.

Amongst the overrepresented pathways enriched in *Lhx2*^flox/flox^ ovaries was that of angiogenesis. In addition, VEGF and PDGF signaling pathways, EGF receptors, and Wnt signaling pathways that are known to aid in vascularization (Gowdak and Krieger, 2018; Gu et al., 2021; Omorphos et al., 2021) were also enriched in the *Lhx2* knockout XX gonads. These observations prompted us to ask if angiogenesis was altered in the absence of *Lhx2* in the developing ovaries. Towards this, we first tested if angiogenesis has occurred in the developing XX gonads in absence of *Lhx2*. The results revealed that almost 13% of the endothelial cell-enriched genes were differentially expressed and most were upregulated upon loss of *Lhx2*, the mRNA of endothelial cell-enriched genes *Jag1* and *Pdgfr*α were upregulated, the transcription factor *Tek* which is specific to endothelial cells was increased by almost two folds in *Lhx2-/-* ovaries as compared to WT controls. Also, there were high numbers of VE-Cadherin positive endothelial cells observed in *Lhx2-/-* ovaries. These results imply that loss of *Lhx2* leads to increased numbers of endothelial cells in the XX gonads. In breast cancer, overexpression of LHX2 is reported to promote tumor growth by controlling vascularization (Kuzmanov et al., 2014). Thus, the requirements of LHX2 in control of vascularization and/or angiogenesis may be context dependent.

We next determined the cell types that expressed *Lhx2* in the developing mouse ovaries. Based on the phenotypes observed, we suspected that *Lhx2* might be expressed by the endothelial cells and its loss might activate their migration. However, single-cell RNAseq analysis revealed that *Lhx2* is exclusively expressed in the germ cells of the developing ovaries with no expression in the endothelial cells. This prompted us to hypothesize that LHX2 in a cell non-autonomous manner must regulate endothelial cell migration. In the developing testis, there is a breakdown of mesonephric vascular plexus and endothelial cells from the mesonephros migrate into the gonads leading to the establishment of angiogenesis and the formation of a coelomic blood vessel. In contrast, the mesonephric vascular plexus remains intact and there is no migration of endothelial cells into the developing ovaries until E18.5 (Coveney et al., 2008; Gu et al., 2021; Kumar and DeFalco, 2018). Thus to test if the loss of *Lhx2* in the germ cells causes endothelial cell migration, we carried out gonad recombination assays where mesonephros (expressing GFP) from wild type animals was co-cultured with the gonads from wild type or *Lhx2* knockout mice and tested for the migration of GFP positive cells in the gonadal region. Previous studies have established that in such gonad recombination assay, the migrating cells are endothelial in origin (Brennan et al., 2002; Capel and Batchvarov, 2008; Coveney et al., 2008). We observed that the GFP-positive endothelial cells from the WT mesonephros migrated into the *Lhx2*-/-ovaries; no such migration was observed in WT XX gonads. Thus, *Lhx2* is essential for suppressing endothelial cell migration and preventing vascularization in the ovaries. Using the gonad recombination assay, a study had shown that meiotic germ cells suppressed mesonephric cell migration in the developing ovary in vitro (Yao et al., 2003). However, an *in vivo* proof for such an event to occur was lacking. To the best of our knowledge, our study provides the first *in vivo* evidence on the role of germ cells in the control of endothelial cell migration.

Since the loss of *Lhx2* leads to angiogenesis in the ovary by a cell non-autonomous control, we next aimed to identify the factors that might promote this process. Previous studies have shown that *Wnt4, Rspo1*, and their downstream effector β-catenin are essential to suppress vascularization in XX gonads (Chassot et al., 2014). Further, GSEA revealed that the Wnt pathway and -catenin binding is overrepresented in the *Lhx2*^flox/flox^ XX gonads. Thus we hypothesized that the ectopic migration of endothelial cells in the XX gonads lacking *Lhx2* could be due to loss of *Wnt4, Rspo1*, or [*-catenin*. However, RNAseq data at E12.5 and qPCR analysis at E13.5 revealed that the levels of *Wnt4, Rspo1*, and *Ctnnb1* mRNA are identical in the *Lhx2*-/- and WT ovaries. These results imply that the migration of endothelial cells in the *Lhx2*-/-XX gonad is not due to loss of *Wnt4, Rspo1*, or β*-catenin*. This is not completely unexpected as *Wnt4* and *Rspo1*are expressed by the somatic cells of the gonads while *Lhx2* is in the germ cells. Thus, *Lhx2* independent of Wnt/β-catenin signaling pathways suppresses angiogenesis in the developing fetal ovaries. Indeed, the expression of *Lhx2* is unaltered in developing ovaries knockout for *Wnt4* (Supplementary Fig 3F). Thus it appears that both *Lhx2* in germ cells and *Wnt4* in somatic cells independently operate to suppress angiogenesis in the developing ovaries.

In the developing testis, several molecular pathways and cellular players are implicated in vascular development. It is speculated that in the developing ovary, long-range factors secreted by the meiotic germ cells in the XX gonads suppress mesonephric cell migration (Yao et al., 2003). However, the identity of such a factor(s) is yet not known. In the developing testis, loss of *Vegf* or blocking of *Vegfr2* completely prevents endothelial cell migration (Cool et al., 2011). Interestingly, disruption of PDGF does not block endothelial cell migration; though it does suppress vascular branching in the developing testis (Cool et al., 2011). Interestingly, in breast cancer cells, LHX2 directly controls the expression of *Pdgfb* by binding to its promoter and controlling endothelial cell migration (Kuzmanov et al., 2014). Beyond VEGF and PDGF, the members of EGF, FGF (*Fgf1* and *Fgf2*), Inhibin and angiopoietin, and the ephrin family are also known to activate angiogenesis in multiple tissues (Gowdak and Krieger, 2018; Gu et al., 2021; Omorphos et al., 2021; Yao et al., 2006). To determine if the migration of endothelial cells in the ovaries in absence of *Lhx2* is due to altered expression of these pro-angiogenic factors, we mined the RNAseq data and observed that there is a significant increase in expression of *Vegfa*, inhibins, *Fgf1*, and *Fgf2* in the *Lhx2* knockout ovaries and their levels are similar or even surpassing to those of the WT testis. As compared to ovaries, several angiogenesis-related genes such as angiopoietins are expected to be downregulated in the testis. Interestingly, these were also downregulated in the *Lhx2*-/-ovaries. Overall in the context of the genes that control endothelial cell migration and angiogenesis, *Lhx2* knockout XX gonads had an expression profile similar to that of the WT testis. Interestingly, we found that amongst these pro-angiogenic factors, *Pdgfa* and *Egf* are expressed by the germ cell in the XX gonads. At present, it is unknown if the increased expression of pro-angiogenic factors in absence of *Lhx2* is due to their altered levels in the germ cells. Nevertheless, our results indicate that in the absence of *Lhx2*, migration of endothelial cells in the developing ovaries is non-autonomously controlled by the germ cells creating an intra-gonadal milieu that is pro-angiogenic similar to that of the testis.

According to the current concepts of sex determination, somatic cells initially take a sex-specific fate which is then relayed to other cells in the developing gonads. The process of vascularization is also thought to be a consequence of such sex-specific fate decisions of the somatic cells to the Sertoli cells. This is evident from the fact that, in the *Wnt4* and *Rspo1* knockout gonads, a subset of somatic cells overexpresses Sertoli cell markers indicative of somatic sex reversal and there is ectopic vascularization (Chassot et al., 2014; Ungewitter and Yao, 2013). However, we did not observe any over-expression of Sertoli cell-specific genes in the *Lhx2*-/-XX gonads while there was mesonephric migration of endothelial cells. These observations indicate that sexual dimorphism of the somatic cells and endothelial cells could be independently governed events in the developing gonads. Although the supporting lineage of the *Lhx2* knockout ovaries is not masculinized, it will be of interest to investigate the effects of loss of *Lhx2* on other genes in the somatic and interstitial lineage to investigate the roles of germ cells in somatic cell programming.

It is known that the migrating endothelial cells are capable of developing a vascular network in the developing testis, while the endothelial cells in the developing ovaries do not develop a vascular network (Coveney et al., 2008). Intriguingly, this ability of the endothelial cells to form a vascular network in the two sexes is associated with the differential expression of several (445) genes in these cells. Microarray analysis has identified a sex-biased expression of several genes in the endothelial cells isolated from XX and XY gonads (Jameson et al., 2012). To determine if the transcriptome changes in the endothelial cells in the XX gonads of *Lhx2*-/-mice, we profiled the expression of endothelial cell-enriched genes in the WT XX, WT XY, and *Lhx2*-/-XX gonads. As expected, in the control animals the expression of a proportion of endothelial genes was sex-biased with higher levels in the XY gonads as compared to the XX gonads at E12.5. We next compared the expression of all these genes in *Lhx2*^flox/flox^ ovaries and observed that in the XX gonads of *Lhx2* knockout embryos, a reasonable proportion of endothelial cell-enriched genes had a male-biased pattern of expression. For example, *Cav1* is expressed at higher levels in endothelial cells of the developing ovaries as compared to the testis (Bullejos et al., 2002; Piprek et al., 2017), the expression of *Cav1* was reduced in the *Lhx2* knockout XX gonads at levels almost similar to those of the testis. In contrast, *Esm1* is an endothelial cell-specific gene overexpressed in the XY gonads as compared to XX gonads (Jameson et al., 2012). Our results showed that in *Lhx2* knockout XX ovaries, *Esm1* was elevated similar to those of the testis (Supplementary Fig 5). These results suggest that in absence of *Lhx2* in the developing ovaries, the endothelial cells have a gene expression profile like that in the developing testis. Whether this male-like expression of the endothelial cell-enriched genes in the developing ovaries of *Lhx2* mutants is due to overexpression of the pro-angiogenic factor or is induced by other factors needs to be determined.

In the developing testis, the endothelial cells have a restrictive path of migration and form a distinct male-specific vascular pattern that is not detected in the ovaries. Time-lapse, as well as static imaging studies, have shown that in the XX gonads, individual endothelial cells undergo active movements, and few endothelial cells even venture into the coelomic domain but fail to establish any recognizable coherent structures (Coveney et al., 2008). In contrast, in the XY gonads, the endothelial cell migration is directed toward the coelomic domain where cells aggregate to form the coelomic vessel (Coveney et al., 2008). We tested if the endothelial cells that have migrated in the *Lhx2-/-* XX gonads can develop a vascular pattern similar to that of XY gonads. Towards this, we extended the gonad recombination assays until 72h that allowed the endothelial cells to form a distinct network in the XY gonads. We then quantified the length and branching of GFP-positive cells that accumulated in the gonads of both WT testis and *Lhx2-/-* ovaries. As expected, GFP-positive cells in the developing testis formed a network resembling a microvasculature. Quantitatively there were distinct master segments, branches, junctions, nodes, and that ended in twigs. No such microvascular network was detected in the WT XX gonads except for a few isolated segments (not shown). Interestingly, akin to the testis, GFP positive WT cells were seen to coalesce in the *Lhx2-/-* ovaries and developed a vascular network. Quantitatively, the numbers of master segments and segments representing the vasculature networks were comparable in the *Lhx2*-/-XX gonads and the WT testis. Albeit, in the XX *Lhx2* knockouts, the numbers of isolated segments were lower whereas the number of twigs, branches, junctions, and nodes were elevated as compared to the WT testis suggesting that vasculature is established in the ovaries in absence of *Lhx2*; the pattern might be slightly different as compared to the male gonads. These results imply that in the absence of *Lhx2*, the migrating endothelial cells in the XX gonads, to a certain extent have an ability to pattern the vasculature similar to that in the testis.

In the developing testis, the vascular endothelial cell network culminates into a single coelomic blood vessel that lines the ventral aspect of the gonad. Detailed imaging experiments have shown that endothelial cells that detach from mesonephric vascular plexus actively migrate through the testis where they rearrange to form a blood vessel subjacent to the coelomic epithelium (Coveney et al., 2008). In the case of *Lhx2-/-* ovaries, although there was a migration of cells that organized into a capillary network similar to that of the testis, we failed to detect a coelomic vessel at E13.5. This was not due to a delay in the organization, as even at E15.5 when the testis had a well-patterned vascularization, the endothelial cell that migrated into the *Lhx2-/-* ovaries failed to assemble into a coelomic blood vessel. Instead, we observed hemorrhagic areas due to the accumulation of blood in the *Lhx2-/-* ovaries. To determine why the coelomic blood vessel failed to form in the *Lhx2-/-* ovaries, we analyzed the RNAseq data and observed that although there is a significant increase in the expression of most angiogenesis factors, expression of *Pdgfa* and *Pdgfb* which are crucial for branching morphogenesis in the developing testis (Cool et al., 2011) were not elevated in the *Lhx2*-/-ovaries to the levels seen in the testis. Thus, the failure to form a coelomic blood vessel could be due to an altered ratio of PDGF and VEGF in the *Lhx2-/-* ovaries. It will be of interest to study the effects of supplementation and suppression of these factors to prove this speculation.

## Conclusion

To date, the supporting lineage and the interstitial lineage were thought to control vascularization in the developing testis and/or its suppression in the developing ovaries. Our results have hitherto identified the involvement of the germ cells in this control. We show that *Lhx2* in the germ cell of the developing XX gonads creates an intragonadal milieu that suppresses vascularization. We believe that the knowledge generated through such studies in the long term will be useful in the development of management strategies for disorders of sex development and infertility.

## Funding and Acknowledgement

We thank Prof. Shubha Tole, TIFR for providing *Lhx2*-/- and *Lhx2*^flox^ mice. The project was funded by the Department of Biotechnology (DBT) Government of India under the Grant ID: BT/PR10368/MED/97/223/2014 to DM. The DM lab is also funded by grants from the Indian Council of Medical Research (ICMR) Government of India. NS is thankful to ICMR-Senior Research Fellowship. DS is thankful to UGC for the Junior Research Fellowship and AB is a recipient of a Junior Research Fellowship from the DBT project. MKJ was supported by the Ramanujan Fellowship awarded by the Science and Engineering Research Board (SERB), Department of Science and Technology (DST), Government of India (SB/S2/RJN-049/2018). The manuscript bears the NIRRH ID: RA/1123/09–2021.

## Conflict of Interest/ Declaration of interest

The authors declare no conflict of interest.

## Notes

### Competing Interest Statement

The authors have declared no competing interest.

