## Supplementary figures and images for "*Lhx2* in germ cells suppresses endothelial cell migration in the developing ovary"

### Suppl Fig 1

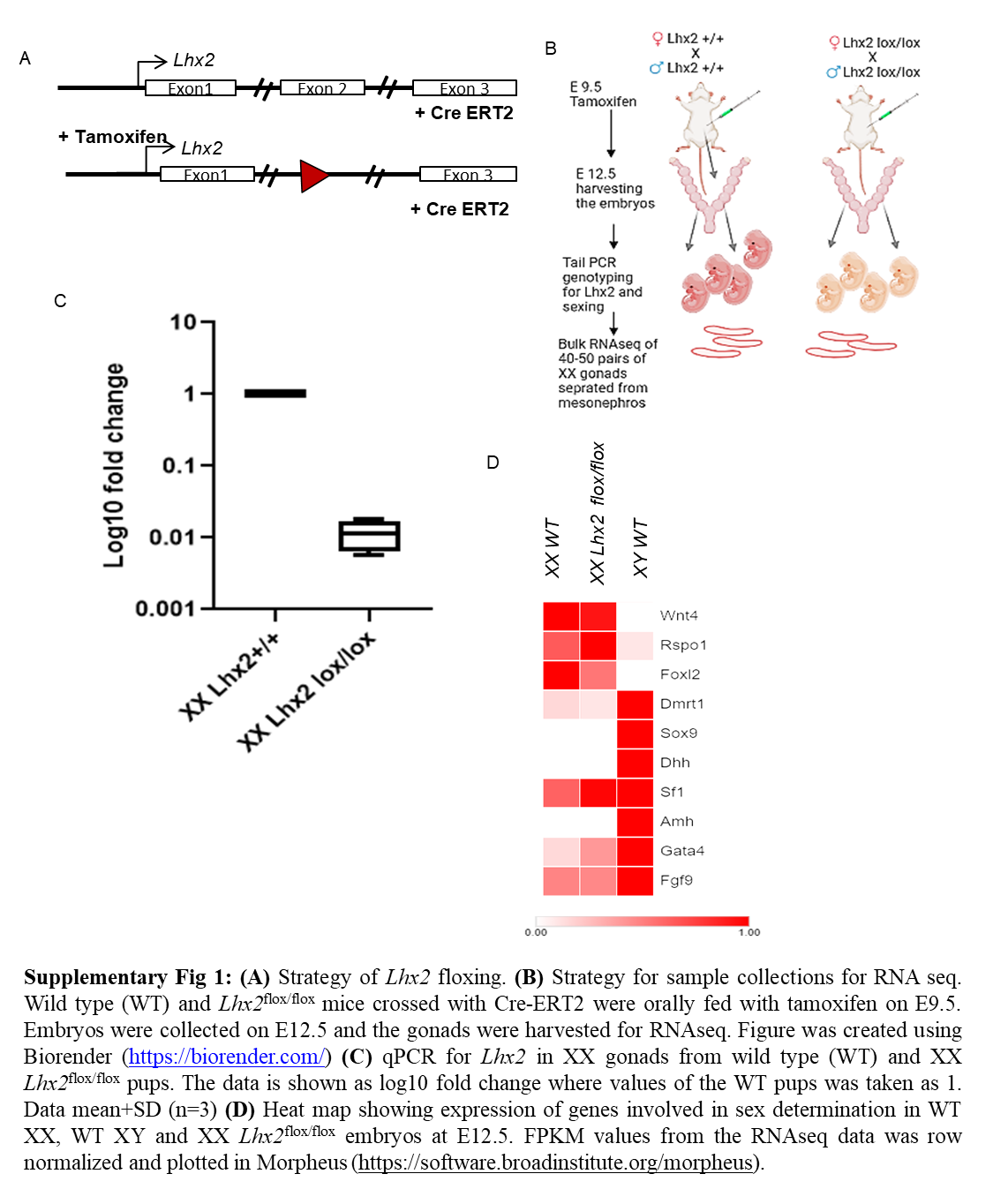

### Suppl Fig 2

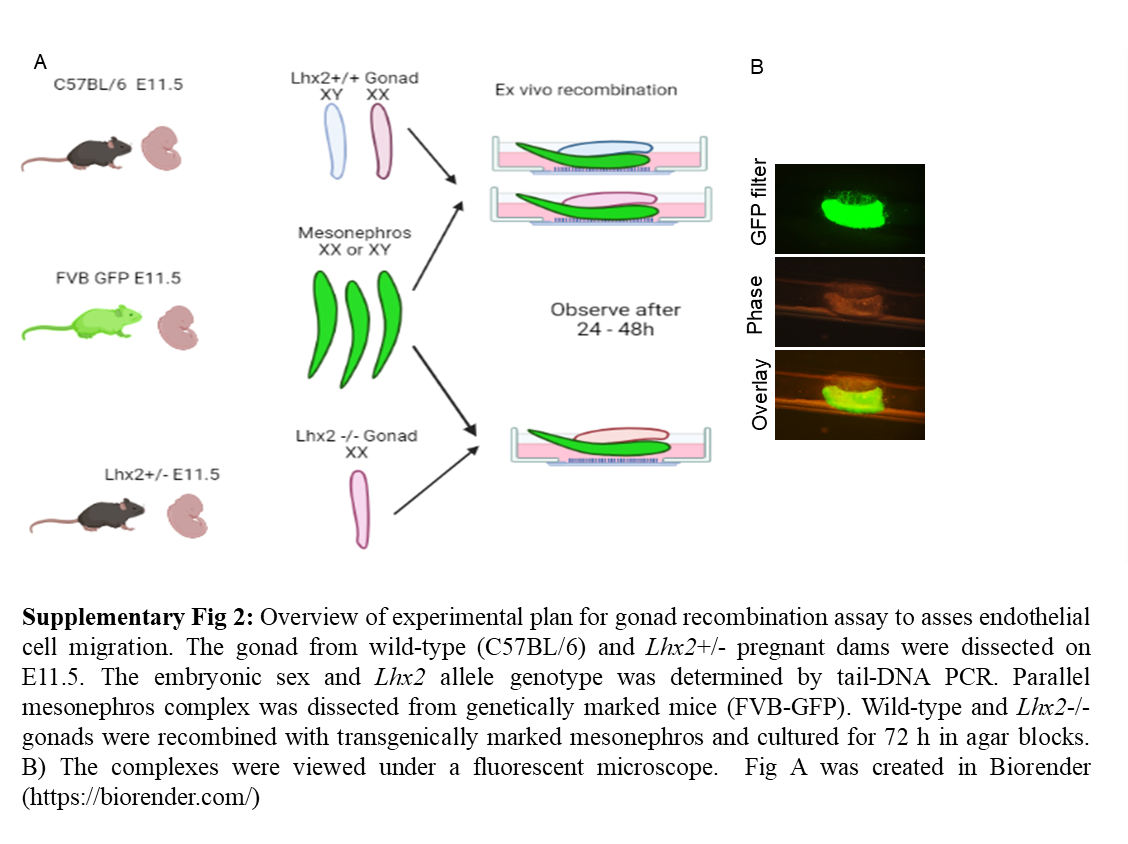

### Suppl Fig 3

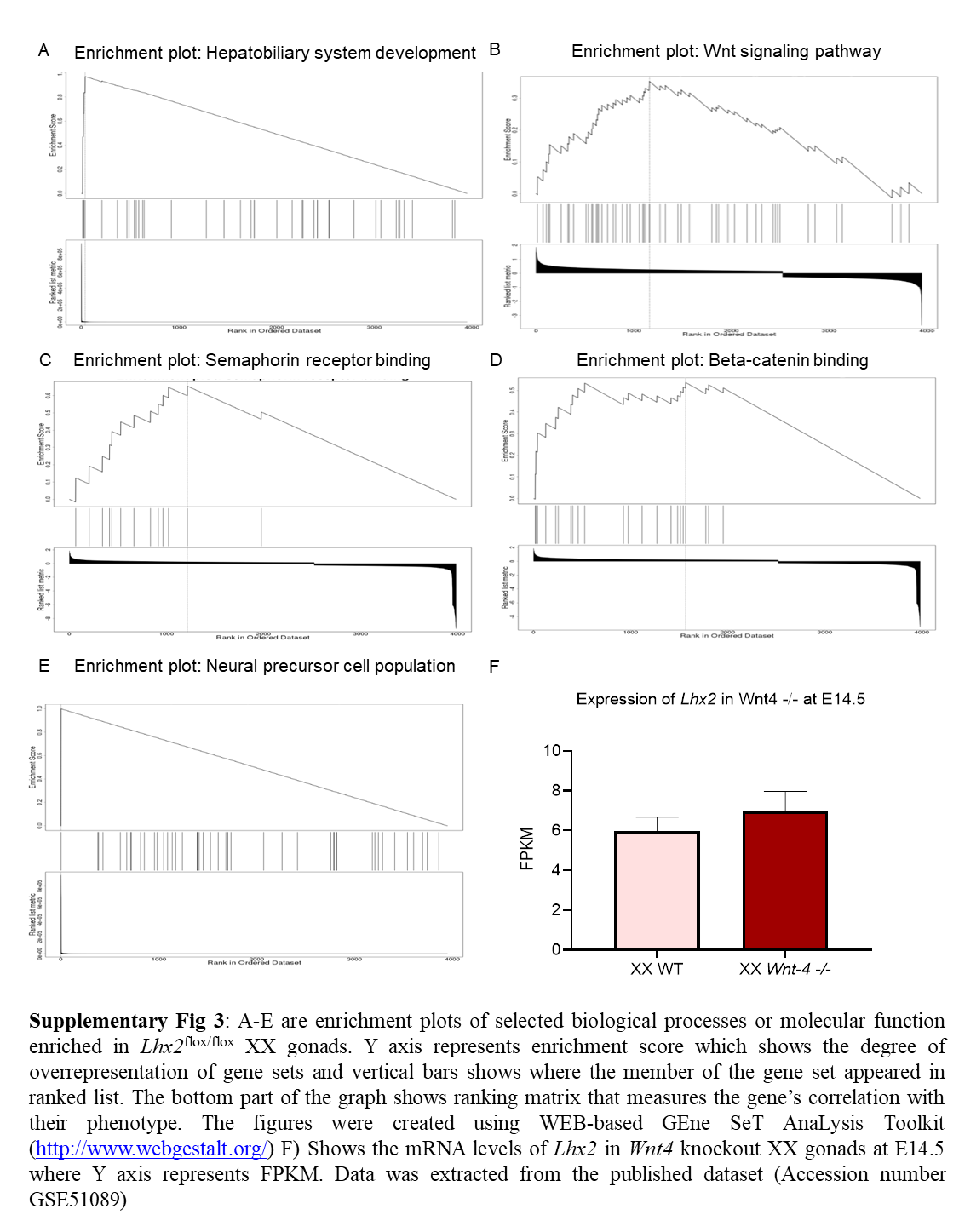

### Suppl Fig 4

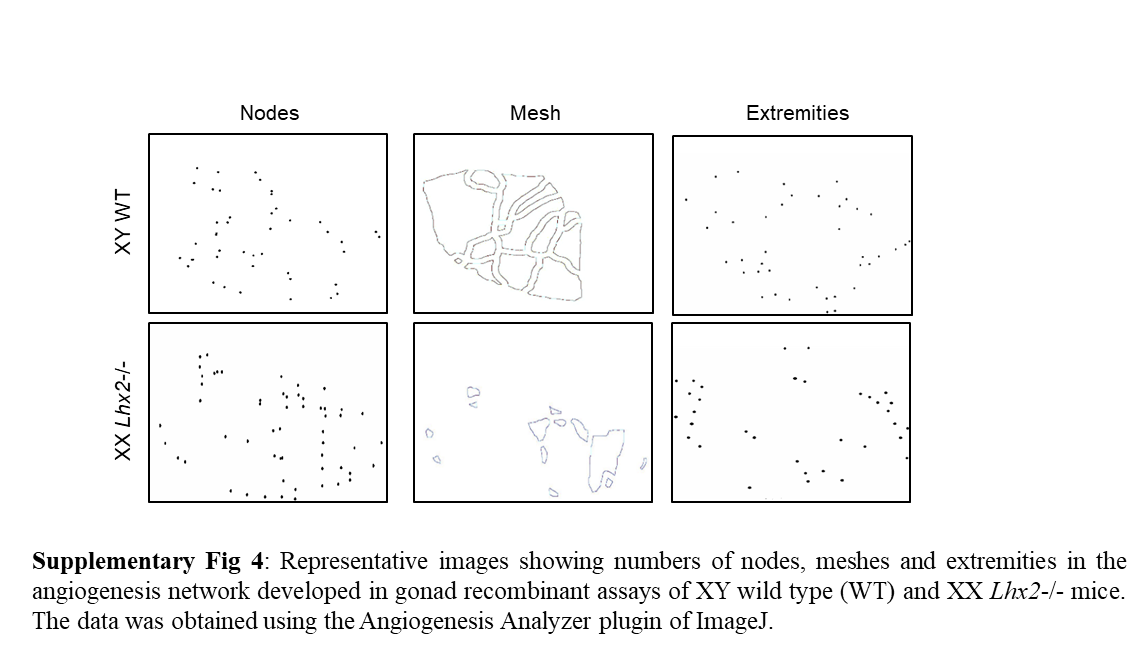

### Suppl Fig 5

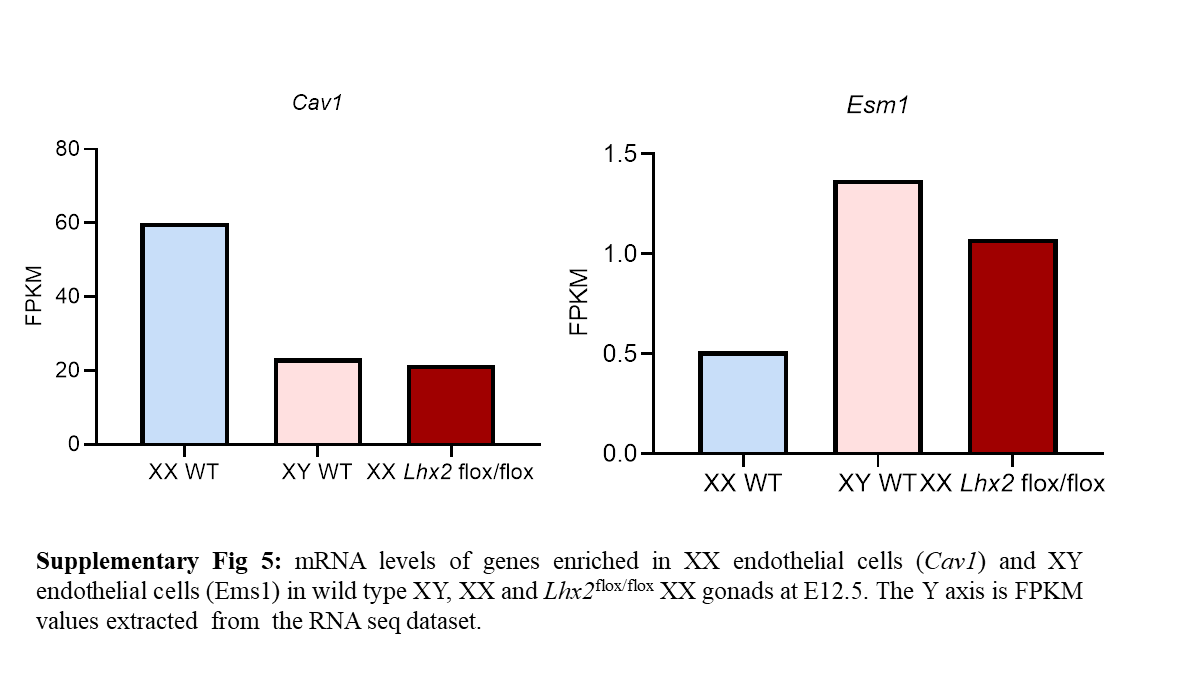
