## Supplementary material for "*Lhx2* in germ cells suppresses endothelial cell migration in the developing ovary": Suppl Table 1

**Supplementary Table 1: Accession number and datasets details used in this study**

| | Species | Dataset Accession number | Reference |
|---|---|---|---|
| 1. | Mice | E-MTAB-6798 | (Cardoso-Moreira et al., 2019) |
| 2. | Rat | E-MTAB-6811 | (Cardoso-Moreira et al., 2019) |
| 3. | Human | E-MTAB-6814 | (Cardoso-Moreira et al., 2019) |
| 4. | Lizard (*Pogona vitticeps)* | PRJNA699086 | (Whiteley et al., 2021) |
| 5. | Turtle ( *Trachemys scripta elegans*) | SRP079664 | (Czerwinski et al., 2016) |
| 6. | Chicken | E-MTAB-6769 | (Cardoso-Moreira et al., 2019) |
| 7. | Opossum (*Monodelphis domestica*) | E-MTAB-6833 | (Cardoso-Moreira et al., 2019) |
| 8. | Mice | SRP076584 | (Zhao et al., 2018) |
| 9. | Human | GSE116278 | (Lecluze et al., 2020) |
| 10. | Flk1-mCherry tag endothelial cells of mice | GSE27715 | (Jameson et al., 2012) |
| 11. | Wnt4-/- E14.5 XX Mice | GSE51089 | (Naillat et al., 2015) |
| 12. | Lhx2-/- XX Mice<br>$Lhx2^{flox/flox}$ XX Mice | | |
