## Supplementary material for "*Lhx2* in germ cells suppresses endothelial cell migration in the developing ovary": Suppl Table 2

**Supplementary Table.2: Primers used in the study.** Primers with * marks are used for genotyping the embryonic sex and *Lhx2* allele.

| Gene | Sequence (5' to 3') | Tm | Amplicon Size |
|---|---|---|---|
| *Ctnnb1* | FP-5' GCGGCCGCGAGGTACCTGAA 3'<br>RP-5' GAAGGAGCTGTGGTGGTGGCA 3' | 60ºC | 192 bps |
| *Dmrt1* | FP-5' TGGAAACCAGTGGCAGATGAA 3'<br>RP-5' TTCGAGCTCTCGTTGCTCAT 3' | 64ºC | 240 bps |
| *Foxl2* | FP-5' GGCTCTTCGGGAGCGGAGGA 3'<br>RP-5' TGGCAGGAGGCGTAGGGCAT 3' | 61ºC | 163 bps |
| *Jag1* | FP-5' GCACCCGCGACGAGTGTGAT 3'<br>RP-5' TGTAGGACCTCGGCCAGGCG 3' | 61 ºC | 201 bps |
| *Jarid** | FP-5' CTGAAGCTTTTGGCTTTGAG 3'<br>RP-5' CCACTGCCAAATTCTTTGG 3' | 61ºC | 331 bps<br>302 bps |
| *Lhx2*<br>WT* | FP-5' ACCAGACTCAGGGGAAACTCAG 3'<br>RP-5' GTGACTGAACTCCGAACCATTG 3' | 61ºC | 385 bps |
| *Lhx2*<br>Null* | FP-5' ACCAGACTCAGGGGAAACTCAG 3'<br>RP-5' ATGCCTGCTTGCCGAATATC 3' | 61ºC | 550 bps |
| *Lhx2* | FP-5' CCTACCCCAGCAGCCAAAAG 3'<br>RP-5' CTGGAGGACTCTCTTGGTGAG 3' | 61ºC | 158 bps |
| *Lhx9* | FP-5' GGACCTCAAACAGCTTGCTC 3'<br>RP-5' AATTTTCAAACGTCGGGATG 3' | 60ºC | 103 bps |
| *Pdgfrα* | FP-5' GAGCGTGCTAGGGCGGAACC 3'<br>RP-5' CCCGGCCCTGTGAGGAGACA 3' | 66 ºC | 167 bps |
| *Rspo1* | FP-5' GCCGCTGCGCCAGGTCTATC 3'<br>RP-5' AGAGCCAGGCCCGGATCCAC 3' | 63ºC | 175 bps |
| *Sdha* | FP-5' TTGGCGTTAACTGGGGCGTGGC 3'<br>RP-5' CCAAATGCAGCTCGCAAGCCTG 3' | 68ºC | 191 bps |
| *Sox9* | FP-5' CACAAGAAAGACCACCCCGA 3'<br>RP-5' GGACCCTGAGATTGCCCAGA 3' | 68ºC | 209 bps |
| Tek | FP-5' TCCTTGCCGCCAACTTGTAA 3'<br>RP-5' ATGGCGCCTTCTACTACTCCATA 3' | 58 ºC | 202 bps |
| *Wnt4* | FP-5' TGGACTCCCTCCCTGTCTTTGGGA 3'<br>RP-5' TCCTGACCACTGGAAGCCCTGTG 3' | 64ºC | 188 bps |
