## Supplementary material for "*Lhx2* in germ cells suppresses endothelial cell migration in the developing ovary": Suppl Table 3

**Supplementary Table 3: Summary of different measurements to analyse cell migration using angiogenesis analyser**

| | **Components** | **Descriptions** |
|---|---|---|
| 1. | Node | Represented as circle |
| 2. | Junctions | Connects more than two nodes |
| 3. | Branches | Delimited by a junction and ends on extremities |
| 4. | Segments | Length between to junctions |
| 5. | Twigs | Branches whose size is lower than the defined threshold |
| 6. | Isolated elements | Unbranched binary lines |
| 7. | Master junctions | Linking at least three master segments |
| 8. | Master segments | Consists in peaces of tree delimited by two junctions |
| 9. | Meshes | Area that enclosed by segments or master segments |
