## Supplementary material for "*Lhx2* in germ cells suppresses endothelial cell migration in the developing ovary": Suppl Table 4

**Supplementery Table 4: FPKM values of genes differentially expressed in the wild type and Lhx2 flox/fl**

| | **XX wild type** | **XX *Lhx2* flox/flox** |
|---|---|---|
| Isg15 | 0.951039 | 68.5934 |
| Usp18 | 0.270095 | 11.8579 |
| Lgals3bp | 0.779064 | 26.3361 |
| Irf7 | 0.5971622 | 18.64416 |
| Adcy7 | 0.67162142 | 20.47217 |
| Oasl2 | 0.567818 | 16.2745 |
| Oas2 | 0.3732953 | 10.51775 |
| Rtp4 | 0.934538 | 15.8761 |
| Oas1b | 0.3910934 | 5.884423 |
| Oasl1 | 0.247183 | 3.35123 |
| Ifit1 | 0.239466528 | 3.062184557 |
| Oas1a | 0.776581272 | 9.86917 |
| Pax2 | 0.521105898 | 5.924351 |
| Gbp3 | 0.383095389 | 4.282858 |
| Xaf1 | 0.754065353 | 7.848782 |
| Ifi47 | 0.25263166 | 2.569570124 |
| Dvl3 | 1.51359 | 15.36969 |
| Trim30a | 0.288404132 | 2.9144 |
| Pnliprp2 | 2.06818 | 19.0404 |
| Irgm1 | 2.0025703 | 17.85505 |
| Ifit3 | 0.301086081 | 2.65298 |
| Cd1d2 | 0.286679188 | 2.41737762 |
| Esr1 | 1.8192071 | 15.33008401 |
| Nfix | 0.240877324 | 2.0124343 |
| Gbp9 | 0.396984022 | 3.196112251 |
| Ifi204 | 0.254466 | 1.99009 |
| Syngap1 | 1.162189605 | 8.767018019 |
| Trim12a | 0.273212 | 2.00342 |
| Parp10 | 0.830175123 | 5.727104 |
| Fn3krp | 0.68988794 | 4.71363 |
| Irgm2 | 0.835855 | 5.5832 |
| Lmtk3 | 0.550593506 | 3.6624833 |
| Sned1 | 0.451607 | 2.97442 |
| Ntng2 | 0.4430251 | 2.876016966 |
| Cyp4f17 | 0.450811144 | 2.8988714 |
| Stat1 | 2.280198 | 14.56357 |
| Ikzf4 | 0.2514156 | 1.604050877 |
| Lgals9 | 6.7126054 | 42.4915965 |
| Akna | 0.442702286 | 2.714673054 |
| Gltscr1 | 1.097732488 | 6.6160779 |
| Kdm6b | 1.903523295 | 11.39861 |
| Upf1 | 3.70545 | 21.739978 |
| Pcdhgb4 | 0.190082 | 1.06117 |
| Jph4 | 0.247061 | 1.33916 |
| Irf9 | 4.70447 | 25.45581 |

| | | |
|---|---|---|
| Shroom3 | 1.37023153 | 7.397843 |
| Map3k14 | 0.241681 | 1.2971 |
| Slc37a1 | 0.735034 | 3.916888692 |
| Arhgap8 | 1.610877073 | 8.559250753 |
| Tet2 | 0.644219709 | 3.3936872 |
| Mir301 | 0.536707 | 2.81898 |
| Arhgef11 | 1.074954179 | 5.635526392 |
| Rsad2 | 0.957578 | 4.88569 |
| Scgb1a1 | 0.422354 | 2.15236 |
| Parp14 | 0.259531647 | 1.320824 |
| Nos1 | 0.217785 | 1.08878 |
| Dtx3l | 0.508482 | 2.51605 |
| Gigyf1 | 3.00911064 | 14.74544 |
| Msi1 | 2.701917 | 13.204186 |
| H2-T23 | 3.31315 | 15.7363 |
| Ankrd52 | 2.09275 | 9.90159 |
| Plekhn1 | 0.5208067 | 2.428659 |
| Spen | 1.13466838 | 5.288713 |
| Fcgr4 | 0.4680371 | 2.12973 |
| Sema5b | 2.21556 | 9.99937 |
| Alpk1 | 0.326968 | 1.47093 |
| Prr12 | 3.19303 | 14.2556 |
| Lmx1b | 0.238708 | 1.06075 |
| Parp9 | 1.52327 | 6.76769 |
| Gabre | 0.352097 | 1.5382653 |
| Fam160b2 | 1.292707961 | 5.6054343 |
| Ssc5d | 0.373423 | 1.61446 |
| Kmt2e | 1.766815 | 7.56814 |
| Lrp5 | 3.6117 | 15.3947 |
| Pcdhga3 | 0.27914 | 1.18405 |
| Reep6 | 0.97106 | 4.117823 |
| Dnm1 | 4.1017 | 17.2354984 |
| Srsf4 | 2.13451 | 8.92953 |
| Srcap | 3.30762621 | 13.761394 |
| Sp110 | 0.28041074 | 1.16592592 |
| Nos1ap | 0.693595 | 2.863885 |
| Tmem59l | 0.425373837 | 1.739693051 |
| Kmt2d | 1.619245 | 6.60777 |
| Pcdhgb6 | 0.552406 | 2.24836 |
| AI450353 | 0.551891 | 2.24444 |
| Gm15545 | 0.712935 | 2.89025 |
| Epha8 | 0.246802 | 0.999468 |
| Mapk15 | 0.709045904 | 2.8609 |
| Helz2 | 0.7155632 | 2.874991 |
| Pkd1 | 1.15859 | 4.64624 |
| Zfp169 | 0.2575469 | 1.032467244 |
| Fan1 | 0.476835209 | 1.911554 |

| | | |
|---|---|---|
| Prrt2 | 0.845352 | 3.37222 |
| Cbl | 1.10775 | 4.40011 |
| Igtp | 1.8868 | 7.41871 |
| Padi2 | 0.594143 | 2.33062 |
| Nckap5l | 3.41002 | 13.301395 |
| Begain | 1.14193 | 4.44504 |
| Grik1 | 1.051609556 | 4.081274 |
| Trank1 | 0.947352949 | 3.676208858 |
| Ahdc1 | 4.3061159 | 16.673016 |
| Cybb | 0.279713 | 1.08255 |
| Nlrp9b | 0.29161832 | 1.125146536 |
| Scaf1 | 8.453254 | 32.58797 |
| Wnt9b | 0.964618 | 3.71821 |
| H2-K1 | 5.541667 | 21.307169 |
| Tap1 | 1.021375175 | 3.90742707 |
| Zgrf1 | 0.55482913 | 2.10951016 |
| Myh9 | 5.71665 | 21.6543 |
| Adnp | 2.602098 | 9.8429928 |
| Gbp7 | 0.328369 | 1.238125093 |
| Apol10b | 0.374811 | 1.40989 |
| Tcof1 | 3.52524 | 13.25884 |
| Pcdhga7 | 0.607336 | 2.28231 |
| Adar | 3.0028513 | 11.22239 |
| Mtr | 3.624049438 | 13.41142308 |
| Tfap2a | 0.27111524 | 1.00236323 |
| Zfp605 | 0.78117614 | 2.878283525 |
| Spata5l1 | 1.32515 | 4.86402 |
| Mdga1 | 0.358466154 | 1.315159 |
| Nr2c2 | 0.674719 | 2.4726 |
| Cntnap1 | 0.4292061 | 1.572879958 |
| Atxn2l | 7.837209976 | 28.679916 |
| Zfp408 | 1.40173982 | 5.119893 |
| Themis2 | 0.337875 | 1.22664 |
| Med12 | 1.509663428 | 5.460294237 |
| Dagla | 0.518192 | 1.86713 |
| St8sia2 | 2.370828 | 8.51707 |
| Lrrc32 | 0.768564062 | 2.75773 |
| Prx | 0.728481159 | 2.600605 |
| Hap1 | 0.802968063 | 2.861068634 |
| Cdc42bpg | 0.912023444 | 3.2438 |
| Ppard | 2.96168 | 10.5274 |
| Soga1 | 2.2544588 | 8.00413 |
| Agap2 | 0.4353014 | 1.54342169 |
| Bst2 | 88.842 | 314.574 |
| Iigp1 | 0.853630582 | 3.016699 |
| Plekhg2 | 5.4093819 | 19.08954 |
| Gm5901 | 0.431822 | 1.52112 |

| | | |
|---|---|---|
| Wnt7a | 0.95674453 | 3.359135 |
| D930007P1 | 0.7782673 | 2.73045 |
| Psmb9 | 1.03589 | 3.63248 |
| Proser3 | 1.3009839 | 4.553769 |
| Uprt | 0.330789 | 1.15698 |
| Tnrc18 | 5.37031929 | 18.75465825 |
| Frem2 | 1.564714398 | 5.46234909 |
| Lhx1 | 0.953068 | 3.3240949 |
| Foxp4 | 5.720204344 | 19.80823826 |
| Pcdhgc3 | 6.6196 | 22.81 |
| Rusc2 | 1.1690721 | 4.01985 |
| Fam163a | 0.389181 | 1.33407 |
| Sipa1l3 | 1.576166555 | 5.3922411 |
| Fat3 | 0.856262764 | 2.922297881 |
| Fbxo44 | 0.681434885 | 2.325117382 |
| AW549542 | 1.61843 | 5.50341 |
| Syt6 | 0.332115248 | 1.120758754 |
| Zfp335 | 2.24463 | 7.53552 |
| Csf2ra | 2.97074738 | 9.9655471 |
| Ccl9 | 1.02923 | 3.44351 |
| Lrch3 | 1.38351 | 4.60848 |
| Pcdhga2 | 0.404782 | 1.34788 |
| Cmip | 2.475382 | 8.2332 |
| Col7a1 | 0.61415594 | 2.0410617 |
| Ifitm3 | 98.8604 | 325.336 |
| Pde7b | 0.557967 | 1.829404 |
| Notch3 | 3.164185212 | 10.37214 |
| Gm15800 | 1.005835736 | 3.28975314 |
| Ntn1 | 1.03438 | 3.35911 |
| Gli1 | 3.1559971 | 10.246939 |
| Pnliprp1 | 4.40727 | 14.23933 |
| Adam1a | 0.29808 | 0.958045 |
| Cic | 9.580604 | 30.72459 |
| Zc3h3 | 3.652607 | 11.70459216 |
| Mocs1 | 1.870002 | 5.964946 |
| Hk3 | 0.400808917 | 1.277006252 |
| Zfpm1 | 4.1087837 | 13.06816 |
| Srrm2 | 12.36741 | 39.2888 |
| Szt2 | 1.83773 | 5.83531 |
| Nyap1 | 1.044937828 | 3.3155982 |
| Trim34a | 0.822774 | 2.6105489 |
| Erdr1 | 51.633754 | 162.43844 |
| Ltbp4 | 6.032471 | 18.9678378 |
| Ddx58 | 2.616066 | 8.19961 |
| P3h2 | 0.86765 | 2.718513 |
| Hivep1 | 1.673681 | 5.240416932 |
| Ccdc85c | 5.33953 | 16.71465 |

| | | |
|---|---|---|
| Slfn2 | 1.21153 | 3.79216 |
| Spred3 | 0.986135 | 3.083502 |
| Cmpk2 | 1.0461 | 3.26444 |
| Psmb8 | 1.90514 | 5.91708 |
| Hif3a | 3.13953921 | 9.749351 |
| Adamts2 | 2.187631266 | 6.78714 |
| E130102H2 | 0.591243 | 1.83117 |
| Asic2 | 1.437415536 | 4.4481 |
| Igsf9 | 3.163314419 | 9.786021 |
| Eif2ak2 | 1.895061 | 5.84225265 |
| Arhgap33 | 1.692827984 | 5.214510386 |
| Sf3a2 | 11.0868 | 34.0602 |
| Cep250 | 1.279455811 | 3.929777874 |
| Zfp423 | 2.593508492 | 7.9571598 |
| Khsrp | 11.1637 | 34.2403 |
| Lrrtm1 | 2.3421221 | 7.17765271 |
| Ptch2 | 2.0045207 | 6.12879 |
| Cnot3 | 7.185956318 | 21.96640456 |
| Sema4g | 4.29441 | 13.12112 |
| Carm1 | 10.20489 | 31.17667 |
| Chd5 | 0.361671096 | 1.104897 |
| Wnk4 | 3.0576411 | 9.338220224 |
| Thsd4 | 1.444486419 | 4.399204515 |
| Polr2a | 11.17391 | 34.0211 |
| Afap1 | 2.37198 | 7.21685 |
| Tmem181a | 1.01586 | 3.0862 |
| Plekhg3 | 2.08260636 | 6.31609483 |
| Mast2 | 5.21157974 | 15.80473457 |
| Kmt2b | 4.05928242 | 12.29953039 |
| Ptpn23 | 4.91432 | 14.8862 |
| Nav2 | 3.1020424 | 9.392001 |
| Lama5 | 2.21674652 | 6.69312 |
| Ttc28 | 1.694656 | 5.111674987 |
| Elk4 | 0.543269 | 1.635776 |
| Tram2 | 1.169327 | 3.52025 |
| Man2a2 | 5.801778 | 17.46169334 |
| Gfra1 | 0.754415891 | 2.266102362 |
| Celsr2 | 0.548187126 | 1.63526241 |
| Thy1 | 6.97626 | 20.7839 |
| Rapgefl1 | 0.958795 | 2.85278 |
| Tnxb | 0.82961031 | 2.465080194 |
| Hmbox1 | 1.67138021 | 4.946415 |
| Vps33b | 1.071781 | 3.166788996 |
| Tmem220 | 0.427861484 | 1.264003158 |
| Bcl9l | 3.935715313 | 11.61707 |
| Leng8 | 8.2585 | 24.2938 |
| Trpc2 | 0.515616 | 1.51418 |

| | | |
|---|---|---|
| Dclk1 | 2.36572584 | 6.933240238 |
| Pcsk4 | 0.840133 | 2.460489 |
| Rasgrp4 | 0.613186 | 1.786021257 |
| Cacna1h | 3.33276126 | 9.690953 |
| Pde4a | 1.184088488 | 3.442383 |
| B4galnt3 | 0.409656 | 1.190005397 |
| Taok2 | 4.695702 | 13.63081 |
| Scn2b | 0.822701077 | 2.386687589 |
| Furin | 7.018657563 | 20.3077215 |
| Mir210 | 0.422242 | 1.22086 |
| Flywch1 | 5.12533 | 14.81115 |
| Fsbp | 0.428012 | 1.23536 |
| Ankrd13b | 3.56471 | 10.28772 |
| Sv2a | 1.895947 | 5.449375374 |
| Pcdhgb7 | 0.906127 | 2.60271 |
| Mbd6 | 6.20824 | 17.8264 |
| Pcdhga11 | 0.3312 | 0.950228 |
| Clmn | 0.798052616 | 2.2851527 |
| Snx29 | 1.836131543 | 5.2516101 |
| Slc6a6 | 3.842286626 | 10.98543191 |
| Kif26a | 3.11745 | 8.91041 |
| Zcchc11 | 3.69749 | 10.5514 |
| Tbx3 | 4.68909738 | 13.3581316 |
| Bcl9 | 5.079660769 | 14.45665464 |
| Kif20a | 4.93418 | 14.026252 |
| Zfp369 | 0.733334 | 2.081156 |
| Mybpc1 | 1.27161307 | 3.602182176 |
| Kif1a | 1.066744507 | 3.021113669 |
| Tbc1d9 | 0.868469 | 2.458767 |
| Zfp646 | 2.185978 | 6.18881 |
| Slc22a15 | 0.840148 | 2.376890165 |
| Zfp385a | 4.869490453 | 13.77527907 |
| Mapkbp1 | 1.7718455 | 5.00754 |
| Cog7 | 2.33759 | 6.595896 |
| Moxd1 | 2.24204 | 6.32281 |
| Gm10560 | 0.43356 | 1.22212 |
| Scube1 | 2.4729902 | 6.953471 |
| Lrrc24 | 0.347777 | 0.976424447 |
| Fbrs | 7.922828 | 22.23318 |
| Tnfrsf25 | 0.57785929 | 1.620304109 |
| Fndc3c1 | 0.5174663 | 1.450782764 |
| Sipa1l1 | 2.751494908 | 7.688589 |
| Adgrb1 | 0.41485465 | 1.158685887 |
| Setd1b | 1.85052 | 5.16725 |
| Bahcc1 | 2.4864601 | 6.9399161 |
| Irs2 | 2.29935 | 6.4169 |
| Hoxb8 | 3.087401743 | 8.60610411 |

| | | |
|---|---|---|
| Rgma | 3.69031 | 10.2749 |
| Pnlip | 2.76638 | 7.6894 |
| Grik4 | 0.60805226 | 1.6872583 |
| Col1a1 | 33.6455 | 93.3139 |
| Kif26b | 1.68865 | 4.66213 |
| Traf3 | 1.9571112 | 5.403013 |
| Atn1 | 17.23447351 | 47.47203 |
| Scara5 | 2.7874816 | 7.6745425 |
| Prrc2a | 26.2359656 | 72.22108 |
| Lamc3 | 0.379025113 | 1.043101693 |
| Zfp609 | 2.80824 | 7.72422 |
| Lcat | 0.58624 | 1.6108 |
| Xpnpep3 | 0.7018119 | 1.926784678 |
| Peg3os | 28.398 | 77.9443 |
| Zfp142 | 1.3383011 | 3.66875215 |
| Dzip1l | 2.13178597 | 5.841258168 |
| Krt17 | 0.427117213 | 1.168105 |
| Caskin2 | 4.55419 | 12.4531 |
| Dlk1 | 48.685367 | 132.85609 |
| Hand2 | 7.06118 | 19.2669 |
| Clip3 | 12.1308753 | 33.02745 |
| Tubb4a | 2.48514 | 6.76094 |
| Maml3 | 1.5693831 | 4.26934 |
| Pitpnm3 | 0.892355745 | 2.426466001 |
| Ctsk | 0.859606 | 2.33525 |
| Pitpnm2 | 2.739211673 | 7.43692396 |
| Ntrk2 | 3.097167 | 8.4011273 |
| Cacng7 | 7.596586 | 20.59623 |
| Ppfia4 | 1.003179756 | 2.715275064 |
| Tns2 | 5.33405487 | 14.42208 |
| Ttc22 | 0.678820215 | 1.831415893 |
| Slc7a8 | 2.0439486 | 5.51352 |
| Micall1 | 1.762813 | 4.753857457 |
| Foxk1 | 2.79906 | 7.54433 |
| Xrcc3 | 1.61707 | 4.35813 |
| Fam219a | 0.98025967 | 2.641421 |
| Wiz | 9.32118292 | 25.0415185 |
| Bgn | 22.0921 | 59.2808 |
| Lmx1a | 0.774634507 | 2.075531216 |
| Sugp2 | 2.604346 | 6.975949 |
| Tmem108 | 2.05652329 | 5.504903 |
| Snhg17 | 2.4375736 | 6.519087 |
| Adm | 4.07266 | 10.8854 |
| Slc4a3 | 6.0035934 | 16.0380447 |
| Arhgap39 | 2.402806278 | 6.411546515 |
| Ctnna2 | 0.52809408 | 1.408099 |
| Sympk | 13.6327 | 36.3267 |

| | | |
|---|---|---|
| Ptger1 | 2.50375 | 6.66522 |
| Hoxd4 | 5.711312092 | 15.164465 |
| P2rx2 | 0.387772 | 1.0282293 |
| Npff | 0.787687 | 2.088629 |
| Tonsl | 2.93177533 | 7.76987211 |
| Pianp | 0.692791797 | 1.833643731 |
| Hoxd3 | 1.28614 | 3.40342 |
| Zmiz2 | 9.5434176 | 25.253453 |
| Wdr62 | 1.1009642 | 2.91293999 |
| Ciz1 | 5.4040976 | 14.295005 |
| Arid1a | 8.947158 | 23.65382 |
| Gpr4 | 0.384654 | 1.0169 |
| Ssh1 | 2.80268 | 7.39766 |
| Myo19 | 1.647283 | 4.345437042 |
| Kmt2a | 1.75526464 | 4.6265531 |
| Sema6a | 2.27692 | 5.99957 |
| Tln1 | 7.49381 | 19.74365 |
| Med13l | 1.78861367 | 4.708742094 |
| Acot2 | 1.6113 | 4.23666 |
| Ifi35 | 3.28757 | 8.63816 |
| Peli3 | 0.546830492 | 1.43210215 |
| Smg7 | 5.894282 | 15.42536 |
| Sdk2 | 1.092533172 | 2.858641235 |
| Ece1 | 5.575118192 | 14.576367 |
| Tns1 | 6.08661252 | 15.907278 |
| Cep164 | 4.394546223 | 11.483957 |
| Adamts12 | 1.3145114 | 3.43166493 |
| Rexo1 | 5.10863 | 13.3307 |
| Fbrsl1 | 6.03037 | 15.70839 |
| Ddi2 | 4.331354 | 11.27949 |
| Gpatch2 | 1.72049 | 4.48005 |
| Zmiz1 | 5.8373812 | 15.19427 |
| Nup210 | 2.182633 | 5.674369 |
| Spock2 | 1.57541 | 4.09475 |
| Fmnl1 | 0.497260077 | 1.29179142 |
| Ptprs | 25.15689256 | 65.195443 |
| Osr2 | 8.76777 | 22.71359 |
| Arhgef1 | 10.60244961 | 27.4641507 |
| Arid1b | 3.591066887 | 9.300446727 |
| Lrig3 | 1.93822134 | 5.011839 |
| Mtss1l | 12.50106751 | 32.30165 |
| Polrmt | 3.70733 | 9.57546 |
| Runx2 | 0.531647 | 1.3728512 |
| Adamts4 | 0.497158 | 1.283118 |
| Casz1 | 0.400711 | 1.033211 |
| Tmem151a | 2.27087 | 5.8549 |
| Chst3 | 0.691414498 | 1.782105 |

| | | |
|---|---|---|
| Fam171a2 | 13.50904 | 34.81895 |
| Raver1 | 17.1834 | 44.2708 |
| Sema5a | 2.71998 | 7.0018 |
| Kansl1 | 5.2775723 | 13.585545 |
| Ncapd2 | 20.37846 | 52.4492 |
| Nab2 | 7.24888 | 18.64863 |
| Fgf1 | 0.395517 | 1.01576 |
| Grb7 | 2.76476 | 7.09465 |
| Myo1f | 0.637526 | 1.63502 |
| Kat6b | 3.207051759 | 8.223738 |
| Ccdc157 | 1.688027104 | 4.322382895 |
| Sp2 | 2.295207 | 5.868181 |
| Plekha6 | 1.744857671 | 4.46025703 |
| Tcf7l1 | 7.599128534 | 19.41892782 |
| Adamtsl4 | 0.424153516 | 1.08333711 |
| Slitrk1 | 0.645313 | 1.648151 |
| Trim14 | 1.068221885 | 2.727916155 |
| Elfn1 | 1.37523763 | 3.5065021 |
| Tubg2 | 1.099 | 2.80173 |
| Fxyd4 | 1.27944189 | 3.25809 |
| Mink1 | 4.88706 | 12.44279887 |
| Zbtb12 | 9.6604 | 24.5958 |
| Snapc4 | 2.3175778 | 5.895686032 |
| Asap2 | 1.7043104 | 4.334969939 |
| Ypel4 | 0.923618 | 2.348522 |
| Hck | 0.590048782 | 1.498629 |
| Plxnb1 | 5.534933789 | 14.04406 |
| Spsb1 | 1.50872 | 3.82805 |
| Cramp1l | 2.47793941 | 6.28549993 |
| Gltscr1l | 1.334639696 | 3.384285 |
| Cacna1g | 5.87996647 | 14.90258375 |
| Camta2 | 1.749978226 | 4.4346082 |
| Calcoco1 | 11.5997 | 29.3656 |
| Papln | 2.113307 | 5.34793 |
| Nlgn2 | 10.70391 | 27.00986 |
| Msmp | 1.9023 | 4.79911 |
| Gnao1 | 0.56974 | 1.436708 |
| Evpl | 0.886443 | 2.23497 |
| Laptm5 | 3.75534 | 9.46308 |
| Zfp574 | 4.83951141 | 12.19192 |
| Sbf2 | 3.440578944 | 8.662133 |
| Gldc | 7.54168 | 18.9498 |
| Fam83h | 0.9335548 | 2.34159931 |
| Sema4b | 3.09196 | 7.75538 |
| Satb1 | 2.293953643 | 5.745998493 |
| Acrbp | 3.022681 | 7.539879 |
| Adrbk2 | 0.689505325 | 1.719327758 |

| | | |
|---|---|---|
| Fam222a | 2.04743 | 5.10363 |
| Parp4 | 0.548193 | 1.36633 |
| Rfx2 | 1.772239732 | 4.4040765 |
| Gm4349 | 0.386765 | 0.95898 |
| Hspg2 | 6.68561639 | 16.57562568 |
| Dpysl5 | 1.075247605 | 2.66480494 |
| Samd4b | 9.09921904 | 22.5473479 |
| Smoc2 | 9.166866 | 22.68493 |
| Gpatch8 | 3.24788 | 8.03581 |
| Arhgap23 | 5.349486001 | 13.233558 |
| Pcnx | 2.925907105 | 7.23444 |
| Cdh16 | 5.7694429 | 14.25179599 |
| Garem | 1.04379 | 2.57791 |
| Krba1 | 2.728803572 | 6.733175 |
| Hoxb3 | 2.852716512 | 7.03738 |
| Hcn3 | 0.433757038 | 1.068748 |
| Zfp710 | 4.636728102 | 11.4214299 |
| Clip2 | 4.466316121 | 10.99105 |
| Capn6 | 8.19518 | 20.1633 |
| Dlgap3 | 1.854397049 | 4.562000103 |
| Fhod1 | 3.71941 | 9.14722 |
| Sh3bp1 | 8.93581 | 21.972269 |
| Mef2d | 5.17781 | 12.7275102 |
| Mov10 | 15.45263987 | 37.91952 |
| Dhx37 | 3.394907 | 8.324762985 |
| Dlg4 | 6.27142764 | 15.36163 |
| Irx5 | 1.71062 | 4.18677 |
| Zscan10 | 1.038733206 | 2.5412526 |
| Sgcd | 0.5872891 | 1.436677 |
| Spaca6 | 1.271932105 | 3.108243 |
| Npy | 5.63397 | 13.7599 |
| Apobec1 | 0.70556706 | 1.723037 |
| Lrch4 | 3.82053 | 9.325037 |
| Dpp10 | 0.466807 | 1.13643 |
| Astn1 | 0.66931 | 1.629008075 |
| Rai1 | 3.5332161 | 8.584371 |
| Gtf2ird1 | 5.663758 | 13.73397608 |
| Rab11fip1 | 1.346080725 | 3.2626365 |
| Trpm5 | 1.97426 | 4.7748 |
| Atf7 | 3.269443 | 7.9023932 |
| Adgra3 | 12.5598 | 30.3233 |
| Klc2 | 3.694651584 | 8.916557 |
| Setd1a | 6.19559 | 14.90813 |
| Gon4l | 2.171664 | 5.22057563 |
| D630045J1 | 1.31594362 | 3.1623926 |
| Clu | 17.45056 | 41.89446 |
| Zfp865 | 5.390114 | 12.93645 |

| | | |
|---|---|---|
| Lrrc14b | 0.73423 | 1.76055 |
| Jade2 | 0.674356424 | 1.616498846 |
| Plekhg5 | 4.330055403 | 10.36643052 |
| Pld4 | 2.06051 | 4.93266 |
| Nol10 | 3.3769435 | 8.08304 |
| Dhx9 | 14.27033168 | 34.15185 |
| Chd6 | 4.60942 | 11.019783 |
| Fam20a | 0.507308 | 1.21257 |
| Ppp1r13l | 2.017936932 | 4.815672 |
| Stx1b | 0.900369 | 2.1469 |
| Adamts6 | 0.908947613 | 2.164864808 |
| Kcp | 5.5242217 | 13.15022 |
| Chd3 | 14.49728215 | 34.50857 |
| Ptpn3 | 0.647905783 | 1.540819607 |
| Rai2 | 1.16602034 | 2.77277 |
| Ubr4 | 6.965517 | 16.56212104 |
| Trerf1 | 2.31121963 | 5.4941323 |
| Mpeg1 | 1.166360017 | 2.771999021 |
| Ccnt1 | 1.650958 | 3.9192 |
| Nfkb2 | 2.467375612 | 5.856272095 |
| Tbc1d24 | 1.4806811 | 3.513567646 |
| Hoxb5 | 1.80606 | 4.28471 |
| Pced1b | 6.535029 | 15.49359 |
| Tmem180 | 3.54969 | 8.41031 |
| Car15 | 0.608524 | 1.44111 |
| Iglon5 | 0.947398704 | 2.242074388 |
| Gm12992 | 7.389883 | 17.4785723 |
| Sdc3 | 8.42404 | 19.9226 |
| Ankhd1 | 3.9625945 | 9.37101 |
| Git1 | 13.15611712 | 31.09914 |
| Plekhh1 | 1.39697 | 3.30157 |
| Slc38a3 | 0.98993767 | 2.338365488 |
| Scmh1 | 10.591171 | 25.01487 |
| Clasrp | 7.014493466 | 16.56715709 |
| Fry | 0.53124796 | 1.254415 |
| Arhgap32 | 1.865496201 | 4.399471 |
| Col9a2 | 6.01242 | 14.1755 |
| Cep152 | 0.4404794 | 1.038067365 |
| Sfswap | 6.11214 | 14.3939 |
| Hapln3 | 3.998088982 | 9.41442386 |
| Flnb | 5.921964536 | 13.936005 |
| Dnmbp | 4.032045 | 9.484701563 |
| Vgll4 | 7.4135 | 17.43843 |
| Dpp9 | 5.59121 | 13.1519 |
| Zfr2 | 0.926980159 | 2.176055648 |
| Ash1l | 2.479706 | 5.82036 |
| Ern1 | 1.890926 | 4.438144 |

| | | |
|---|---|---|
| Gtf3c1 | 8.7668 | 20.574 |
| Notum | 8.72513 | 20.47067 |
| Sp140 | 1.41905 | 3.329101 |
| Rgag4 | 1.35149 | 3.16997 |
| Daam2 | 7.8898395 | 18.49348 |
| Itpr3 | 2.52805 | 5.9253 |
| Tead1 | 4.13398374 | 9.6691032 |
| Khnyn | 5.57755 | 13.0435 |
| Hjurp | 1.89344 | 4.42359 |
| Abca2 | 1.92022596 | 4.48434 |
| Tmem181c | 0.49598 | 1.15827 |
| Hoxa2 | 1.20231 | 2.80713 |
| Rnf44 | 19.69580958 | 45.95479873 |
| Cul9 | 0.8034147 | 1.87363192 |
| Eml2 | 4.584627919 | 10.67984 |
| Ubap2 | 13.3325 | 31.0161 |
| Cux2 | 0.796700808 | 1.852409053 |
| Dhx8 | 5.54294 | 12.88773 |
| Plekha7 | 1.451543721 | 3.371986311 |
| Ptpru | 0.7985512 | 1.8497022 |
| Kansl3 | 7.760279 | 17.97386795 |
| Sema6c | 6.452472721 | 14.94394894 |
| Nfkbid | 0.60514442 | 1.401226283 |
| Plcb3 | 11.6595061 | 26.99450511 |
| Zfp618 | 3.815585318 | 8.82995382 |
| Med25 | 20.10733 | 46.51067 |
| Prrc2b | 13.167324 | 30.44413 |
| Ago2 | 4.03418 | 9.325 |
| Slc39a14 | 1.896787568 | 4.383086988 |
| Ttyh3 | 36.12622 | 83.4756 |
| Sdk1 | 0.898398 | 2.07419 |
| Kifc5b | 2.24308 | 5.17737 |
| Vegfa | 16.005403 | 36.940755 |
| Pnoc | 0.69304 | 1.598773 |
| Zfp526 | 1.00648 | 2.32066 |
| Akap13 | 2.0597952 | 4.748990367 |
| Trim39 | 3.163563266 | 7.2930816 |
| Rnf112 | 0.555359143 | 1.280147 |
| Lamb2 | 4.36866 | 10.06847 |
| Slc25a23 | 6.22079 | 14.3303 |
| Creb3l1 | 8.42672 | 19.41171 |
| Adra2a | 1.55468 | 3.58118 |
| Vwa2 | 0.901705 | 2.07688 |
| Peg10 | 15.492356 | 35.6681887 |
| Arhgef17 | 7.921446 | 18.23198223 |
| Tfcp2l1 | 1.07812 | 2.48079 |
| Vps13d | 1.186703963 | 2.72962358 |

| | | |
|---|---|---|
| Sipa1 | 6.969076 | 16.0216413 |
| Emilin1 | 31.7166 | 72.8398 |
| Wdr46 | 5.042045 | 11.57127 |
| Slc25a47 | 1.277158 | 2.9292003 |
| Kif12 | 3.03104 | 6.94495 |
| Klhl18 | 1.850167606 | 4.23725 |
| Gatad2b | 3.912204692 | 8.959538246 |
| Fam13c | 0.480866 | 1.1003409 |
| Gadd45g | 12.9609 | 29.6518 |
| Tapbp | 8.68602 | 19.86369 |
| Nipsnap3b | 6.844665 | 15.6499 |
| Cntrl | 1.49613493 | 3.417225369 |
| Kdm5c | 7.02567673 | 16.03089 |
| Prpf8 | 17.0731 | 38.9499 |
| Hoxaas3 | 0.493874 | 1.12574 |
| E2f8 | 1.33402825 | 3.04032 |
| Arhgef40 | 5.23369 | 11.92506 |
| Clk2-scamp | 5.925871 | 13.49678 |
| Nkd1 | 12.559 | 28.5878 |
| Btbd19 | 1.164887012 | 2.649314713 |
| Repin1 | 5.09320218 | 11.579532 |
| Brsk1 | 1.329886 | 3.022011 |
| Cep170b | 5.499212513 | 12.47755588 |
| Fzd6 | 1.377874 | 3.125889 |
| Arhgef16 | 1.02921934 | 2.33379785 |
| Rara | 8.021728346 | 18.17983679 |
| Dapk1 | 9.277920926 | 21.02067 |
| Zfp395 | 11.62581863 | 26.33369 |
| Auts2 | 2.73334146 | 6.190751213 |
| Pcnxl3 | 8.00626998 | 18.133034 |
| Mir6385 | 0.596921 | 1.35033 |
| Zfp36 | 4.80136 | 10.8502 |
| Cpm | 1.54151609 | 3.482388642 |
| Folr2 | 3.093106088 | 6.987073382 |
| D430042O( | 1.49098019 | 3.367958136 |
| Rad54l2 | 3.20948 | 7.24597 |
| H2-D1 | 12.2074 | 27.551315 |
| Cyp2d22 | 2.189097475 | 4.94004 |
| Capn15 | 4.299592 | 9.701222 |
| Dhx34 | 3.81875 | 8.610995 |
| Tert | 1.42749763 | 3.218831 |
| Esrp2 | 0.661998 | 1.49265 |
| Dok3 | 0.542621 | 1.22345 |
| Nav1 | 2.02253752 | 4.559777121 |
| Trim56 | 0.707247 | 1.593480726 |
| Stra6 | 8.677404 | 19.53607 |
| Cit | 1.439113046 | 3.23820772 |

| | | |
|---|---|---|
| Ercc4 | 1.29976 | 2.92459 |
| Helz | 3.63346 | 8.175152787 |
| Cfb | 0.577079029 | 1.298162844 |
| Nrbp2 | 3.449075 | 7.75726 |
| Cox15 | 2.382363258 | 5.35533 |
| Znf512b | 6.868116876 | 15.43478103 |
| Hk1 | 7.08727629 | 15.92282545 |
| Sox13 | 6.769712 | 15.20797651 |
| Tcirg1 | 7.038186751 | 15.792047 |
| Pip5k1c | 10.64039629 | 23.81948 |
| Atp2b4 | 3.8412341 | 8.59735 |
| Hspa12a | 1.25683 | 2.81089 |
| Osr1 | 1.69445 | 3.78553 |
| Arhgap20 | 1.387133 | 3.096871 |
| Avpr1a | 1.1621 | 2.59326 |
| Sfxn2 | 2.28915266 | 5.106424 |
| Mybbp1a | 19.3073 | 43.0542 |
| Col6a2 | 20.3696 | 45.3833 |
| Ncf1 | 0.88394879 | 1.96847676 |
| Fzd10 | 3.66118 | 8.1517 |
| Mllt6 | 5.230732556 | 11.63761 |
| Apcdd1 | 8.8092 | 19.59503 |
| Vdr | 0.886184 | 1.97098991 |
| Mical2 | 2.751632 | 6.1197521 |
| Serpinb9b | 0.489131 | 1.08754 |
| Vps13c | 0.779732821 | 1.731696471 |
| Prpf40b | 6.21893 | 13.80459 |
| Supt20 | 6.55664293 | 14.55271159 |
| Gm16023 | 0.437861 | 0.971785 |
| Slc9a1 | 4.82007 | 10.6794 |
| Phactr4 | 4.334712368 | 9.60133 |
| Cdh3 | 6.8714949 | 15.220011 |
| Abca3 | 1.59293 | 3.524957 |
| Nhsl1 | 1.5876419 | 3.51315667 |
| Lime1 | 4.85813 | 10.7404 |
| Lrp4 | 0.8135286 | 1.7979379 |
| Srek1 | 4.45921 | 9.85297 |
| Mast4 | 1.524514 | 3.368038 |
| Hipk2 | 1.072797987 | 2.369542 |
| Kremen1 | 3.1895609 | 7.043113 |
| Lhx9 | 19.105482 | 42.18212 |
| Ulk3 | 1.20209 | 2.65307 |
| Dennd4b | 1.97235 | 4.35294 |
| Amotl2 | 6.9898 | 15.42567 |
| Nf1 | 3.215859 | 7.086698 |
| Plxna1 | 6.63122 | 14.6102 |
| Trim8 | 9.33041 | 20.548 |

| | | |
|---|---|---|
| Gabbr1 | 4.404427429 | 9.69814086 |
| Me3 | 0.514192 | 1.13204 |
| Arhgef2 | 8.646879 | 19.025944 |
| Crtc2 | 6.42298 | 14.1308 |
| Igfbp3 | 105.259 | 231.551 |
| Fam193b | 11.292998 | 24.8362 |
| Frem1 | 2.68619303 | 5.902513526 |
| Sbf1 | 8.1964631 | 18.00671149 |
| B4galnt4 | 7.30045 | 16.0367 |
| Eps15l1 | 9.007437 | 19.780856 |
| Flt4 | 1.05699 | 2.31958 |
| Pml | 6.44209324 | 14.11098 |
| Trim25 | 9.03185 | 19.77257 |
| Fryl | 1.08621 | 2.37783 |
| Brpf3 | 5.54496386 | 12.1260591 |
| Map7 | 1.93641114 | 4.234102 |
| Arhgef18 | 3.35035 | 7.31819 |
| Tifa | 1.56583501 | 3.41991 |
| Zscan25 | 2.94256992 | 6.425706 |
| Cnksr1 | 0.559006 | 1.22015 |
| Sema3g | 1.018110044 | 2.219353817 |
| Itga3 | 6.67535956 | 14.54800068 |
| Stat2 | 2.64501 | 5.76312 |
| Ccl3 | 0.568699 | 1.23893 |
| Smg6 | 6.1265 | 13.33539 |
| Phf2 | 5.66223 | 12.30963 |
| Celsr1 | 2.240102649 | 4.86976 |
| Cdk18 | 0.573143 | 1.24524 |
| Trpc7 | 1.382621134 | 3.002959907 |
| Lmo3 | 0.53007617 | 1.151079727 |
| Atp1b2 | 13.43 | 29.1631 |
| Arsb | 2.58742 | 5.61419 |
| Clcn6 | 2.618479 | 5.68049196 |
| Ttc38 | 5.497392323 | 11.91808 |
| Fasn | 12.51185621 | 27.08906435 |
| Rnf169 | 2.779493 | 6.01684 |
| Pprc1 | 7.4260928 | 16.07280787 |
| Taf15 | 14.619 | 31.6365 |
| Mertk | 1.19347 | 2.58265 |
| Smtn | 5.30119839 | 11.46797986 |
| Mdm4 | 7.80168371 | 16.869104 |
| Caskin1 | 0.831396 | 1.79631 |
| Ncstn | 13.4297 | 29.0112 |
| Elmsan1 | 2.027639 | 4.378766 |
| Nfat5 | 4.500146639 | 9.716032 |
| Cc2d1a | 4.342424 | 9.36242 |
| Ecm1 | 37.26146244 | 80.28984 |

| | | |
|---|---|---|
| Pitpnm1 | 1.05107 | 2.264258 |
| Rsbn1 | 0.817284 | 1.76047 |
| Phf21a | 4.4655811 | 9.617641756 |
| BC065397 | 0.608897 | 1.31067 |
| Sobp | 0.770485697 | 1.656041 |
| Kalrn | 2.0802468 | 4.4680395 |
| Zkscan2 | 0.707438 | 1.519194 |
| Myrf | 23.5353984 | 50.474667 |
| Usp49 | 1.934689 | 4.148222 |
| Dnaaf5 | 3.49561 | 7.49493 |
| Tmem132a | 21.3499 | 45.7473 |
| Alms1 | 0.90401864 | 1.93700028 |
| Erbb2 | 8.49925 | 18.1883 |
| Atp10d | 1.99007735 | 4.25848 |
| Ano8 | 3.32977 | 7.1184 |
| Triobp | 7.356345 | 15.71732 |
| Dpp6 | 0.4527987 | 0.966534286 |
| Tgfb1 | 5.95289 | 12.7058 |
| Zfp839 | 2.034921178 | 4.341277 |
| Rab36 | 2.08786377 | 4.45373173 |
| Xlr3a | 0.828775 | 1.76785 |
| Fus | 80.8975 | 172.554 |
| Tshz2 | 7.93286 | 16.91534 |
| Mss51 | 0.68744234 | 1.465456173 |
| Zfp57 | 8.6156599 | 18.36512671 |
| Dzip1 | 3.43894046 | 7.330285 |
| Supt6 | 8.72185 | 18.5847 |
| Nfkbiz | 0.927796441 | 1.976441294 |
| Unc13b | 1.02982346 | 2.193273735 |
| Lsamp | 0.7009104 | 1.49204 |
| Gng7 | 1.260366646 | 2.681086031 |
| Polm | 2.614290847 | 5.55805 |
| Phc3 | 1.069500793 | 2.272620143 |
| Ifi27l2a | 30.3616 | 64.512349 |
| Agrn | 41.0908215 | 87.2920297 |
| Igfbp5 | 81.5537 | 173.2298 |
| Rnf157 | 3.43961 | 7.30406 |
| Fgd1 | 7.927307 | 16.8307803 |
| Plekhm1 | 2.95801 | 6.27076 |
| Trp53 | 44.1934 | 93.6741 |
| Hoxc5 | 8.84524 | 18.7299 |
| Prrc2c | 7.84125 | 16.59976 |
| Sirt6 | 8.849621 | 18.7316 |
| Dock5 | 0.463117 | 0.980098 |
| Cadps2 | 1.652106197 | 3.494922934 |
| Fnbp1 | 3.646848774 | 7.71379172 |
| Acsf2 | 7.02493483 | 14.85374 |

| | | |
|---|---|---|
| Tnpo2 | 9.7508748 | 20.610552 |
| Gm3002 | 0.468974 | 0.991115 |
| Ube4b | 13.216198 | 27.91758019 |
| Rprm | 8.13967 | 17.186 |
| Nphp3 | 1.5188424 | 3.206592 |
| Ccl6 | 2.01387 | 4.2505 |
| Lamc1 | 12.1505992 | 25.6379344 |
| Adam11 | 1.32878039 | 2.80159 |
| Asah2 | 0.540694689 | 1.139589975 |
| Pou6f1 | 1.687212361 | 3.553049893 |
| Midn | 24.6155 | 51.8322 |
| Wnk2 | 0.734473336 | 1.546472538 |
| Slc15a3 | 0.737694 | 1.55324 |
| Adcy6 | 7.522541123 | 15.82928296 |
| Atg2b | 1.68777329 | 3.54946235 |
| Kcnb1 | 0.870422 | 1.830413 |
| Zfp276 | 1.996 | 4.19579 |
| Map4k2 | 3.286209627 | 6.904742696 |
| Pcgf2 | 13.0024 | 27.29852244 |
| Grik5 | 11.271374 | 23.660206 |
| Gm1821 | 556.8008 | 1168.749 |
| Epb4.1l1 | 6.421551067 | 13.47509489 |
| Esm1 | 0.513326 | 1.07573 |
| Aplp1 | 11.1264 | 23.3154 |
| Cdc42bpb | 11.1545 | 23.367 |
| Tmem2 | 4.196523721 | 8.788614465 |
| Zfp219 | 13.1374393 | 27.511137 |
| Col23a1 | 1.829861926 | 3.830653 |
| Ntsr1 | 1.52801 | 3.19859 |
| Arhgef12 | 3.68142629 | 7.70598944 |
| Nfrkb | 3.42845862 | 7.174624167 |
| Kif21b | 1.436436084 | 3.0056931 |
| Gnpda1 | 5.34445 | 11.1778 |
| Ccdc114 | 1.389361 | 2.905052 |
| Spry1 | 10.87084 | 22.72892083 |
| Tnfrsf18 | 0.8988799 | 1.87937564 |
| Unk | 7.078705725 | 14.793526 |
| Zfp608 | 1.782622 | 3.722827 |
| Atp10a | 1.643376 | 3.43181552 |
| Tpcn1 | 7.369979788 | 15.385658 |
| Parp12 | 7.947366 | 16.56433 |
| Atxn2 | 5.545072 | 11.529145 |
| Fstl3 | 0.551622 | 1.14669 |
| Col11a1 | 1.860918 | 3.867531689 |
| Bcorl1 | 2.1613009 | 4.48875266 |
| Col4a6 | 5.058372 | 10.50499 |
| Whrn | 1.9413004 | 4.030144504 |

| | | |
|---|---|---|
| Cep104 | 2.71921 | 5.64421 |
| Lmln | 0.5262735 | 1.092259585 |
| Nktr | 4.4806142 | 9.2968981 |
| Megf8 | 6.647935 | 13.7919 |
| Smarcc2 | 11.30462377 | 23.441809 |
| Spata2l | 0.45971 | 0.952238 |
| Pdzrn3 | 2.707384674 | 5.607274021 |
| Sox12 | 18.9807 | 39.2719 |
| Hspb6 | 2.134946282 | 4.416949 |
| Tctn1 | 3.826824 | 7.91687 |
| Pigg | 0.719581 | 1.488549 |
| Kdm2a | 8.18894609 | 16.93546306 |
| Hoxc4 | 5.2349987 | 10.825965 |
| Hcn2 | 0.903909 | 1.86788 |
| Efnb2 | 7.95229 | 16.4318 |
| Plekhh2 | 2.52511834 | 5.217381 |
| Neat1 | 2.497877 | 5.160027 |
| Smg1 | 2.80430823 | 5.792001 |
| Zbed6 | 1.64289 | 3.39179 |
| Wbscr17 | 1.09504 | 2.25992 |
| Eif4ebp2 | 16.9542 | 34.9757 |
| Stxbp5 | 1.17635083 | 2.42663 |
| Psd | 0.78736 | 1.62384 |
| Plekha4 | 1.834108935 | 3.780824219 |
| Usp2 | 1.64103454 | 3.380103419 |
| Nt5dc3 | 1.36692419 | 2.815278 |
| Plec | 4.5109643 | 9.29019463 |
| Lilrb4a | 0.544071 | 1.119923 |
| Plaur | 0.796915 | 1.63977 |
| Katnal1 | 1.044759618 | 2.14765997 |
| Ncdn | 5.82769 | 11.97176 |
| Ptpn21 | 1.371849976 | 2.817369 |
| Mir143hg | 1.02367 | 2.09927 |
| Gm21119 | 5.177801 | 10.59999 |
| Nat6 | 1.4556 | 2.97945 |
| Acvr1b | 4.47534 | 9.16051 |
| Eps8l2 | 1.027261794 | 2.101719 |
| Ebf4 | 5.367940816 | 10.97885 |
| Asxl3 | 0.890683 | 1.8213343 |
| Adamts7 | 3.90088887 | 7.976054 |
| Gpbp1l1 | 5.622756 | 11.49613 |
| Clasp1 | 3.91283343 | 7.993158235 |
| Rfx1 | 5.140065 | 10.49517 |
| Kcne4 | 1.79324 | 3.65997 |
| Fos | 0.661774 | 1.35064 |
| Mapk1ip1 | 3.502279 | 7.147534 |
| Yeats2 | 7.165772 | 14.6218447 |

| | | |
|---|---|---|
| Troap | 1.947579938 | 3.97391 |
| Pfkfb3 | 2.704877732 | 5.5177708 |
| Npr2 | 4.665191 | 9.51609 |
| Tacstd2 | 2.67495 | 5.45604 |
| Kat6a | 3.6860708 | 7.51658071 |
| Tshz3 | 2.43378 | 4.96111 |
| Sema3f | 9.10299 | 18.53856 |
| Gsk3a | 14.4177 | 29.3523 |
| Lad1 | 0.938964 | 1.91095 |
| Mapk8ip3 | 7.4017764 | 15.06072099 |
| Cep162 | 0.598255 | 1.21671042 |
| Cep85l | 0.500797561 | 1.018087 |
| Ccl4 | 0.809647 | 1.64459 |
| Poli | 2.794460984 | 5.675819102 |
| Tescl | 1.67784 | 3.40736 |
| Zmat3 | 1.8655156 | 3.78822 |
| Usp36 | 3.2803823 | 6.661172249 |
| Mospd3 | 7.877505 | 15.99386 |
| BC021891 | 1.030861 | 2.09276 |
| Med24 | 12.38664374 | 25.14076 |
| Gm3893 | 1.5094 | 3.06214 |
| Cacng4 | 4.032270106 | 8.176553333 |
| Sfxn5 | 1.105621857 | 2.240011 |
| Ankle1 | 1.85929 | 3.763666 |
| Iqce | 3.065361327 | 6.203153 |
| Efr3b | 1.838730514 | 3.71997816 |
| Tmc8 | 0.835345552 | 1.689862941 |
| Mpp2 | 1.867541 | 3.775855001 |
| Wdr76 | 2.67900145 | 5.415034871 |
| Rnf150 | 0.872955 | 1.764127 |
| Abl1 | 15.61346485 | 31.543833 |
| Gatad2a | 12.30428964 | 24.834147 |
| Patl1 | 5.982741 | 12.07413 |
| Taf3 | 1.55531 | 3.1353 |
| Specc1 | 1.150014123 | 2.318140464 |
| Top3a | 2.74783 | 5.52868 |
| Dnmt3a | 11.56363043 | 23.26228204 |
| Sik1 | 5.85617 | 11.77629888 |
| Bag6 | 41.05577 | 82.51907668 |
| Braf | 1.3573423 | 2.725092124 |
| Nfic | 7.610285922 | 15.27683 |
| Pitpnc1 | 2.044384858 | 4.103718 |
| Reck | 4.717 | 9.46364 |
| Ndst1 | 7.8383895 | 15.71631 |
| Ap2a1 | 19.1472 | 38.3812 |
| Sestd1 | 3.96985881 | 7.95533 |
| Zc3h7b | 13.39967 | 26.84558 |

| | | |
|---|---|---|
| Rap1gap2 | 0.80392384 | 1.610282706 |
| Igf1r | 4.129176 | 8.26946 |
| Slc9a5 | 1.954578652 | 3.912556 |
| Hgs | 10.6708 | 21.35645 |
| Slc2a10 | 0.72291 | 1.44616 |
| Hoxc6 | 11.7892 | 23.5489 |
| Zc3h4 | 7.111435 | 14.19998 |
| Brd2 | 24.68418736 | 49.267 |
| Zfp516 | 6.63464 | 13.21735 |
| Dchs1 | 5.97146 | 11.89501 |
| Pcyox1l | 3.905342216 | 7.77858 |
| Adgrb2 | 8.07139067 | 16.072278 |
| Rgs1 | 6.34848 | 12.63784 |
| Loxl2 | 6.0565175 | 12.0397275 |
| Nlrp5-ps | 1.015185601 | 2.017639216 |
| Mcc | 2.008229364 | 3.990533 |
| Usp11 | 11.7224 | 23.29 |
| Tgm5 | 0.708717 | 1.40739 |
| Dpf3 | 2.613624 | 5.182299 |
| Cux1 | 9.988718435 | 19.7989781 |
| Fgf18 | 0.71489561 | 1.416710629 |
| Sec24b | 6.55997128 | 12.998882 |
| Slc2a4rg-ps | 1.122574 | 2.224193939 |
| Cbln2 | 0.853942801 | 1.691869 |
| Ppp1r9a | 3.006053965 | 5.954584368 |
| Slc7a1 | 1.4322633 | 2.835491069 |
| Zbtb34 | 2.20610096 | 4.364395 |
| Myo18a | 3.629036302 | 7.17699788 |
| Mmp2 | 55.2696 | 109.225 |
| Col2a1 | 0.504634236 | 0.996907 |
| Rtkn | 5.904335 | 11.65968 |
| Ppp1r9b | 18.0874 | 35.71758 |
| Phlpp1 | 1.81676 | 3.58533 |
| Epha1 | 0.958153 | 1.89048 |
| Fbxl19 | 22.23906 | 43.87526 |
| F2rl1 | 0.588713 | 1.1611 |
| Sgpp2 | 0.679041 | 1.3391 |
| Trim21 | 1.98414977 | 3.9111981 |
| Pvrl1 | 2.905767 | 5.72619 |
| Timeless | 7.198971 | 14.18549491 |
| Chd2 | 4.451973802 | 8.76995 |
| Ak5 | 0.78149 | 1.53917 |
| Sema4a | 1.675771343 | 3.298669832 |
| Gstm3 | 5.82486 | 11.4556 |
| Negr1 | 1.10759 | 2.17816 |
| Nop2 | 10.3768 | 20.3991 |
| Adamts20 | 0.49518 | 0.972906 |

| | | |
|---|---|---|
| Htt | 1.93672 | 3.8049 |
| Rhbdf1 | 8.567098 | 16.8250388 |
| Atg2a | 4.63541 | 9.09644 |
| Itga1 | 1.304439 | 2.559781 |
| Polh | 2.639558822 | 5.179258 |
| Taf1a | 3.057098073 | 5.996924037 |
| Gsk3b | 5.91528 | 11.60268 |
| Kank2 | 8.80138 | 17.2526 |
| Mta1 | 30.6927 | 60.1265 |
| Pfas | 4.7719284 | 9.344535 |
| Adamts14 | 0.805887 | 1.57769 |
| Svil | 3.526826705 | 6.902765 |
| Ccl12 | 1.15152 | 2.25301 |
| Lmtk2 | 2.24586412 | 4.393177 |
| Agbl5 | 6.453981 | 12.6208456 |
| Dab2ip | 9.211950828 | 17.98810272 |
| Fv1 | 1.14227527 | 2.229388 |
| Zfp777 | 6.467040582 | 12.615117 |
| Bmp4 | 3.271203634 | 6.38059 |
| Spef1 | 4.08243 | 7.96083 |
| Firre | 4.77042299 | 9.30124895 |
| Rere | 13.098542 | 25.53413 |
| Lgals4 | 0.832579 | 1.62257 |
| Flrt1 | 0.753401 | 1.46745 |
| Sox5 | 0.888807556 | 1.731168254 |
| Mal2 | 1.07738 | 2.09768 |
| Gmip | 1.716719 | 3.341961 |
| Bcl11a | 1.66308662 | 3.2367608 |
| Fnbp4 | 5.197371 | 10.11401 |
| Mysm1 | 1.434024159 | 2.79044 |
| Rdh5 | 0.662055 | 1.28825 |
| Cacnb1 | 2.97675296 | 5.791072517 |
| Rin1 | 0.52493101 | 1.020984625 |
| Malt1 | 1.358207 | 2.641351 |
| Fam160b1 | 4.8220981 | 9.37656 |
| C77080 | 5.212171158 | 10.13139327 |
| Parvg | 0.981042792 | 1.905626416 |
| Ski | 17.766287 | 34.50974 |
| Naalad2 | 1.338985536 | 2.600712391 |
| Sh2b2 | 1.8317236 | 3.5567037 |
| Lrrc16b | 1.6494753 | 3.202239061 |
| Trpv4 | 1.11314655 | 2.160540374 |
| Gli2 | 2.92702 | 5.67768 |
| Col4a3 | 12.2764001 | 23.8004996 |
| Osbpl7 | 1.84433243 | 3.573661 |
| Sema4f | 0.7093986 | 1.374540691 |
| Smarcc1 | 20.5107 | 39.74119 |

| | | |
|---|---|---|
| Gas7 | 2.5987441 | 5.0346225 |
| Ubap2l | 22.2230846 | 43.0494076 |
| Wdtc1 | 11.47845589 | 22.221876 |
| Usp19 | 18.7007728 | 36.18748454 |
| Arhgef10 | 2.8060696 | 5.429344905 |
| Cdk12 | 3.3499707 | 6.479479 |
| Cplx2 | 2.15696861 | 4.171926 |
| Rdh13 | 4.10620028 | 7.939939 |
| Stk35 | 5.72893 | 11.07254 |
| Dlg5 | 8.61959 | 16.65429 |
| Proser1 | 4.18148 | 8.07655 |
| Lrrc46 | 0.596978 | 1.15306 |
| Wrn | 0.98573723 | 1.903535066 |
| Arrdc2 | 1.74353 | 3.36686 |
| Tgm3 | 1.384519339 | 2.673082487 |
| Bace1 | 4.822792 | 9.310378709 |
| Tnks1bp1 | 8.95159673 | 17.27932 |
| Hdac10 | 2.271988551 | 4.383275844 |
| Piezo1 | 7.61651 | 14.6848 |
| Fbf1 | 4.847104 | 9.33455104 |
| Scyl2 | 6.55093 | 12.61019 |
| Slc41a3 | 1.760822516 | 3.388993 |
| Skiv2l | 7.58154 | 14.59088747 |
| Dync1h1 | 22.89981 | 44.05281 |
| Mcf2l | 1.170142813 | 2.250766324 |
| Sptb | 0.9520841 | 1.831096 |
| Zfp532 | 5.714956 | 10.9885362 |
| Gfpt2 | 1.110470918 | 2.134799 |
| Unc5b | 5.1447554 | 9.889095 |
| Tanc1 | 3.662384456 | 7.03767376 |
| Rbak | 2.45256551 | 4.711704 |
| Map9 | 0.78353876 | 1.50515208 |
| Pfkl | 29.8119 | 57.264 |
| Tbc1d16 | 6.13481492 | 11.78276199 |
| Zfp691 | 2.3001073 | 4.414798 |
| Rnf38 | 6.699434309 | 12.858529 |
| Efna5 | 2.81681926 | 5.40612 |
| Kif7 | 2.7048835 | 5.190404008 |
| BC017158 | 6.97873 | 13.38371 |
| Tmem184b | 15.89948453 | 30.47490301 |
| Neurl1b | 0.732064 | 1.40308 |
| Zfp939 | 0.554993894 | 1.063662298 |
| Megf6 | 2.497767 | 4.786531 |
| Fbn2 | 6.12709 | 11.73265 |
| Gm608 | 1.158334361 | 2.21761443 |
| Slc25a27 | 1.916572 | 3.667791 |
| Pcid2 | 9.07528 | 17.3624 |

| | | |
|---|---|---|
| Dnm2 | 10.65063257 | 20.372743 |
| Srrm1 | 23.15739 | 44.28814 |
| Cfp | 2.279931 | 4.360321 |
| Cpsf7 | 16.2395483 | 31.05386 |
| Ppp6r1 | 19.215 | 36.7426 |
| Chst2 | 1.616 | 3.08949 |
| Tet3 | 3.94365694 | 7.538612384 |
| Hk2 | 4.69854 | 8.98119 |
| Sema4c | 7.74854282 | 14.807196 |
| Mark2 | 11.03010409 | 21.05672863 |
| Lag3 | 3.285452 | 6.27024 |
| Ewsr1 | 61.2989295 | 116.98155 |
| Scarf2 | 16.5908 | 31.655 |
| Gtf2ird2 | 2.469057 | 4.71062 |
| P3h3 | 12.55916531 | 23.948084 |
| Fxyd7 | 0.982001 | 1.87229 |
| Ccdc8 | 15.78496445 | 30.08001039 |
| Simc1 | 2.01763 | 3.84467 |
| Ngfr | 6.49319 | 12.3706 |
| Slc7a5 | 13.316 | 25.3599 |
| Gfra2 | 1.572448951 | 2.994556307 |
| Prdm11 | 3.017547711 | 5.74523598 |
| Stxbp4 | 1.8015 | 3.42846 |
| Wfikkn2 | 0.9751501 | 1.855162 |
| Trim46 | 1.151095486 | 2.187715 |
| Mphosph9 | 1.11008539 | 2.1095218 |
| Wfs1 | 4.10621 | 7.80293 |
| Zfp236 | 2.098999 | 3.988297917 |
| Orai2 | 2.198527577 | 4.175332 |
| Tmem194b | 2.57440524 | 4.8870309 |
| Crispld1 | 0.90613275 | 1.719515808 |
| Lrrc45 | 7.734826 | 14.67449 |
| Nuak1 | 4.744144 | 9.00055 |
| Ccdc142 | 1.82873 | 3.46891 |
| Ylpm1 | 4.51544 | 8.55746 |
| Fchsd1 | 1.99286 | 3.77634 |
| Ccdc24 | 2.31531 | 4.38641 |
| Sart1 | 8.1834469 | 15.5011 |
| Fgfr4 | 0.638945 | 1.20997 |
| Clk4 | 9.65941 | 18.28919 |
| Gpatch4 | 4.956752 | 9.38375 |
| Sntb2 | 2.69563 | 5.10114 |
| Spred2 | 6.93505 | 13.121577 |
| Asap3 | 1.352069228 | 2.55736 |
| Sh3pxd2a | 6.55707765 | 12.40222334 |
| Magef1 | 2.39762 | 4.53448 |
| Atp6v0a1 | 6.001818972 | 11.34944 |

| | | |
|---|---|---|
| Wnt5a | 15.31576 | 28.96191597 |
| Slc6a9 | 4.25091 | 8.03668 |
| Pxylp1 | 2.303414375 | 4.351253774 |
| Col22a1 | 0.558597 | 1.05521 |
| Dnmt1 | 16.50780887 | 31.18232333 |
| Sash1 | 3.749623823 | 7.082426 |
| Pfkfb2 | 1.623758718 | 3.066764153 |
| Dclk2 | 5.776414952 | 10.90806901 |
| Bend4 | 1.560552377 | 2.9460516 |
| Zfp362 | 21.86163 | 41.26926 |
| Dopey2 | 1.733756327 | 3.271235584 |
| Dmpk | 6.359786722 | 11.99004415 |
| Pax8 | 4.80380637 | 9.054550451 |
| Phlpp2 | 1.4995676 | 2.825014 |
| Adam19 | 6.7819516 | 12.77554108 |
| Sh2d3c | 6.31293 | 11.891672 |
| Ipo9 | 16.3371 | 30.7734 |
| Tcaf1 | 7.872404923 | 14.8119489 |
| Rnf123 | 3.567425 | 6.71101 |
| Pdxk | 2.62933 | 4.94617 |
| Grik3 | 0.635799 | 1.19593 |
| Zfp653 | 3.44728 | 6.4805 |
| Myo1e | 3.393469 | 6.37905 |
| Lpcat4 | 4.09395 | 7.69499 |
| Fbxl17 | 4.149493 | 7.791361 |
| Madd | 1.350822018 | 2.536093438 |
| Mon1b | 1.866312 | 3.503896 |
| Dennd1a | 4.00269 | 7.51162 |
| Atxn7l2 | 3.891502 | 7.302205 |
| E4f1 | 4.38556789 | 8.228638 |
| Kcnn2 | 1.571441 | 2.94789 |
| Slc8b1 | 2.175121465 | 4.079362218 |
| Rac2 | 2.00451 | 3.7575 |
| Csdc2 | 11.6269 | 21.7835 |
| Myh8 | 0.622948 | 1.16692 |
| Etv1 | 1.23548167 | 2.313927116 |
| Dpyd | 0.848272 | 1.58867 |
| Tcte2 | 0.619506 | 1.159768 |
| Tceanc2 | 1.63055 | 3.0524 |
| Nxf1 | 24.29614 | 45.4814 |
| Ctif | 1.723719 | 3.22600083 |
| Ankrd23 | 1.884486 | 3.5248376 |
| Mpp7 | 0.8875342 | 1.659935 |
| Klhdc4 | 8.81422792 | 16.48232 |
| Smarca2 | 16.32901867 | 30.51889 |
| D130017N( | 0.669159 | 1.25058 |
| Rhot2 | 6.18926 | 11.5662 |

| | | |
|---|---|---|
| Gga3 | 7.09758 | 13.26021 |
| Tead4 | 0.637862502 | 1.191269781 |
| Prickle1 | 9.970396 | 18.61534 |
| Irgq | 3.73637 | 6.9724 |
| Fbxo42 | 5.92957 | 11.05663 |
| Wasf3 | 2.192730221 | 4.08813175 |
| Rptor | 4.70623 | 8.766394 |
| Dhdh | 0.780424 | 1.45368 |
| Ambra1 | 6.58516 | 12.25984 |
| Gramd1a | 14.678324 | 27.31635668 |
| Zfp462 | 3.31417477 | 6.165666 |
| Anpep | 5.62372 | 10.4621 |
| Ano1 | 9.553612 | 17.77036937 |
| Ago1 | 6.543811266 | 12.17056157 |
| Axin2 | 8.0724623 | 15.00834 |
| Ccnt2 | 4.97264 | 9.24146 |
| Kirrel | 8.6248 | 16.0195451 |
| Mst1r | 1.93531652 | 3.59434543 |
| Dnm3os | 1.656488 | 3.076382 |
| Sec31a | 23.84324874 | 44.27152 |
| Spats2 | 15.317911 | 28.42923 |
| Kcnt1 | 3.455446587 | 6.4107231 |
| Espl1 | 6.04208904 | 11.20732 |
| Gramd4 | 7.928982257 | 14.70505552 |
| Doc2b | 7.908903611 | 14.66489 |
| Erf | 13.45344883 | 24.94401997 |
| Gulo | 0.561357 | 1.04074 |
| Gm15284 | 0.558076 | 1.03452 |
| Hoxc9 | 3.30865 | 6.1327 |
| Mpp3 | 1.088869612 | 2.017570941 |
| Sbno2 | 3.248839 | 6.0172094 |
| Nrsn2 | 1.5023297 | 2.782171 |
| Lama1 | 3.5038394 | 6.486969521 |
| Wdr81 | 2.493910718 | 4.616321 |
| Esrp1 | 3.46015745 | 6.401674 |
| Plxna3 | 3.524193 | 6.51938 |
| Kdr | 6.3867 | 11.8142 |
| Mnt | 4.787169 | 8.854278348 |
| Card10 | 2.022603466 | 3.739152421 |
| Elf4 | 2.993891 | 5.53316738 |
| Arl4c | 8.67937 | 16.0371 |
| Tnrc6b | 1.95901727 | 3.619468 |
| Stc1 | 1.34861 | 2.49007 |
| Cpsf1 | 14.312417 | 26.42438 |
| Tsc2 | 9.346398 | 17.25039824 |
| Tnfrsf1b | 0.880133 | 1.62435 |
| Aplp2 | 25.651695 | 47.340392 |

| | | |
|---|---|---|
| Rab11fip3 | 5.4067 | 9.97612836 |
| Fbxl16 | 1.24966 | 2.30527 |
| Stx1a | 6.41114 | 11.81854 |
| Zfyve26 | 2.54769 | 4.69583 |
| Mfn2 | 6.2524688 | 11.52310784 |
| Kremen2 | 0.75062486 | 1.38324 |
| Kif5c | 6.412821 | 11.813501 |
| Tle3 | 9.879454329 | 18.19585981 |
| Map1a | 1.484902828 | 2.734745 |
| Fat4 | 2.877994 | 5.30006 |
| Slc25a22 | 6.88933 | 12.67511 |
| C2cd5 | 4.04064019 | 7.43316949 |
| Ubash3b | 1.186043725 | 2.181455232 |
| Trio | 8.476201 | 15.589631 |
| Col4a5 | 6.506332152 | 11.96594708 |
| Trp53bp2 | 5.14442 | 9.45957 |
| Abi3 | 1.09745 | 2.017971 |
| Elmo1 | 1.743243 | 3.20509 |
| Amigo1 | 1.0140136 | 1.864232 |
| Dusp18 | 1.047982 | 1.925835 |
| Pvr | 2.3245981 | 4.271512052 |
| Setd7 | 2.53423 | 4.65609 |
| Dtx3 | 25.335792 | 46.544126 |
| Hdac4 | 2.342645247 | 4.3031056 |
| Pla2g6 | 5.3243574 | 9.778116 |
| Iqsec1 | 3.503365833 | 6.431599772 |
| Lrrc1 | 4.25325 | 7.80786 |
| Ighmbp2 | 1.44901 | 2.65918 |
| Gab2 | 3.92728 | 7.20689 |
| Adamts10 | 12.4146707 | 22.7636861 |
| Pik3c2b | 0.8588664 | 1.574409113 |
| Tirap | 1.50070088 | 2.749983769 |
| Dnal1 | 1.684341401 | 3.085505 |
| Icosl | 0.823705 | 1.508776 |
| Znrf3 | 1.603253339 | 2.936479953 |
| Magi1 | 4.5065462 | 8.2485718 |
| Ksr1 | 2.12428 | 3.8862 |
| Sema3b | 1.003151625 | 1.834978536 |
| Urb2 | 2.59697424 | 4.75035 |
| Shank3 | 2.2858081 | 4.181054 |
| Fbxo10 | 4.17439 | 7.63391 |
| Baz2a | 6.140681 | 11.22904 |
| Arap3 | 4.0216864 | 7.351859 |
| Depdc5 | 2.592453224 | 4.73692525 |
| Car12 | 1.328238181 | 2.42666395 |
| Heatr6 | 2.502381 | 4.5717197 |
| Huwe1 | 13.56113797 | 24.7725654 |

| | | |
|---|---|---|
| Git2 | 4.3112169 | 7.8740433 |
| Rcbtb1 | 9.81106 | 17.91667 |
| Trim62 | 6.74513 | 12.3144 |
| Cers6 | 2.653478475 | 4.84391 |
| Rbfox2 | 13.372522 | 24.4058228 |
| Tcf20 | 4.49408949 | 8.20145 |
| Kbtbd11 | 0.836196 | 1.526 |
| Ap5z1 | 2.239994 | 4.087317 |
| Prob1 | 0.804667 | 1.46791 |
| Mtx3 | 1.937518925 | 3.531916 |
| Inppl1 | 17.186508 | 31.32538 |
| Tbx2 | 27.10746 | 49.407 |
| Enthd2 | 3.784997 | 6.89823 |
| Zscan26 | 5.37929015 | 9.802907 |
| Ric1 | 3.31121 | 6.03357879 |
| Itsn1 | 8.258161 | 15.045359 |
| Zfp316 | 2.51062 | 4.573915859 |
| Tenm3 | 4.626676085 | 8.428046515 |
| Itga5 | 7.1132 | 12.9532 |
| Sema4d | 3.045268843 | 5.545023086 |
| Endov | 2.9474383 | 5.366322 |
| Ptafr | 0.42507 | 0.77380687 |
| Enc1 | 6.98027 | 12.7064 |
| Bcl3 | 1.076478778 | 1.959471 |
| Mob3c | 1.47558 | 2.68551 |
| Iqsec2 | 1.196729722 | 2.177188805 |
| Apc2 | 0.587461 | 1.068636 |
| Dip2a | 3.69361 | 6.7181 |
| Tgfa | 1.438 | 2.61429 |
| Lrrc4b | 1.69177 | 3.07558 |
| Pip4k2b | 8.72147 | 15.8434 |
| Hcls1 | 1.48 | 2.68835 |
| Rgl3 | 2.33373 | 4.237607 |
| Piezo2 | 0.83523062 | 1.515974828 |
| Leo1 | 6.11201 | 11.0935 |
| Mroh1 | 4.71249815 | 8.5531346 |
| Adcy3 | 3.2808503 | 5.95452255 |
| Dock1 | 5.79667 | 10.5163 |
| Insr | 2.81150745 | 5.098416 |
| Stk4 | 4.951464 | 8.978891 |
| Nid2 | 14.0671 | 25.5043 |
| Aqr | 6.068569 | 11.0013562 |
| Pcdhga4 | 0.535667 | 0.970713 |
| Dstyk | 3.24261 | 5.87483 |
| Fer1l5 | 2.0673792 | 3.7453348 |
| AI414108 | 0.548492 | 0.993509 |
| Rpap1 | 8.31553271 | 15.05873716 |

| | | |
|---|---|---|
| Gpr162 | 2.911616239 | 5.270795044 |
| Elac1 | 1.38119 | 2.49921 |
| Eif4g1 | 38.87286431 | 70.324538 |
| Bbs1 | 1.35946 | 2.45916 |
| Ddx46 | 8.534221 | 15.43386 |
| Cuedc1 | 4.49857 | 8.132852 |
| Pdcd6ip | 9.53279909 | 17.23287494 |
| Stxbp1 | 3.899394 | 7.047822 |
| Anks3 | 5.766228243 | 10.41855 |
| Lrp1 | 9.929824 | 17.9411 |
| Clec7a | 0.546963294 | 0.988065 |
| Hgf | 1.743429676 | 3.149401521 |
| Vsig10l | 1.14667 | 2.070458 |
| Aebp1 | 7.8904543 | 14.24015447 |
| Iffo2 | 2.667928 | 4.814331 |
| Gpatch1 | 1.94389 | 3.50778 |
| Mirg | 7.98541792 | 14.40913137 |
| Pard3 | 6.606696709 | 11.919809 |
| Ankzf1 | 4.181994 | 7.540827888 |
| Nbeal2 | 0.84480577 | 1.52291051 |
| Fosl2 | 3.12702 | 5.63687 |
| Pam | 18.99734 | 34.241294 |
| Ifi27 | 61.1543 | 110.22317 |
| Pak4 | 13.3626 | 24.0805 |
| Kif1c | 7.70085891 | 13.87170762 |
| Atp11a | 5.637780195 | 10.153985 |
| Dync1li2 | 10.8449 | 19.5233 |
| Ep300 | 4.667 | 8.40021 |
| Neurl4 | 8.2738519 | 14.89038301 |
| Slc12a4 | 6.68253941 | 12.02377983 |
| Bcor | 4.1633245 | 7.48930313 |
| Abcg4 | 0.825464 | 1.48462 |
| Peak1 | 3.20894 | 5.77055 |
| Zbtb7b | 1.81805304 | 3.268848677 |
| B130024G1 | 2.705727 | 4.86334 |
| Gm13375 | 5.106562435 | 9.17743 |
| Trim9 | 0.63990094 | 1.1497125 |
| Ppp1r12b | 1.205844 | 2.166316 |
| Igfn1 | 2.9400155 | 5.2811072 |
| Mttp | 0.69601553 | 1.249810373 |
| Mgat5 | 1.8853831 | 3.384750893 |
| Cdc37l1 | 4.815942 | 8.644039 |
| Zc3h18 | 9.91962585 | 17.803535 |
| Pdzd2 | 0.976118 | 1.751431 |
| Man1c1 | 1.840877 | 3.301972285 |
| Dopey1 | 0.645391232 | 1.157407 |
| Nol4l | 6.144643327 | 11.0192761 |

| | | |
|---|---|---|
| Armc5 | 4.56965 | 8.19334 |
| Fto | 16.4678 | 29.5182 |
| C3ar1 | 1.34559 | 2.41116 |
| Aspg | 0.89445794 | 1.602710507 |
| Tsc1 | 3.2185669 | 5.765302896 |
| Mllt1 | 16.79331516 | 30.07470048 |
| Nckap1l | 1.04452 | 1.87057 |
| Zyg11a | 1.523979 | 2.72873 |
| Rassf2 | 1.521893754 | 2.724761421 |
| Baz2b | 2.5323758 | 4.532640515 |
| AU040320 | 3.44057 | 6.1571788 |
| Wwc1 | 2.2718 | 4.065348679 |
| Six5 | 16.9615 | 30.33856 |
| Ppef2 | 0.931245 | 1.665428 |
| Pelp1 | 15.5379 | 27.7851 |
| Znfx1 | 3.431925 | 6.13413 |
| Gm19757 | 0.643079 | 1.14942 |
| Itfg2 | 5.33267 | 9.53059 |
| Parp8 | 2.973388 | 5.31254 |
| Rbl2 | 2.066350404 | 3.689066109 |
| Kdm7a | 1.28282 | 2.28871 |
| Scrib | 12.0462526 | 21.49132289 |
| Fam217b | 0.621118 | 1.10798 |
| Mical3 | 2.8723576 | 5.12360143 |
| Med23 | 6.492242 | 11.57942 |
| Ice1 | 5.948984 | 10.608446 |
| Gdi1 | 22.74197 | 40.5495 |
| Tmem104 | 1.5954985 | 2.84353 |
| C530008M | 1.3650815 | 2.432126476 |
| Ift57 | 3.77946623 | 6.73322 |
| Ptprv | 5.905035855 | 10.514991 |
| Rerg | 2.211831493 | 3.938521811 |
| Ino80d | 1.31423763 | 2.33722165 |
| Dvl2 | 15.9909 | 28.4275 |
| Upk1b | 5.549363218 | 9.86521 |
| Syne1 | 0.995381416 | 1.7687057 |
| Gm20605 | 2.28913243 | 4.066758 |
| Palmd | 0.613034 | 1.08893 |
| Col6a1 | 22.6588 | 40.2349 |
| Fam168b | 29.35796 | 52.107072 |
| Bud13 | 6.27396 | 11.13441 |
| Camsap1 | 4.026849537 | 7.144377058 |
| Atxn3 | 1.872943731 | 3.32228858 |
| Nfkb1 | 5.58905 | 9.907192 |
| Gyltl1b | 3.17567365 | 5.628876895 |
| Lor | 7.948355 | 14.08626 |
| Zbtb7a | 1.789808004 | 3.171768858 |

| | | |
|---|---|---|
| Brpf1 | 4.45210724 | 7.888888 |
| Scnn1b | 0.989943163 | 1.753892581 |
| Mirlet7i | 0.969326 | 1.71697 |
| Dos | 5.07776 | 8.9912 |
| Coro2b | 2.68364 | 4.75174 |
| Dennd4a | 0.57066864 | 1.010431 |
| Nsf | 4.565341554 | 8.081522633 |
| Cpsf6 | 14.003768 | 24.78403 |
| Lrrc47 | 7.64235 | 13.524801 |
| Cldn9 | 3.80871 | 6.7385 |
| Lipg | 1.38725 | 2.45367 |
| Sik2 | 2.06939 | 3.66009 |
| Hirip3 | 8.10896 | 14.3397 |
| Sirt4 | 3.4089976 | 6.027834283 |
| Ncoa3 | 4.07391 | 7.20203 |
| Zfp446 | 1.373849711 | 2.42860843 |
| Zmym3 | 12.30535608 | 21.750792 |
| Fgf11 | 3.211221 | 5.67508 |
| Klf15 | 0.873121 | 1.54299 |
| Tfap4 | 7.04637 | 12.45 |
| Zfp74 | 0.9906879 | 1.750125 |
| Fam65a | 6.51934 | 11.51544 |
| Nxpe3 | 0.70507777 | 1.24441523 |
| Cep131 | 6.57051 | 11.5942 |
| Brat1 | 4.627104893 | 8.164315822 |
| Pik3cd | 3.284444013 | 5.794823 |
| Ninl | 2.251096 | 3.97124 |
| Csf1r | 3.95596 | 6.9788 |
| Kdm4a | 12.953354 | 22.836977 |
| Plekhm3 | 0.940336 | 1.65772 |
| Sel1l3 | 1.0745 | 1.89327 |
| Crtc1 | 6.64475 | 11.7069 |
| Bcl7a | 7.33425 | 12.92052 |
| E330009J0 | 2.0163899 | 3.552046 |
| Cog1 | 4.58372 | 8.07343 |
| Wdr19 | 1.801965 | 3.173595 |
| Gabpb2 | 3.45907362 | 6.09104358 |
| Atp2b1 | 9.28077 | 16.33506 |
| Ptch1 | 4.0844345 | 7.18685 |
| Dapp1 | 1.900772 | 3.343852087 |
| Nhlrc3 | 2.738653 | 4.81594225 |
| Prdm2 | 4.20386876 | 7.3905113 |
| Kazn | 7.4326281 | 13.06474193 |
| Anxa11 | 9.90797 | 17.41563443 |
| Chd4 | 32.6834 | 57.4381 |
| Bcar3 | 3.19722 | 5.6185 |
| Tgfbrap1 | 2.45311417 | 4.310583 |

| | | |
|---|---|---|
| Vamp2 | 11.9528 | 20.99902 |
| Pacs1 | 13.02843 | 22.88614 |
| Lpp | 1.739260306 | 3.054206007 |
| Pnpla2 | 5.078083607 | 8.914935 |
| Anks1 | 5.27698521 | 9.261338893 |
| Pcif1 | 12.0975 | 21.2266 |
| Slfn5 | 1.154723602 | 2.025632 |
| Ccnl1 | 9.915098 | 17.38488 |
| Tmem8b | 2.253475271 | 3.95094009 |
| Ppp6r2 | 3.6446064 | 6.3898637 |
| Jag1 | 2.24659 | 3.93779 |
| Rab2b | 3.11332 | 5.45652 |
| Fip1l1 | 11.79006 | 20.66278 |
| Ogdhl | 5.043756 | 8.837079237 |
| Kdm4c | 2.174815718 | 3.809292143 |
| Dffa | 6.40344255 | 11.2150109 |
| Tulp1 | 1.74025 | 3.04661 |
| Lipe | 2.730726 | 4.780583 |
| Dcaf5 | 5.779364 | 10.11728544 |
| E530011L2 | 1.13788 | 1.99163 |
| Fam219aos | 0.919023 | 1.60855 |
| Mdc1 | 4.8329742 | 8.45773 |
| Limk2 | 7.54313 | 13.1978 |
| Zbtb40 | 1.677778641 | 2.93288 |
| Tob2 | 4.56523 | 7.98001 |
| Ttc37 | 2.308691242 | 4.03361673 |
| Rasl11b | 5.83327 | 10.1908 |
| Safb | 21.2071 | 37.0487 |
| Obsl1 | 37.68809 | 65.83697 |
| Vash1 | 4.60255 | 8.03833 |
| Swap70 | 6.363291368 | 11.11275456 |
| Baiap3 | 1.40172 | 2.44779 |
| Snai1 | 11.1171 | 19.4126 |
| Cchcr1 | 3.87036033 | 6.7575803 |
| Zmynd8 | 8.37722511 | 14.6229922 |
| Map7d1 | 19.74934367 | 34.47018 |
| Foxo3 | 8.079868 | 14.101337 |
| Dctn1 | 20.15609223 | 35.17411346 |
| Tsr1 | 6.07722 | 10.6019 |
| Mamld1 | 1.343912 | 2.34438 |
| Zfp628 | 3.57924 | 6.24158 |
| Ikbkb | 5.108433 | 8.90726 |
| Adgrl3 | 3.621184062 | 6.3135166 |
| Slitrk5 | 0.753950988 | 1.314212242 |
| Thsd7b | 1.43105 | 2.49432 |
| Isoc2b | 1.562584 | 2.722533724 |
| Sh3pxd2b | 6.26844 | 10.9216 |

| | | |
|---|---|---|
| Trim71 | 2.247338 | 3.915361 |
| Mab21l3 | 0.557882 | 0.971774 |
| Gcn1l1 | 14.17450771 | 24.6890162 |
| Znf41-ps | 1.97571 | 3.44093 |
| Wdfy3 | 3.4111378 | 5.93977917 |
| Kcnd3 | 1.704641826 | 2.968212491 |
| Msx1 | 3.63423 | 6.3279 |
| Ak4 | 2.7583168 | 4.802389788 |
| Irf2bpl | 13.0468 | 22.7132 |
| Heg1 | 4.274636 | 7.43946 |
| Xpr1 | 2.68791 | 4.677434 |
| Smpd4 | 8.59232145 | 14.94889 |
| Rtf1 | 7.93489 | 13.80412662 |
| Bhlhe40 | 2.32854 | 4.04882 |
| Hnrnpul1 | 31.482908 | 54.740632 |
| Erap1 | 1.10288739 | 1.9165365 |
| Tmem181k | 4.141068 | 7.19577 |
| Snx30 | 1.62853 | 2.82954 |
| Plag1 | 2.113926305 | 3.672764 |
| Adat1 | 1.89687 | 3.29541 |
| Tmem201 | 12.47516439 | 21.667184 |
| Ablim1 | 3.596888331 | 6.246294 |
| Mtcl1 | 3.097255048 | 5.376306486 |
| Hnrnpul2 | 19.6858 | 34.171 |
| Epg5 | 1.03676 | 1.79938 |
| Aoc2 | 0.940763 | 1.63223 |
| Cdc42bpa | 2.843631799 | 4.92949442 |
| Plcb2 | 0.626223549 | 1.085227417 |
| Slc9a8 | 3.488576829 | 6.04296389 |
| Setbp1 | 0.821366 | 1.42273 |
| Ntf3 | 1.999029968 | 3.462416964 |
| Prkab2 | 1.6364 | 2.83402 |
| Exoc3l | 3.11829 | 5.3995 |
| Stk36 | 2.025946 | 3.507008753 |
| Net1 | 12.64305691 | 21.885415 |
| Csf1 | 1.685159 | 2.916751 |
| Foxred2 | 1.297031963 | 2.244875 |
| Hpca | 1.633348169 | 2.826309858 |
| Trim6 | 2.871208703 | 4.967597 |
| Drosha | 15.11957162 | 26.154037 |
| Tfcp2 | 4.82198201 | 8.340633 |
| Herc2 | 2.53143 | 4.37853 |
| Atp13a2 | 14.74144 | 25.49299 |
| Notch1 | 2.21477 | 3.83002 |
| Pcsk5 | 4.3115149 | 7.4549726 |
| Nsl1 | 3.32567 | 5.74944 |
| Ehbp1l1 | 2.13398924 | 3.688912338 |

| | | |
|---|---|---|
| Ercc6 | 1.899101 | 3.2827673 |
| Syt7 | 1.50760173 | 2.605852905 |
| Glis2 | 20.7677 | 35.89 |
| Pidd1 | 2.589658 | 4.47406 |
| Per1 | 3.2263785 | 5.571876678 |
| Setd5 | 14.6808248 | 25.3338536 |
| C2cd2l | 1.301989099 | 2.245241 |
| Zbtb21 | 1.8758434 | 3.234772172 |
| Rrp1b | 5.487685764 | 9.4628553 |
| Pnisr | 9.97011 | 17.1911613 |
| Cdh2 | 8.4147 | 14.5092 |
| Samd14 | 11.293 | 19.4632 |
| Slc12a5 | 0.676088211 | 1.164936 |
| Ccdc88c | 1.41481 | 2.43706 |
| Cd52 | 6.45765 | 11.1231 |
| Nfatc1 | 0.8146985 | 1.4031193 |
| Pisd-ps2 | 1.858900345 | 3.201008284 |
| Spry4 | 2.0172 | 3.47348 |
| Ly6e | 34.267842 | 58.99846205 |
| Myo7a | 1.83718715 | 3.16242176 |
| Sbspon | 2.852099 | 4.90939 |
| Pear1 | 4.8890894 | 8.414270224 |
| Syde1 | 10.57071 | 18.19184 |
| Scaf4 | 5.39533801 | 9.2851854 |
| E2f3 | 4.71167281 | 8.10787 |
| Glis3 | 0.963377 | 1.657662 |
| Vstm2b | 2.657869 | 4.57086 |
| Brd3 | 14.95339681 | 25.705262 |
| Agtr2 | 28.8914 | 49.6581 |
| Col5a1 | 22.13567 | 38.0375 |
| Lrfn2 | 1.24419 | 2.13758 |
| Kif3c | 2.05279 | 3.52565671 |
| Phactr2 | 2.831698 | 4.8609924 |
| Accs | 1.69573755 | 2.910956146 |
| Hoxa5 | 4.15186 | 7.12515 |
| Slit2 | 4.6066297 | 7.900580633 |
| D5Ertd579 | 3.31002 | 5.67168 |
| Sgsm2 | 4.003859 | 6.85627 |
| Gramd1b | 0.8897743 | 1.5235632 |
| Ptprj | 1.6598443 | 2.841915687 |
| Hnrnpa2b1 | 134.16778 | 229.627732 |
| Trrap | 20.1864493 | 34.542211 |
| Egfl6 | 2.511655752 | 4.29684 |
| Plekho2 | 2.83895 | 4.85591 |
| Zfp382 | 0.571705575 | 0.97785321 |
| Xylt2 | 7.06611 | 12.08437 |
| Gm17296 | 1.49403 | 2.555 |

| | | |
|---|---|---|
| Gm9833 | 0.70758 | 1.20938 |
| Pik3ap1 | 0.695576 | 1.18864 |
| Tmem131 | 6.553444 | 11.19889 |
| Plxna2 | 1.97827 | 3.37941 |
| Supt7l | 4.87132 | 8.31637 |
| Arhgap17 | 6.3430948 | 10.828493 |
| Lrrk1 | 1.864192 | 3.18103 |
| Frmd4a | 3.746996124 | 6.392767412 |
| Rab11b | 17.1150845 | 29.19453942 |
| Rttn | 0.9822607 | 1.675458 |
| Abcg5 | 0.558779 | 0.953033 |
| Fam160a2 | 4.484014514 | 7.644245986 |
| Dusp7 | 12.5213 | 21.3413 |
| Alx4 | 0.875006 | 1.49076 |
| Ascc3 | 2.306573 | 3.92848 |
| Chrd | 1.01448167 | 1.727778 |
| Ccdc9 | 11.778094 | 20.05782 |
| Tecpr1 | 4.4036 | 7.49687 |
| Ampd3 | 0.963818922 | 1.639400265 |
| Numbl | 8.59469 | 14.61749 |
| Pdcd11 | 5.80414 | 9.86024 |
| Bmper | 4.38491 | 7.44868 |
| Nf2 | 11.58272133 | 19.66945278 |
| Cand2 | 9.33236 | 15.846 |
| Stx16 | 8.598426882 | 14.59186 |
| Hoxb4 | 1.42533 | 2.41867 |
| Snhg20 | 25.0179 | 42.4511 |
| Loxl3 | 2.82969 | 4.80013 |
| Ubald1 | 17.77362 | 30.14862 |
| Eng | 13.120497 | 22.255088 |
| Cabin1 | 13.0266759 | 22.091078 |
| Herc3 | 1.76031079 | 2.98517456 |
| Hmgxb3 | 5.66348 | 9.60359 |
| Socs7 | 3.502113 | 5.93715 |
| Rarg | 5.920681 | 10.03717 |
| Tmppe | 1.36807 | 2.31802 |
| Clstn1 | 15.07800446 | 25.54519 |
| Tnk2 | 15.065126 | 25.51943971 |
| Heatr5b | 1.822456 | 3.086168 |
| Lyst | 0.623079 | 1.05485 |
| St8sia1 | 2.280454222 | 3.860143415 |
| Lamb1 | 16.66617 | 28.2042 |
| Armcx4 | 2.8557083 | 4.83258753 |
| Inpp4a | 2.1340979 | 3.61106024 |
| Zfp651 | 4.30107 | 7.27654 |
| Eefsec | 8.58236 | 14.5196 |
| BC030500 | 0.584956 | 0.989439 |

| | | |
|---|---|---|
| Gli3 | 3.65674 | 6.18468 |
| Inpp5d | 1.251629915 | 2.11687378 |
| Gsg1 | 4.614124033 | 7.801827013 |
| Kif13b | 2.4524541 | 4.145805569 |
| Wnt7b | 0.686694 | 1.1606862 |
| Tesk1 | 11.11985 | 18.7945 |
| Plxna4 | 2.39963 | 4.05444 |
| Tom1l2 | 5.2458604 | 8.862426 |
| Nvl | 7.26222 | 12.2670472 |
| St3gal1 | 6.38964 | 10.79296 |
| Rdh10 | 8.822028 | 14.901587 |
| Fubp3 | 9.17708 | 15.50064 |
| Faim2 | 0.706128 | 1.192661156 |
| Cpne5 | 3.2642 | 5.51322 |
| Cblb | 3.27658404 | 5.533785 |
| Usp48 | 11.25646808 | 19.00911 |
| Pan3 | 7.888428 | 13.320576 |
| Arnt2 | 5.12076723 | 8.6454006 |
| Stac2 | 2.16096 | 3.64794 |
| Paxip1 | 3.9264325 | 6.625357006 |
| Frmd4b | 10.2994389 | 17.3772 |
| Cipc | 4.68831 | 7.909931873 |
| Bcl2l2 | 4.72615 | 7.973665 |
| Acvr2b | 4.10481132 | 6.925278 |
| Plxnb2 | 28.90812 | 48.76336194 |
| Rftn1 | 3.1545 | 5.3194 |
| Nbea | 4.321095495 | 7.283389 |
| Src | 15.19585158 | 25.59706 |
| Pom121 | 13.1602 | 22.1678 |
| Nrk | 4.142825 | 6.97738 |
| Dtx1 | 2.236163 | 3.76608 |
| Prdm10 | 1.13436026 | 1.910374994 |
| Pdcd4 | 18.94499 | 31.89523 |
| Epn1 | 29.5295956 | 49.7105633 |
| Gata2 | 28.02759 | 47.16428 |
| Hnf1b | 1.79409446 | 3.019012631 |
| Atxn7 | 0.961046 | 1.616763403 |
| Eya3 | 7.63149 | 12.83724 |
| Ppp1r21 | 5.56983 | 9.36618 |
| Rapgef1 | 10.57546 | 17.782452 |
| Sfi1 | 4.12437623 | 6.93306 |
| Safb2 | 19.16627 | 32.213 |
| Timp2 | 24.646 | 41.4175 |
| Ift122 | 4.19038 | 7.041302986 |
| Rictor | 3.23447 | 5.43474 |
| Fras1 | 2.5076 | 4.21337 |
| St14 | 1.9969002 | 3.35507987 |

| | | |
|---|---|---|
| Ctdsp2 | 70.9699 | 119.2279 |
| Ntm | 1.4721632 | 2.472421 |
| Nynrin | 21.79812 | 36.60485 |
| Crebzf | 10.21628574 | 17.15579927 |
| Mbd5 | 1.493594406 | 2.507462473 |
| Ptpn13 | 7.372907 | 12.37513583 |
| Igdcc3 | 7.16486028 | 12.02505 |
| Fam120c | 1.12009 | 1.87966 |
| Crim1 | 1.89645 | 3.18221 |
| Strn4 | 19.99804 | 33.5561 |
| Dyrk1b | 9.83581888 | 16.49737054 |
| Slc4a4 | 0.97301932 | 1.631884979 |
| Tmem30b | 0.571762 | 0.958916 |
| Ddr2 | 2.896006 | 4.85427 |
| Myo1d | 2.94787 | 4.93714 |
| Prdm15 | 2.066584981 | 3.460266516 |
| Cercam | 8.32289 | 13.93541 |
| Dnajc13 | 5.63104 | 9.42609 |
| Palm3 | 3.167194 | 5.300919085 |
| Ttc3 | 18.281354 | 30.592713 |
| Ccdc166 | 2.102247476 | 3.516645272 |
| Supt5 | 28.4331 | 47.5568 |
| Btbd17 | 0.489254427 | 0.818256 |
| Prpf38b | 9.741021154 | 16.289821 |
| Vps13b | 1.590122 | 2.658484 |
| Csnk1g1 | 2.642012402 | 4.4168072 |
| Aldh1a2 | 51.8056 | 86.5193 |
| Pik3r4 | 4.77103507 | 7.966795 |
| Usp35 | 1.753776 | 2.9279132 |
| Wipf3 | 1.094120136 | 1.826264001 |
| Slc8a1 | 0.863884909 | 1.441717932 |
| Hmha1 | 5.083771 | 8.483506 |
| Atp2a2 | 19.914326 | 33.228706 |
| Peli2 | 2.10493 | 3.51111 |
| Ranbp10 | 5.53268 | 9.22815 |
| Atrnl1 | 17.37246 | 28.97442 |
| Ankrd44 | 1.40382 | 2.3408 |
| Bach2 | 1.403922 | 2.34089608 |
| Mapk8ip2 | 1.314774 | 2.192212 |
| Atrip | 4.021984755 | 6.70603 |
| Rnf31 | 7.683773 | 12.80669 |
| Prr5l | 2.790200168 | 4.64991275 |
| Plekhh3 | 5.615818183 | 9.353850269 |
| Rab11fip4 | 3.6633225 | 6.101105 |
| Tie1 | 6.9344 | 11.5482 |
| Copa | 33.256 | 55.3821 |
| Exoc4 | 6.67319 | 11.1102 |

| | | |
|---|---|---|
| Vprbp | 3.548298 | 5.90634 |
| Mylk | 5.8605281 | 9.748423 |
| Stk26 | 6.39931 | 10.64431 |
| Tor3a | 2.35695 | 3.92025 |
| S100pbp | 14.35679646 | 23.87589015 |
| Fam120a | 19.1251 | 31.7833 |
| Wdr35 | 2.927512963 | 4.864519115 |
| Ston2 | 0.858785988 | 1.426955624 |
| Plcg1 | 17.2205991 | 28.611494 |
| Plk1 | 20.0625 | 33.3332 |
| Nckipsd | 2.74038 | 4.55188 |
| Naaa | 12.47368 | 20.717321 |
| Hunk | 9.00462 | 14.95472 |
| Triml2 | 1.59864 | 2.654615 |
| Myo9b | 5.718014229 | 9.494916 |
| D330050I1 | 1.80332 | 2.99444 |
| Plce1 | 0.880704206 | 1.461735407 |
| Ap4e1 | 2.175944 | 3.6113562 |
| Slc38a6 | 1.510883884 | 2.50712171 |
| Zfp326 | 6.4891309 | 10.76722 |
| Gm20735 | 0.715674 | 1.18746 |
| Prox2 | 0.867535551 | 1.43901 |
| Uhrf1bp1 | 5.51138 | 9.14165 |
| Foxj3 | 5.52182 | 9.15445 |
| Zfp553 | 10.21864 | 16.93869 |
| Sltm | 10.0891052 | 16.722793 |
| Pik3r1 | 10.289597 | 17.05188 |
| Ocrl | 3.9675947 | 6.57340894 |
| Herc1 | 5.76131 | 9.543825 |
| Tub | 0.614506 | 1.01767 |
| Akap9 | 2.20462595 | 3.65072889 |
| Hivep2 | 0.491644829 | 0.814061618 |
| Nr6a1 | 3.8095 | 6.306397 |
| Ppp4r1l-ps | 3.147191562 | 5.2076069 |
| Wipf2 | 2.42873 | 4.01854 |
| Jak3 | 4.29565222 | 7.106807 |
| Rnf43 | 2.240443 | 3.706476 |
| Slc12a9 | 3.79007 | 6.26843 |
| Mtmr12 | 3.32009374 | 5.48827052 |
| Cdk2 | 16.10348 | 26.61937794 |
| Igsf3 | 3.77683 | 6.24314 |
| Arf3 | 29.91413 | 49.43072 |
| Bbs4 | 2.31174 | 3.8198 |
| Col15a1 | 6.594112 | 10.89471 |
| Stard13 | 0.75924038 | 1.254276558 |
| Frs2 | 4.46503 | 7.3747434 |
| Epha2 | 2.33737 | 3.85979 |

| | | |
|---|---|---|
| Meg3 | 93.6038345 | 154.562965 |
| Mvb12b | 7.966 | 13.15004 |
| Sec16a | 6.61591122 | 10.92010291 |
| Gpatch2l | 3.784323 | 6.24600916 |
| Nfx1 | 10.748856 | 17.73976 |
| Igf2bp2 | 22.41141034 | 36.98584675 |
| Kif13a | 2.89312264 | 4.77374 |
| Tmed8 | 1.5416 | 2.54349 |
| Hs3st3b1 | 4.76898 | 7.86815 |
| Mthfr | 3.983511277 | 6.572198 |
| Mecp2 | 2.355324297 | 3.8836 |
| Egr1 | 2.15324 | 3.54978 |
| Jun | 17.0063 | 28.0186 |
| Epn3 | 3.63063787 | 5.980653 |
| Gpld1 | 1.79629 | 2.95867 |
| Utp20 | 3.20912 | 5.28547 |
| Prpf39 | 4.08641 | 6.7298 |
| Map3k11 | 9.09852 | 14.980287 |
| Synj2 | 1.023141 | 1.68444455 |
| BC024978 | 1.91645377 | 3.15492182 |
| Agap3 | 19.68466 | 32.40317 |
| C230035I1( | 3.741872 | 6.158834814 |
| Agl | 1.68764959 | 2.777552017 |
| Oxsm | 0.825502 | 1.35862 |
| Jup | 9.8385 | 16.1872287 |
| Afap1l2 | 2.688491106 | 4.422574693 |
| Dido1 | 5.2576451 | 8.6474389 |
| B2m | 119.3 | 196.137 |
| Zcchc24 | 14.13385 | 23.22891 |
| Ttll1 | 2.90372 | 4.77113 |
| Dock4 | 3.4088 | 5.60032 |
| Tab1 | 7.72106 | 12.6776 |
| Atf7ip | 9.39228 | 15.41917 |
| Gxylt1 | 5.79950207 | 9.5206851 |
| Cnnm3 | 5.41451 | 8.887772 |
| Dpysl4 | 3.0701 | 5.0387 |
| C2 | 1.83082 | 3.00427 |
| Ubr1 | 2.78471834 | 4.568113 |
| Fbxl20 | 4.077142992 | 6.687447 |
| Lonrf2 | 1.30104 | 2.13392 |
| D930048N1 | 1.09178 | 1.79038 |
| D930016D( | 1.99214695 | 3.26657482 |
| Dpy19l3 | 2.535355 | 4.15712 |
| Arhgap21 | 4.32437745 | 7.09042039 |
| Tro | 3.20451548 | 5.25417648 |
| Zfp398 | 1.0960305 | 1.7970159 |
| Pogz | 7.623317 | 12.49711 |

| | | |
|---|---|---|
| Telo2 | 5.57589 | 9.14003572 |
| Tiam1 | 3.439293844 | 5.63590895 |
| Marf1 | 10.45366494 | 17.124322 |
| Ucp2 | 69.1288 | 113.198 |
| Zbed3 | 19.38816 | 31.7465 |
| Foxj2 | 3.0840866 | 5.0481045 |
| Slc12a6 | 3.8027847 | 6.22274 |
| Agpat1 | 29.64838742 | 48.5102546 |
| Tti1 | 5.79367602 | 9.47839 |
| Trp53i11 | 8.9341219 | 14.6159 |
| Zcchc3 | 31.5069 | 51.5416 |
| Gm13251 | 2.49308 | 4.07801 |
| Trappc11 | 6.84087 | 11.1892 |
| Zbtb39 | 3.137606 | 5.13182 |
| Fam13b | 5.55631 | 9.08771 |
| Wdfy2 | 3.21794 | 5.26294205 |
| Abcc10 | 2.00305 | 3.2749499 |
| Ism1 | 0.696146 | 1.13753 |
| Kat2b | 1.549357644 | 2.531506945 |
| Fam135a | 3.900250458 | 6.37167721 |
| Brd4 | 13.3400115 | 21.792479 |
| Adcyap1r1 | 2.487955459 | 4.064356 |
| Zim1 | 9.572493601 | 15.63704315 |
| Nos3 | 1.51695 | 2.47781 |
| Mir1932 | 0.982917 | 1.60508 |
| Sorbs3 | 24.24797433 | 39.5802945 |
| Cdk5r1 | 0.840413904 | 1.371705 |
| Tbc1d31 | 2.351729663 | 3.837099997 |
| Tpp1 | 9.12507 | 14.8878 |
| Scaf11 | 8.029845979 | 13.1003 |
| Map3k2 | 0.917351 | 1.49633 |
| Kidins220 | 5.698317 | 9.29419626 |
| Fcgr1 | 1.51398 | 2.46886 |
| Il18bp | 0.993341 | 1.619847 |
| Lyz2 | 10.7822 | 17.5801 |
| Mterf1b | 0.814844 | 1.32786 |
| Gm16386 | 0.744775059 | 1.213477614 |
| Kdm4b | 8.31985 | 13.55194 |
| Spi1 | 2.97154 | 4.83994 |
| Eef2k | 2.810158515 | 4.576736257 |
| Penk | 3.953021 | 6.43623 |
| Hyal3 | 5.536229 | 9.010747 |
| Nmi | 3.193049622 | 5.195546 |
| Evc | 4.3202 | 7.02911 |
| Nsd1 | 7.15163952 | 11.6355633 |
| Psors1c2 | 1.179884629 | 1.919481 |
| Gm11747 | 1.59427 | 2.59224 |

| | | |
|---|---|---|
| Srebf1 | 8.71206 | 14.161748 |
| Zswim4 | 7.89631 | 12.83466499 |
| Smarcd1 | 19.4816 | 31.6543 |
| Hmga2 | 99.01485037 | 160.862882 |
| Uba1 | 90.8729 | 147.593809 |
| Amer2 | 0.616829976 | 1.001508 |
| Tmem245 | 1.409661139 | 2.287788125 |
| Stk11ip | 9.05863 | 14.6973 |
| Slc38a7 | 5.09497435 | 8.26345 |
| Scrt2 | 3.32803 | 5.39528 |
| Frmd6 | 9.78713879 | 15.86362 |
| Ccdc57 | 1.79267216 | 2.904636 |
| Mapkapk3 | 1.45968865 | 2.365014806 |
| Zfp444 | 6.23836476 | 10.1019 |
| Pyroxd1 | 1.29261 | 2.09311 |
| Ephb3 | 4.36657 | 7.07036 |
| Dlgap5 | 4.20392 | 6.80346 |
| Mlxip | 5.29634 | 8.5695 |
| Slc27a1 | 9.80892 | 15.8703 |
| Msh5 | 0.7473666 | 1.20855405 |
| Tmtc3 | 2.057373737 | 3.32645325 |
| Myo10 | 5.413646 | 8.75248 |
| Micall2 | 1.854785 | 2.99849757 |
| Wdr90 | 5.05281544 | 8.16787 |
| Hip1 | 5.9332996 | 9.59041 |
| Sptan1 | 16.82671088 | 27.19670112 |
| Arhgap10 | 6.8431 | 11.05825 |
| Gpc2 | 10.4561 | 16.88765 |
| Kcnma1 | 0.7852405 | 1.267971847 |
| Necab2 | 3.700623291 | 5.974746949 |
| Rsrp1 | 70.9413 | 114.5355 |
| Plekhm2 | 5.92705436 | 9.56888 |
| F11r | 20.6154 | 33.279949 |
| Usp34 | 3.455011975 | 5.576007429 |
| Cwc22 | 4.920346585 | 7.939929898 |
| Maml1 | 6.61105 | 10.6668 |
| Cd300a | 0.80113013 | 1.29255 |
| Unkl | 4.1233578 | 6.65246136 |
| Grb10 | 60.48577897 | 97.57983821 |
| Ccnd2 | 25.299076 | 40.80754 |
| Ticrr | 2.62678 | 4.23651 |
| Gtf3c2 | 15.462213 | 24.93537 |
| Irf2bp2 | 14.557042 | 23.46904 |
| Tef | 7.668577392 | 12.36302 |
| Zmynd11 | 11.42631537 | 18.41806937 |
| Ino80 | 3.82194 | 6.159901947 |
| U2surp | 5.254403 | 8.4683798 |

| | | |
|---|---|---|
| Tenm4 | 8.87080691 | 14.29392803 |
| Prkd2 | 4.237672 | 6.82826 |
| Aldh2 | 63.9941 | 103.106535 |
| Irf2 | 1.750941576 | 2.819994 |
| Odf2 | 11.56999898 | 18.63327699 |
| Eftud1 | 2.37580218 | 3.82527 |
| Vat1l | 1.56747 | 2.52347 |
| Pkp4 | 5.598727131 | 9.009599003 |
| Slit3 | 2.39476 | 3.85239 |
| Ptpn11 | 14.74766676 | 23.72055 |
| Parp6 | 4.28584284 | 6.88964948 |
| Gpsm1 | 10.89541 | 17.51466 |
| Creb3l2 | 9.47559 | 15.2321 |
| Cul7 | 20.7694 | 33.3787 |
| Clk1 | 26.09569 | 41.93736 |
| Meis1 | 24.57907923 | 39.49809 |
| Mafb | 3.46697 | 5.57032 |
| Usp21 | 26.2961 | 42.249 |
| Ddx23 | 17.35822 | 27.88777396 |
| Tle2 | 10.704629 | 17.1892 |
| Abcc1 | 3.29281548 | 5.285779 |
| Tubgcp4 | 3.23842 | 5.19822 |
| Bora | 5.24741699 | 8.4224893 |
| Apobec3 | 4.83468382 | 7.759522 |
| Pdk2 | 11.22082 | 18.00766 |
| D630003M | 1.75952 | 2.82336 |
| Pcyt2 | 12.13492 | 19.47054 |
| Pogk | 4.63496 | 7.43667 |
| Slc35e2 | 2.70204 | 4.335314468 |
| Efs | 7.99019 | 12.8171 |
| Srsf5 | 68.2921 | 109.545 |
| Rcor1 | 7.62577 | 12.2319 |
| Slc26a11 | 3.354125 | 5.379622838 |
| Mllt10 | 5.123483005 | 8.213289858 |
| Cbx4 | 2.71572 | 4.35324 |
| Plscr4 | 1.19084 | 1.90859 |
| Usp25 | 2.99677 | 4.80165 |
| Atrn | 2.612117 | 4.1846 |
| Nckap1 | 17.24066967 | 27.61580147 |
| Fam193a | 8.435781 | 13.505927 |
| Mob3b | 0.823699 | 1.31868 |
| Gpc6 | 8.835976885 | 14.14403155 |
| Ids | 2.66498 | 4.26463 |
| Ust | 1.7691 | 2.83088 |
| Ralgds | 12.33718952 | 19.740395 |
| Pisd-ps1 | 11.7177 | 18.7455 |
| Col4a1 | 21.95445 | 35.09777818 |

| | | |
|---|---|---|
| Pan2 | 8.190717 | 13.091239 |
| Gtpbp2 | 12.0814 | 19.30343547 |
| Lsm14b | 17.998553 | 28.74517 |
| Zfp689 | 2.922667 | 4.665520871 |
| Sipa1l2 | 5.12798 | 8.181399 |
| Pltp | 12.1778 | 19.4196 |
| Uvssa | 0.81202984 | 1.294869742 |
| Rian | 43.8216 | 69.8516 |
| Dennd6b | 1.739853079 | 2.773031062 |
| Nbr1 | 12.12798 | 19.32472525 |
| Dhx38 | 9.457885665 | 15.06911118 |
| Mettl3 | 8.27998 | 13.1903 |
| Phf3 | 5.424630296 | 8.639731598 |
| Sh3gl2 | 1.69888 | 2.70568 |
| Cachd1 | 5.2752 | 8.401290599 |
| Itprip | 3.92537 | 6.25143 |
| Tspyl2 | 5.866554274 | 9.34172 |
| Pigo | 5.847519 | 9.311130799 |
| Dock7 | 7.938331741 | 12.6369848 |
| Alkbh1 | 6.28481 | 10.0044 |
| Cyp26b1 | 2.897384567 | 4.6120831 |
| Ly96 | 1.849235432 | 2.94336 |
| Diexf | 3.36774 | 5.35958 |
| Med16 | 16.0965 | 25.6165 |
| Aars2 | 5.38767 | 8.57328 |
| Smg5 | 11.0944749 | 17.6479133 |
| Myh11 | 1.04626 | 1.66426 |
| Otud7b | 4.395253496 | 6.989571 |
| Ehmt1 | 10.767473 | 17.11195482 |
| Mapk7 | 14.82742496 | 23.562989 |
| Flii | 11.08483 | 17.61418 |
| Cdan1 | 3.3443 | 5.3135676 |
| Dsp | 2.6557901 | 4.21919 |
| Flnc | 27.42731 | 43.569 |
| Greb1 | 2.469619286 | 3.921841349 |
| Ube4a | 6.078092416 | 9.64823716 |
| Ttc14 | 2.132603368 | 3.384781702 |
| Cpox | 4.21213 | 6.68447 |
| Arhgef28 | 2.10025782 | 3.332865 |
| Slc4a7 | 1.87719571 | 2.978235408 |
| Mamdc4 | 0.94398266 | 1.497391 |
| Fbxl18 | 3.418046 | 5.420609 |
| Pag1 | 1.067137554 | 1.691899 |
| Ttll4 | 5.242012428 | 8.31082734 |
| Slc16a3 | 17.68544911 | 28.037112 |
| Zdhhc17 | 3.672871375 | 5.822374386 |
| Sec22c | 2.22167113 | 3.521762678 |

| | | |
|---|---|---|
| Kpna6 | 7.96505 | 12.6219795 |
| Llgl2 | 4.639623 | 7.351888 |
| Ints1 | 9.661809 | 15.30971 |
| Hcfc1 | 13.93976557 | 22.08334991 |
| Zscan20 | 1.36197 | 2.15757445 |
| Zhx3 | 4.228231 | 6.697553993 |
| Abhd2 | 3.643801 | 5.77094 |
| Prr5 | 14.51765 | 22.98748 |
| Slc11a2 | 9.82082 | 15.54789 |
| Igdcc4 | 14.0451163 | 22.23019866 |
| Slc41a1 | 3.5619071 | 5.637614 |
| Alpl | 9.84714685 | 15.58185438 |
| Ezh1 | 4.72666368 | 7.476563453 |
| Abca4 | 0.691635474 | 1.09397605 |
| Def6 | 1.3087 | 2.06995 |
| Syk | 3.478132544 | 5.500896 |
| Dact3 | 12.7193 | 20.1149 |
| Larp1 | 15.23205 | 24.08737 |
| Zfp94 | 1.43980967 | 2.276858 |
| Wdr91 | 5.3264013 | 8.422913 |
| Atg9a | 6.84416575 | 10.82223282 |
| Nucb1 | 30.637977 | 48.42823 |
| Mboat1 | 1.54157 | 2.43636 |
| Cldn4 | 8.38988 | 13.2573 |
| Xkr5 | 0.830688688 | 1.3126146 |
| Recql4 | 2.43849 | 3.8529 |
| N4bp1 | 6.66544 | 10.5306 |
| Itpr1 | 5.8341754 | 9.216161 |
| Helq | 1.91217 | 3.02049 |
| Zfp598 | 14.40923832 | 22.75569454 |
| Sh3tc1 | 1.453398 | 2.29487461 |
| Cyp27a1 | 1.33416 | 2.10648 |
| Zfp768 | 11.9485 | 18.8611 |
| Dmwd | 26.59932 | 41.9833 |
| Slc4a2 | 11.11597 | 17.54115 |
| Sesn2 | 4.73653 | 7.47305 |
| Sbno1 | 5.0326507 | 7.93699711 |
| Mpdz | 5.545533372 | 8.74508714 |
| Sema6d | 5.5619685 | 8.76892203 |
| Slx4 | 4.169825181 | 6.569375167 |
| Pld2 | 2.3353082 | 3.679018241 |
| Agap1 | 4.062335 | 6.398136 |
| Met | 1.39017 | 2.18934 |
| Znrf1 | 8.980691 | 14.137785 |
| Gpr161 | 2.171362009 | 3.418109 |
| Crb2 | 4.6245199 | 7.2780037 |
| Pvrl2 | 39.614 | 62.333 |

| | | |
|---|---|---|
| Cys1 | 1.624189541 | 2.554887 |
| Cep128 | 1.035931653 | 1.629535 |
| Asap1 | 6.3559586 | 9.9925411 |
| Etnk1 | 10.6625531 | 16.76087 |
| Eln | 5.24007679 | 8.236493867 |
| Abcd1 | 5.936884069 | 9.33079 |
| Etv5 | 12.9216 | 20.3027 |
| Tmem151b | 2.49563 | 3.92079 |
| Zc3h7a | 4.0418 | 6.34806 |
| Mreg | 2.63557 | 4.13941 |
| Dst | 4.571866256 | 7.17665871 |
| Tsku | 4.91716055 | 7.71732617 |
| Cdc14a | 1.592930802 | 2.498836456 |
| Cdc14b | 3.514853 | 5.5137568 |
| Cog3 | 4.95204 | 7.767 |
| Kdm5b | 16.776307 | 26.30984 |
| C2cd3 | 7.540647 | 11.82556 |
| Evi5l | 5.376407 | 8.425669 |
| Baz1a | 1.21655 | 1.906338109 |
| Foxl2os | 1.139719 | 1.78592 |
| D2hgdh | 4.667788174 | 7.313425 |
| Rmi2 | 1.57444 | 2.46629 |
| Emc1 | 4.6114472 | 7.222876092 |
| Zfp445 | 8.266477 | 12.9444225 |
| Pycr1 | 5.981533 | 9.364688 |
| Elp2 | 32.5374 | 50.9322 |
| Chrnb1 | 1.53833 | 2.40559 |
| Klhl42 | 1.82489 | 2.85355 |
| Nedd4 | 144.325 | 225.674 |
| Kcnab2 | 0.9708742 | 1.518074757 |
| Rabggta | 7.052237 | 11.02696854 |
| Zcchc14 | 9.07621 | 14.19059 |
| Ndnf | 1.419214724 | 2.218760203 |
| Ncoa1 | 2.028510635 | 3.170694 |
| Sh3rf1 | 3.11169 | 4.86314 |
| Ppp1r37 | 14.7865 | 23.10879 |
| Cntrob | 8.191467008 | 12.800783 |
| Lrch1 | 1.6102528 | 2.51578 |
| Zfp866 | 2.97279 | 4.64388 |
| Usp24 | 4.31599126 | 6.7418278 |
| Adgrl1 | 18.702952 | 29.20531 |
| Rragd | 1.3090757 | 2.0437111 |
| Mmp15 | 16.76768 | 26.1654 |
| Robo1 | 2.4446898 | 3.8147593 |
| Gnaq | 6.944020906 | 10.83562 |
| Pik3c3 | 6.25324 | 9.75709 |
| Mylip | 5.28792 | 8.24577 |

| | | |
|---|---|---|
| Glcci1 | 1.7867388 | 2.784695253 |
| Ss18l1 | 1.57061 | 2.44741 |
| Pkd2 | 9.94159 | 15.4858 |
| Rnf144b | 1.45499328 | 2.266212251 |
| Sorcs2 | 3.005538525 | 4.68031492 |
| Smad9 | 0.90414112 | 1.407361071 |
| Wbp2 | 22.0476 | 34.3118 |
| Diap1 | 6.3719194 | 9.91382 |
| H2-T22 | 5.50275 | 8.56073 |
| Ulk4 | 1.249787485 | 1.944297655 |
| Fgd6 | 1.19883 | 1.86452 |
| Slc36a1 | 2.670775 | 4.15339 |
| Phf1 | 7.323396 | 11.38785 |
| Hs3st3a1 | 4.14984 | 6.45151 |
| Zzef1 | 2.759534581 | 4.2894969 |
| Dach1 | 3.004263836 | 4.6673515 |
| Spats2l | 2.877427112 | 4.46997575 |
| Slc26a6 | 1.405841206 | 2.183539 |
| Srebf2 | 25.5895 | 39.7378 |
| B4galt3 | 14.78232 | 22.9466 |
| Faxc | 1.4126593 | 2.192859 |
| Nxt2 | 2.30662014 | 3.580389 |
| Zfp575 | 1.0217 | 1.58585 |
| Tnks | 5.740048 | 8.90869 |
| Hspa1l | 0.713525 | 1.10726 |
| Fkbp15 | 3.30083 | 5.12106 |
| Pigb | 1.66655 | 2.58539 |
| Tut1 | 7.97499 | 12.37112 |
| Zscan2 | 7.837891 | 12.1575907 |
| Rab6b | 2.43604 | 3.77667 |
| Nptxr | 1.36251 | 2.1121 |
| Bcl2l12 | 17.315933 | 26.8386 |
| Eda2r | 0.973627545 | 1.508634519 |
| Lppr3 | 21.706695 | 33.62603 |
| Nes | 6.31152 | 9.77547 |
| Irs1 | 4.83421 | 7.48463 |
| Gm15612 | 0.878895 | 1.36038 |
| Mapk1ip1l | 23.6749 | 36.6438 |
| Chst11 | 3.6827454 | 5.7001 |
| Wsb1 | 23.635873 | 36.58090514 |
| Ganc | 0.928003 | 1.43618 |
| Apbb3 | 4.81153 | 7.44551 |
| Zc3hav1l | 8.8322 | 13.66286 |
| Nol6 | 14.7256 | 22.7786 |
| Arhgap9 | 1.42795 | 2.20846 |
| Mdm2 | 5.498009 | 8.50176 |
| Ltbp1 | 12.46163022 | 19.259903 |

| | | |
|---|---|---|
| Cpz | 4.623200046 | 7.144827385 |
| Kctd18 | 3.87121995 | 5.98212101 |
| Ttc17 | 2.54107 | 3.92662 |
| Glg1 | 13.7224 | 21.2017 |
| Ccnl2 | 32.0945 | 49.5858 |
| Dusp4 | 5.63933 | 8.71183 |
| Pcyt1b | 0.81828503 | 1.26396673 |
| Senp1 | 4.51778 | 6.97632 |
| Pou2f1 | 2.006054003 | 3.096511512 |
| Ccar2 | 17.35443 | 26.7805 |
| Serac1 | 1.209397 | 1.866177 |
| Hnrnpm | 74.5149 | 114.9792 |
| Pds5a | 8.012936222 | 12.36278747 |
| Erbb3 | 1.5017 | 2.31677 |
| Ick | 3.2962 | 5.084996 |
| Wbp11 | 29.5099395 | 45.49445 |
| Adamts17 | 0.99801 | 1.53822 |
| Bod1l | 1.76984381 | 2.727721 |
| Plekha5 | 7.60991883 | 11.72345678 |
| Arid4a | 1.83633121 | 2.82842072 |
| Plbd2 | 12.7413 | 19.6159 |
| Pacsin2 | 13.45537264 | 20.71476005 |
| Zfp280b | 4.638746 | 7.14070858 |
| Nr2c1 | 3.36693 | 5.1827 |
| Cxxc1 | 18.6918 | 28.7666 |
| Kantr | 1.598742234 | 2.460411231 |
| Mir17hg | 3.8004 | 5.84821 |
| Camkk2 | 5.555433894 | 8.54535477 |
| Spag5 | 3.47941 | 5.352 |
| Stxbp6 | 2.71678 | 4.17868 |
| Hdac7 | 14.313373 | 22.015383 |
| Klhl17 | 8.064524327 | 12.397678 |
| Tulp4 | 4.2328858 | 6.5071934 |
| Terf2ip | 3.37832 | 5.19268 |
| Ints3 | 17.44507 | 26.80767 |
| Armc7 | 2.07899 | 3.19468 |
| Nelfa | 19.8144 | 30.4401 |
| Uba7 | 1.92843 | 2.96217 |
| Sort1 | 4.319536 | 6.634699 |
| Sgms1 | 4.188219464 | 6.4328946 |
| Sgsh | 1.23632 | 1.89835 |
| Gga1 | 16.7317 | 25.6902 |
| Pick1 | 7.428123092 | 11.404436 |
| Zcchc7 | 5.769507 | 8.85744 |
| Tmem8 | 3.60365 | 5.53146 |
| Ltbp3 | 11.4322 | 17.5471 |
| R3hdm2 | 17.72702922 | 27.207777 |

| | | |
|---|---|---|
| Gns | 14.2282 | 21.8297 |
| Ikbke | 1.174525 | 1.802021117 |
| Dpp4 | 2.208006 | 3.3869796 |
| Nolc1 | 13.98123 | 21.44587 |
| Prr14l | 4.417308023 | 6.773761755 |
| Camk1g | 4.35029 | 6.67055 |
| Amotl1 | 5.360598001 | 8.218949 |
| Pura | 2.56388971 | 3.929354 |
| Nr4a1 | 2.900024895 | 4.444041 |
| Ttll5 | 1.865547 | 2.85839 |
| Myadm | 18.954474 | 29.03716 |
| Zfp652 | 4.561808748 | 6.987003059 |
| Abcd4 | 4.53947754 | 6.952632011 |
| Stk40 | 11.0874 | 16.978506 |
| Tet1 | 1.74973705 | 2.678906874 |
| Gtpbp1 | 14.59455 | 22.342183 |
| Hoxd8 | 12.25407478 | 18.75135 |
| Slc25a36 | 12.72330519 | 19.45734 |
| H2afy | 76.26707 | 116.61651 |
| Sos2 | 3.092859193 | 4.72902 |
| Cry1 | 15.02548554 | 22.968547 |
| Cnnm2 | 2.765774431 | 4.226297128 |
| Wnk1 | 12.19395691 | 18.632636 |
| Lss | 2.900617928 | 4.42918841 |
| Ascc2 | 5.582114 | 8.522531 |
| Cpne2 | 2.22326 | 3.3943 |
| Crip3 | 2.881628 | 4.39816 |
| Brd1 | 13.9071 | 21.22486 |
| Myof | 1.78916515 | 2.730442 |
| Zfp61 | 6.255755 | 9.54623 |
| Snx27 | 9.35766 | 14.27573 |
| Vangl1 | 3.47065 | 5.293943 |
| Tlr2 | 1.44684 | 2.20685 |
| Wnt2b | 4.432204 | 6.75981 |
| Dicer1 | 6.0670414 | 9.252578963 |
| Cyp2s1 | 6.95046 | 10.59891 |
| Il2rg | 1.604551719 | 2.4462797 |
| Hic1 | 29.12952 | 44.39419 |
| Susd6 | 5.433927 | 8.2811 |
| Lrfn1 | 6.945074917 | 10.58356193 |
| Cmklr1 | 0.958365 | 1.459653 |
| Vwa8 | 3.30009 | 5.02562 |
| Lmnb1 | 53.4963 | 81.4558 |
| Sf3b4 | 41.6708 | 63.4423 |
| Taf1c | 5.234 | 7.9675 |
| Hsd11b2 | 19.29635 | 29.37042719 |
| Sod3 | 0.692749 | 1.05389 |

| | | |
|---|---|---|
| Ralgapa2 | 0.893645 | 1.359134 |
| Cpne1 | 11.2745 | 17.14323 |
| Map4k4 | 27.55222415 | 41.887435 |
| Ezr | 14.7616364 | 22.43063902 |
| Col1a2 | 92.7556 | 140.944 |
| Clcf1 | 5.2544387 | 7.981691408 |
| Zfp146 | 17.8882 | 27.16536 |
| Tubgcp6 | 4.115264 | 6.24928 |
| Prex1 | 7.830528 | 11.89062 |
| Zmym4 | 7.760545705 | 11.783958 |
| Gcfc2 | 1.20302 | 1.82661 |
| Hmgb1-rs1 | 2.25644 | 3.42527 |
| Hcfc2 | 3.18366 | 4.83265 |
| Man2c1 | 10.0461 | 15.2436 |
| Mxi1 | 5.332072687 | 8.089438 |
| Trim44 | 15.73 | 23.8618 |
| Cwf19l1 | 3.63878135 | 5.518856917 |
| Eif4enif1 | 13.756567 | 20.86144 |
| Camsap3 | 5.21589561 | 7.90847 |
| Rapgef6 | 2.03597627 | 3.086453716 |
| Fgd2 | 1.342515 | 2.035108681 |
| Cpt1c | 7.504998 | 11.37476 |
| Arvcf | 8.7925833 | 13.32604131 |
| Isyna1 | 165.617 | 250.916 |
| Armc9 | 4.07003083 | 6.165310841 |
| Pes1 | 21.54531 | 32.631348 |
| Arfgef2 | 2.808248 | 4.252664445 |
| Lrp3 | 8.79013 | 13.310659 |
| Trim12c | 3.03037326 | 4.588607 |
| Tpbg | 6.429641367 | 9.73555 |
| Rhbdl3 | 2.685413704 | 4.0659955 |
| Espn | 5.056348 | 7.65413 |
| Iqgap2 | 0.911705 | 1.3801 |
| Foxm1 | 7.9155 | 11.9786 |
| Vav2 | 7.35927 | 11.13143 |
| Traf6 | 2.42595238 | 3.669196 |
| Crat | 7.7837 | 11.772257 |
| Ndor1 | 4.479807 | 6.774795 |
| Bcr | 6.41612 | 9.7028 |
| Ppm1h | 0.718559 | 1.08629 |
| Ccnj | 2.680498462 | 4.051947575 |
| Mapk6 | 13.56578096 | 20.5028 |
| Zcchc9 | 5.054419 | 7.63838 |
| Ankrd16 | 4.904648605 | 7.411702 |
| Hoxa7 | 4.133945865 | 6.24670204 |
| Letm2 | 1.16093771 | 1.753981 |
| Eif2ak4 | 3.334353 | 5.03743 |

| | | |
|---|---|---|
| Sptbn2 | 5.127315 | 7.743353 |
| Col9a3 | 0.666328 | 1.00629071 |
| Plagl2 | 6.3094 | 9.526 |
| Dnaaf3 | 4.09837 | 6.18718 |
| Tox | 0.910979 | 1.37493477 |
| Dhx57 | 4.46911125 | 6.744379 |
| Hnrnpl | 124.1514 | 187.3468 |
| Ankfy1 | 6.717294 | 10.13077 |
| Klf7 | 1.77749745 | 2.680707 |
| Poc5 | 3.05981 | 4.61436 |
| Tead3 | 12.30177 | 18.549436 |
| Mmp16 | 1.30249246 | 1.963655 |
| Erg | 0.719202372 | 1.084145036 |
| Zmym6 | 4.788554374 | 7.21720421 |
| Shroom1 | 0.830561 | 1.251694 |
| Cdon | 7.483739 | 11.273862 |
| Atm | 1.777923 | 2.67813 |
| Pnpla7 | 3.761624557 | 5.66549 |
| Slc38a1 | 6.806095112 | 10.249862 |
| Mknk2 | 39.9713763 | 60.1917 |
| Ddhd2 | 5.8598853 | 8.822395 |
| Sdc2 | 14.3259 | 21.5639 |
| Dgkq | 2.49155 | 3.74876 |
| Gal3st3 | 1.9787 | 2.97712 |
| Sap130 | 10.84594554 | 16.3126996 |
| Mipol1 | 1.53995 | 2.31611 |
| Chfr | 6.409364541 | 9.63940531 |
| Dach2 | 0.828371045 | 1.245720254 |
| Ctc1 | 19.65240103 | 29.550844 |
| Zfp384 | 14.44498 | 21.71444 |
| Usp31 | 1.606 | 2.41381 |
| Bmp6 | 1.65075 | 2.48078 |
| Myc | 7.4929977 | 11.25986 |
| Sh3bp2 | 3.638309906 | 5.466071 |
| Scaf8 | 9.6283 | 14.4649 |
| Mmp9 | 0.82986 | 1.24672 |
| Ddr1 | 25.33503806 | 38.0574011 |
| Gdf10 | 6.09142 | 9.14826 |
| Mturn | 2.83107522 | 4.25152157 |
| Taf2 | 5.569662 | 8.36207 |
| Bnc2 | 2.731170521 | 4.100470477 |
| Zfp185 | 2.564304128 | 3.849716019 |
| Mrc2 | 12.1766 | 18.2709 |
| Acp2 | 5.19154 | 7.78925 |
| Dnajc5 | 15.779554 | 23.674865 |
| Mical1 | 5.96949 | 8.95439 |
| Lrfn3 | 5.19438 | 7.78928 |

| | | |
|---|---|---|
| Asxl2 | 1.729559488 | 2.592984528 |
| Krt7 | 12.4601 | 18.6801 |
| Mtbp | 3.068945 | 4.600485 |
| Acin1 | 41.6848875 | 62.485256 |
| Zfhx3 | 1.17103 | 1.75529 |
| Tbce | 6.544551 | 9.80947 |
| Ppp1r10 | 8.19048027 | 12.2720263 |
| Atf6b | 23.4252 | 35.0883 |
| Kifc3 | 4.997019 | 7.48457 |
| Prr11 | 3.61516 | 5.4126 |
| Akap8 | 16.16611 | 24.19835 |
| Crebl2 | 1.02273 | 1.530836593 |
| Hinfp | 3.660505257 | 5.478667 |
| Podxl | 74.84695 | 112.0059 |
| Vps33a | 7.05816 | 10.56053 |
| Acss3 | 0.6457234 | 0.9660144 |
| Fam213a | 11.01536 | 16.47912 |
| Bdh1 | 1.62496 | 2.43023576 |
| Micalcl | 0.7595338 | 1.1357524 |
| Zfp592 | 4.599281 | 6.8751 |
| Ankrd10 | 20.58440661 | 30.76717758 |
| Selo | 2.48254 | 3.70901 |
| Fez1 | 2.53828 | 3.79216 |
| Zbtb8b | 2.0850453 | 3.114749 |
| Notch2 | 5.75607 | 8.59854 |
| Ccdc97 | 10.1457 | 15.1546 |
| Dnajb12 | 6.098215468 | 9.10888 |
| Trim3 | 6.464233574 | 9.6539418 |
| Etv6 | 5.311398 | 7.930215471 |
| Kdsr | 2.17078 | 3.24109 |
| Rabep2 | 6.50348 | 9.70806 |
| Ralgps1 | 1.019211237 | 1.521342258 |
| Phf12 | 12.46215 | 18.59857 |
| Abl2 | 3.7607548 | 5.612159314 |
| Boc | 9.6430703 | 14.389076 |
| C1ra | 2.32851 | 3.473692 |
| Usp43 | 0.880192 | 1.312978067 |
| Ddx27 | 13.105 | 19.5475 |
| Idua | 1.3931175 | 2.077921 |
| Cd14 | 1.07886 | 1.60894 |
| Fam78b | 1.26267621 | 1.882328 |
| Ptk2 | 9.9936425 | 14.8965777 |
| Clec2d | 1.497509487 | 2.2321732 |
| Ccnk | 10.932658 | 16.29287065 |
| Zyx | 36.695724 | 54.679003 |
| Lrp6 | 6.83654 | 10.18651 |
| Has2 | 1.79697 | 2.67693 |

| | | |
|---|---|---|
| Golim4 | 4.71539592 | 7.02302487 |
| Acvrl1 | 3.729748296 | 5.552722868 |
| Trak1 | 6.413147 | 9.54662296 |
| Zfp579 | 16.09014 | 23.94692 |
| Fam101a | 0.943674 | 1.404409 |
| Mib2 | 10.54120145 | 15.68656357 |
| Rprd1b | 7.429532 | 11.05490119 |
| Ndrg2 | 6.583651205 | 9.796057 |
| Cecr2 | 2.478445388 | 3.6876563 |
| Xylt1 | 0.937354 | 1.39466 |
| Rbm33 | 4.37832 | 6.51326 |
| Gemin5 | 7.03264127 | 10.4610727 |
| Tmem86b | 1.725735 | 2.56654 |
| Tbrg4 | 15.3321 | 22.7939883 |
| Irf5 | 4.79642 | 7.12983 |
| Zc3h14 | 15.67133338 | 23.29166204 |
| Col16a1 | 10.7899 | 16.031112 |
| Usp20 | 4.283017 | 6.36319 |
| Gm17762 | 1.60786 | 2.38812 |
| Mfhas1 | 2.267980971 | 3.36754 |
| Gab1 | 13.034172 | 19.3512885 |
| Hnrnph3 | 20.7444552 | 30.79828 |
| Mon2 | 4.7322192 | 7.024626362 |
| Pard6b | 4.107757 | 6.096874571 |
| Tns3 | 27.82309 | 41.2903 |
| D930015E( | 8.04916137 | 11.9434867 |
| Hook3 | 2.32946 | 3.45607 |
| Cdca7l | 5.641784 | 8.3699 |
| Dot1l | 12.30172 | 18.24950897 |
| Atp13a1 | 13.00114 | 19.28697 |
| Eno1 | 295.9374 | 438.9927 |
| Grhl2 | 0.725907 | 1.076714 |
| Parp1 | 51.35179 | 76.1574 |
| Gsr | 15.3421 | 22.7527 |
| Opa1 | 5.520303995 | 8.185421368 |
| Aff1 | 2.477355894 | 3.672201 |
| Lhfp | 16.8921 | 25.0353 |
| Clcn2 | 8.33898 | 12.3586 |
| Zak | 1.1307429 | 1.67572 |
| Arid3a | 3.76576 | 5.580444 |
| Kif3b | 3.3713 | 4.99456 |
| Rtn4r | 1.25675 | 1.86184 |
| Xab2 | 17.5437 | 25.9872 |
| Notch4 | 1.33308 | 1.97346 |
| Zfp747 | 1.41701 | 2.09739 |
| Sh3d21 | 3.09438302 | 4.579603738 |
| Rassf1 | 12.1806 | 18.02463 |

| | | |
|---|---|---|
| Golga3 | 4.591937347 | 6.794993101 |
| Sbk1 | 37.13997 | 54.95693 |
| Cog5 | 3.04369 | 4.50382 |
| Osbpl6 | 0.87333441 | 1.292132912 |
| Sall1 | 1.27281919 | 1.88315525 |
| Clk2 | 5.89959 | 8.72769 |
| Nacc1 | 23.0646 | 34.12104 |
| Usp6nl | 3.77108044 | 5.57857552 |
| Ror2 | 6.28549 | 9.2973 |
| Gm53 | 2.054452 | 3.03838 |
| Podxl2 | 9.84399 | 14.55615 |
| Rfng | 9.532272956 | 14.094843 |
| Fscn1 | 100.1126 | 148.0224 |
| Lrp12 | 3.61453 | 5.34424 |
| Fktn | 2.2302946 | 3.297392677 |
| Flcn | 10.83412716 | 16.01565 |
| Ddhd1 | 2.540674373 | 3.755539057 |
| Mbd1 | 11.29371118 | 16.69381728 |
| Aak1 | 4.132557282 | 6.107767145 |
| Ehbp1 | 2.598358819 | 3.83970632 |
| Gps2 | 14.84257 | 21.92727 |
| Adat2 | 8.00387 | 11.8195 |
| Acap3 | 4.96662 | 7.3341 |
| Ogfod1 | 2.183969 | 3.224944899 |
| Mmp17 | 1.68318763 | 2.485300846 |
| Lrrc8d | 5.61000733 | 8.28199118 |
| Hoxc10 | 3.57294 | 5.27461 |
| Srf | 13.18512 | 19.46472 |
| Hira | 11.828102 | 17.46135 |
| Celf3 | 0.97360329 | 1.436732412 |
| Zfyve27 | 3.761148049 | 5.548961339 |
| Lpar6 | 5.25415 | 7.75045 |
| Kdm2b | 19.394109 | 28.60695 |
| Kdm8 | 1.999669 | 2.949524 |
| Epb4.1l2 | 30.10984 | 44.4109733 |
| Zfp296 | 2.07895 | 3.06597 |
| Casc3 | 14.7936 | 21.8133 |
| Ero1l | 7.28229 | 10.7362 |
| Arhgef7 | 11.76751 | 17.3464348 |
| Slc39a13 | 6.47624137 | 9.543701711 |
| Ube3b | 19.21486 | 28.31573 |
| Fam53b | 4.442049132 | 6.544653 |
| Adarb1 | 1.104660653 | 1.62750544 |
| Rcsd1 | 10.75962678 | 15.85049 |
| Fam214b | 4.359804909 | 6.421483577 |
| Tcf3 | 64.816058 | 95.45575 |
| Vldlr | 1.738333632 | 2.558918 |

| | | |
|---|---|---|
| Arhgef4 | 1.661501 | 2.44575 |
| Fxr2 | 25.1133 | 36.9669 |
| Plscr3 | 18.203845 | 26.795145 |
| Fam21 | 9.16897 | 13.49608 |
| Scyl1 | 14.8299 | 21.8284 |
| Zbtb4 | 1.31974 | 1.94248 |
| Camk2n2 | 2.11582 | 3.11379 |
| Kif14 | 1.17984602 | 1.7361718 |
| Rbm45 | 10.8266 | 15.9251 |
| Lrrc56 | 3.43415 | 5.0500019 |
| Ciart | 1.75119299 | 2.575161743 |
| Slc25a37 | 4.082872 | 6.00335 |
| Eif2ak1 | 10.74757 | 15.80293 |
| Nppc | 0.820606 | 1.20656 |
| Prcc | 27.37247 | 40.24208 |
| Foxo6 | 4.4071 | 6.4791 |
| Mettl17 | 6.97843 | 10.2581 |
| Pros1 | 4.27803 | 6.28796 |
| Cmtm4 | 5.72104 | 8.40828 |
| Scube3 | 5.734543 | 8.427528775 |
| Ggcx | 4.10533 | 6.03282 |
| Adgrf5 | 1.80523208 | 2.652693926 |
| Doc2g | 2.122487 | 3.118493 |
| Prkci | 10.8646 | 15.9626 |
| Slc23a2 | 4.6842373 | 6.881713 |
| Pde9a | 2.567129242 | 3.771203232 |
| Plscr1 | 8.652316 | 12.709795 |
| Ankrd26 | 1.2950484 | 1.901962 |
| Trit1 | 5.013420862 | 7.3629078 |
| Fbxl3 | 5.8594363 | 8.60513 |
| Cbs | 1.05567986 | 1.550316169 |
| Mtf1 | 1.58865 | 2.33284 |
| Rfx3 | 1.024638157 | 1.50456138 |
| Ogt | 25.39116 | 37.2827 |
| Coro7 | 5.32786 | 7.82097 |
| Brd8 | 8.331 | 12.226754 |
| Pcsk6 | 8.19996 | 12.03332 |
| Gmeb1 | 3.958024 | 5.80828 |
| Dcbld2 | 6.60832 | 9.696164 |
| Gm11110 | 1.509437 | 2.214597111 |
| Ppfibp2 | 4.06012 | 5.956275 |
| Stk38 | 14.7211 | 21.59104555 |
| Smcr8 | 2.5078962 | 3.678085 |
| Pkmyt1 | 8.35456 | 12.25263 |
| Fgfr1 | 21.1988217 | 31.08736368 |
| Grm4 | 1.319766247 | 1.935188772 |
| Gnl2 | 12.9305 | 18.9536 |

| | | |
|---|---|---|
| Lrrc71 | 2.44354 | 3.58064 |
| Fam222b | 4.82234 | 7.06607 |
| Lnx2 | 2.468080881 | 3.61577 |
| Zfp46 | 3.491995 | 5.11557 |
| Smarcal1 | 3.927946 | 5.753421476 |
| Snrnp70 | 103.3744 | 151.36556 |
| Aatf | 14.352986 | 21.015178 |
| Aldh1l1 | 1.09983747 | 1.610291188 |
| Bend3 | 3.186866538 | 4.66528 |
| Tjp2 | 6.68544186 | 9.78586 |
| Prkcsh | 40.1208 | 58.72482 |
| Plcl1 | 1.68865 | 2.47134 |
| Taok3 | 1.246361365 | 1.823975392 |
| Cep295 | 1.9605853 | 2.868907 |
| Dnajc1 | 2.671778 | 3.90953 |
| Ipo13 | 10.5727 | 15.468727 |
| C1ql1 | 2.09473 | 3.06457 |
| Zfp334 | 5.29449 | 7.74577 |
| Chd1 | 3.64754 | 5.33515 |
| Mxd1 | 2.89727 | 4.235761824 |
| Particl | 6.10559 | 8.92471 |
| Mgat3 | 7.569266 | 11.06362 |
| Arid5b | 5.32258995 | 7.778506344 |
| Rnf145 | 17.8579285 | 26.097647 |
| Il17rd | 3.0099535 | 4.39797 |
| Abca7 | 1.101305 | 1.60902 |
| Csnk1e | 20.03799919 | 29.27327029 |
| Trub1 | 6.995517 | 10.21829 |
| Slc43a2 | 1.430109279 | 2.08811918 |
| Drp2 | 3.453655 | 5.041915 |
| Ankrd45 | 1.040726465 | 1.519171 |
| Kcnh2 | 9.48420288 | 13.843775 |
| Ctr9 | 9.89247 | 14.4396 |
| Col25a1 | 0.84959359 | 1.240112877 |
| Hsf1 | 14.97729 | 21.84819 |
| Coq5 | 9.52421 | 13.8918 |
| AW549877 | 8.49065 | 12.38371 |
| Dhtkd1 | 2.386106 | 3.480106 |
| Luzp1 | 4.606928949 | 6.717080427 |
| Ttc7 | 1.644706 | 2.39801 |
| Nr2f2 | 31.52291 | 45.95858 |
| Wdr70 | 4.89148 | 7.12943 |
| Exosc10 | 15.88404 | 23.15060349 |
| Gaa | 14.46285056 | 21.07755 |
| Zfp503 | 33.619302 | 48.98743 |
| Nup155 | 6.6015014 | 9.61802404 |
| Vars | 31.2126 | 45.4643 |

| | | |
|---|---|---|
| Adgrg1 | 13.76337 | 20.04423909 |
| Nfatc4 | 12.93882 | 18.84258 |
| Cry2 | 3.49764 | 5.09276 |
| Galnt12 | 1.976103 | 2.87731 |
| Gfpt1 | 7.20382 | 10.48880735 |
| Ubr3 | 4.96018 | 7.222049 |
| Svep1 | 1.68373 | 2.451442 |
| Dyrk2 | 4.40251 | 6.409836 |
| Ftx | 1.506516572 | 2.19298809 |
| Atxn1l | 4.52472 | 6.585748 |
| Krit1 | 2.0565635 | 2.9933055 |
| Zeb1 | 7.360803763 | 10.71199716 |
| Hdac5 | 17.58276 | 25.5855709 |
| Usp13 | 0.971480191 | 1.41268269 |
| Vrk1 | 6.205548 | 9.022711 |
| Gripap1 | 4.199978682 | 6.106225 |
| Col13a1 | 6.942131485 | 10.09277282 |
| Mybl2 | 17.58233212 | 25.55367 |
| Map1b | 2.5878 | 3.76078 |
| Ampd2 | 14.56712436 | 21.16777 |
| Dfna5 | 1.39694 | 2.02985 |
| Tnfrsf22 | 0.774829 | 1.1251 |
| Mrvi1 | 2.1111551 | 3.065053 |
| Ptgfrn | 13.59565 | 19.7350419 |
| Klc3 | 2.518819968 | 3.65552 |
| Tbc1d25 | 6.401236 | 9.28835 |
| Guf1 | 3.477903 | 5.046196 |
| Actn1 | 15.91808 | 23.09391 |
| Stox2 | 1.866189646 | 2.707079268 |
| Enox1 | 2.10638698 | 3.0552965 |
| Smpdl3b | 17.42307 | 9.576284 |
| Gpx7 | 59.9927 | 32.9592 |
| Tomm7 | 184.036 | 101.093 |
| Cdk2ap2 | 54.147 | 29.7391 |
| Asnsd1 | 63.7611 | 35.01478 |
| Ccdc90b | 38.116603 | 20.929415 |
| Mxra7 | 13.9715 | 7.67009 |
| Nap1l1 | 725639.7073 | 398315.8632 |
| Vamp5 | 8.610141 | 4.72591 |
| Grtp1 | 9.46226 | 5.19248 |
| Hagh | 19.60040014 | 10.74798267 |
| Sh3bgrl | 51.8807 | 28.4405 |
| Ict1 | 96.2629 | 52.7628 |
| Rpl8 | 5182.44 | 2840.36 |
| AU022751 | 3.801684 | 2.08302 |
| Acsbg1 | 13.47706 | 7.38249 |
| Rwdd1 | 83.205 | 45.5772 |

| | | |
|---|---|---|
| Pycard | 7.89992 | 4.326 |
| Pop1 | 6.54895 | 3.58579 |
| Cldn5 | 54.6133 | 29.8885 |
| Pla2g1b | 1.49043 | 0.815586 |
| Wfdc1 | 16.71982 | 9.149054865 |
| Ovca2 | 16.4143 | 8.98145 |
| Uqcc3 | 81.1537 | 44.3949 |
| Prss36 | 15.52275143 | 8.4905112 |
| Vdac3 | 275.62983 | 150.6724 |
| Timm22 | 35.439836 | 19.37071333 |
| Mrpl18 | 90.7583 | 49.5947 |
| Tmem242 | 143.301 | 78.2868 |
| Fabp3 | 17.7845 | 9.71482 |
| Cdkl1 | 1.34141 | 0.732646 |
| Hmgcs2 | 49.41516593 | 26.989189 |
| Taf9 | 112.97822 | 61.65708 |
| Cklf | 13.5569376 | 7.397215196 |
| Angpt4 | 2.58073 | 1.4081344 |
| Scml2 | 3.145917851 | 1.716482213 |
| Xpa | 14.29152 | 7.79429 |
| Rpl6 | 2707.1219 | 1476.34296 |
| Nudt1 | 46.4261 | 25.3071 |
| Mixl1 | 1.00079 | 0.5454 |
| Trib3 | 7.35084 | 4.00581295 |
| Trappc2l | 63.1547 | 34.4145 |
| Emc6 | 67.252207 | 36.638428 |
| Rnf128 | 15.763908 | 8.586571 |
| Hist1h4b | 12.9864 | 7.07291 |
| Tmem251 | 12.02446 | 6.54712 |
| Zfp934 | 2.8750246 | 1.56523632 |
| Psma4 | 231.483 | 126.024 |
| Avpi1 | 25.0481 | 13.6354 |
| Tmem42 | 14.6962 | 7.99712 |
| Ndufv3 | 201.42654 | 109.6031 |
| Mbip | 12.3854 | 6.7368 |
| Ccdc28b | 30.9634 | 16.8382 |
| Itgb1bp1 | 17.61233 | 9.5751 |
| Naa15 | 25.62883095 | 13.93332375 |
| Itgb6 | 0.690403063 | 0.375329483 |
| Pctp | 8.04531 | 4.37354848 |
| Mettl22 | 14.2604047 | 7.751526 |
| Cgrrf1 | 15.70047 | 8.53229568 |
| Zfp867 | 4.223883 | 2.293963 |
| Zc3h12b | 2.174375 | 1.1806901 |
| Nagk | 31.75976 | 17.23551 |
| Zfp791 | 1.82279 | 0.988976 |
| Mrpl49 | 42.0965 | 22.8388 |

| | | |
|---|---|---|
| Phpt1 | 244.669 | 132.729 |
| Ntf5 | 5.70679368 | 3.094702073 |
| Nudt21 | 181.244 | 98.1823 |
| Cops6 | 297.277 | 161 |
| Gamt | 62.9077 | 34.063 |
| Ackr1 | 1.32754 | 0.718558 |
| Yif1a | 45.3211 | 24.5293 |
| Cox5b | 574.565 | 310.964 |
| Tmem160 | 155.2684 | 84.0198 |
| Pusl1 | 21.4606 | 11.6127 |
| Npm3 | 188.874 | 102.177 |
| Tmem9b | 20.4509 | 11.0612 |
| Hyi | 38.82932 | 20.97373078 |
| Commd6 | 41.73093 | 22.53523 |
| Crls1 | 29.5296 | 15.94401505 |
| Asns | 47.0201 | 25.3873 |
| Emc8-1190 | 27.08403 | 14.6218048 |
| Glrx3 | 222.5127 | 120.1147 |
| Tmem29 | 15.652146 | 8.4484641 |
| Mthfd2l | 12.03435 | 6.49463 |
| Atp5g1 | 464.61831 | 250.695971 |
| Rnd2 | 90.5812 | 48.85923 |
| Rpl37a | 5692.37 | 3068.52 |
| Rps3a1 | 3784.19 | 2039.81 |
| Eif4ebp1 | 127.22 | 68.574 |
| Etnk2 | 61.2683 | 33.01533 |
| Fam210b | 11.2717 | 6.07214 |
| Rapsn | 1.17214 | 0.631402 |
| Nfu1 | 59.63403 | 32.09656 |
| Ift27 | 40.7837 | 21.950278 |
| Tceb2 | 948.621 | 510.554 |
| Arhgap5 | 12.134067 | 6.529681 |
| Gstk1 | 25.3857 | 13.66041 |
| Cenpi | 9.7376 | 5.239771 |
| Tbcb | 99.2434 | 53.3854 |
| Tusc2 | 38.2736 | 20.5836 |
| Kbtbd8 | 1.302186489 | 0.700274 |
| Anapc13 | 199.591 | 107.332 |
| Gde1 | 62.225 | 33.45403 |
| Vbp1 | 58.5765 | 31.4628 |
| Mgst1 | 146.415 | 78.6138 |
| C77370 | 1.350602294 | 0.72512 |
| Lamtor1 | 174.98 | 93.9382 |
| Art1 | 0.97474409 | 0.523255803 |
| Mea1 | 130.75416 | 70.14664 |
| Ubr5 | 40.5515926 | 21.74813199 |
| Ppp4c | 214.241 | 114.887 |

| | | |
|---|---|---|
| Med11 | 34.6919 | 18.6033 |
| Smim12 | 60.869 | 32.6208 |
| Colec12 | 80.2288 | 42.996 |
| Emc7 | 49.7494 | 26.6586 |
| Prpf4b | 659914.2687 | 353494.9762 |
| Esd | 141.465799 | 75.74369 |
| Zfp672 | 16.677209 | 8.927615 |
| Hras | 44.657 | 23.894223 |
| Jade3 | 12.37254 | 6.615273358 |
| Cks1b | 229.97 | 122.953 |
| Pacrgl | 14.03391 | 7.50145493 |
| Vps29 | 69.4205 | 37.097 |
| Rbmxl2 | 21.2941 | 11.3678 |
| Actr10 | 20.8724 | 11.133907 |
| Gm13242 | 1.12824 | 0.601818 |
| Klhl15 | 4.748383 | 2.532513 |
| Uqcrb | 352.019 | 187.699 |
| Polr3k | 23.2451 | 12.3852 |
| Cxcl14 | 1.85883 | 0.990221 |
| Cyba | 148.87535 | 79.24877 |
| Arl5c | 2.42265 | 1.28949 |
| Tm2d2 | 94.9877 | 50.5519 |
| Psmb7 | 340.809 | 181.37 |
| Ssu2 | 1.256315 | 0.668513129 |
| Il13ra1 | 3.82347 | 2.03429 |
| Fbxo3 | 930482.5545 | 495042.3563 |
| Mrgpra6 | 39.2988 | 20.9057 |
| Snrpc | 264.098 | 140.489 |
| Mgarp | 2.97292097 | 1.58098504 |
| Wbp5 | 316.851 | 168.458 |
| Immp1l | 80.1996 | 42.6314 |
| Icam4 | 3.41135 | 1.813045 |
| Thbs2 | 2.87409 | 1.52692 |
| Phf7 | 9.572654542 | 5.083897 |
| Myl1 | 13.23518242 | 7.028688 |
| Txnl4b | 9.843 | 5.22697 |
| Asf1a | 24.390521 | 12.9521 |
| Trim59 | 32.8044 | 17.41989 |
| Bag1 | 176.642 | 93.7828 |
| Krtcap2 | 401.589 | 213.162 |
| Spp2 | 2.40412 | 1.27604 |
| Foxc1 | 5.0387387 | 2.673643 |
| Echdc3 | 4.93856 | 2.6193 |
| Fkbp2 | 74.7187 | 39.611 |
| Lppos | 6.04779 | 3.20568 |
| Adad2 | 3.37714 | 1.790066 |
| Plekhf1 | 10.6466 | 5.64263 |

| | | |
|---|---|---|
| Bud31 | 84.24851 | 44.648228 |
| Cops5 | 105.64986 | 55.988673 |
| Rps4x | 4945.37 | 2620.16 |
| Gm6654 | 19.124 | 10.1317 |
| Orc4 | 13.36335 | 7.079225 |
| Manf | 62.6848 | 33.20661 |
| Gbx2 | 2.17909 | 1.15434 |
| Gpat2 | 12.828599 | 6.7927362 |
| Tm2d3 | 72.50937 | 38.37687 |
| Cyc1 | 317.909 | 168.173 |
| Lamtor5 | 100.689 | 53.2563 |
| Znrd1as | 5.89823 | 3.118803639 |
| Taf1d | 36.26257 | 19.1721 |
| Sult1d1 | 1.22377 | 0.646934 |
| Rmst | 2.494139834 | 1.318434 |
| Cdkl3 | 2.098466223 | 1.108942 |
| Dpy30 | 147.572812 | 77.98145 |
| Lysmd2 | 19.411651 | 10.25754 |
| Fam187b | 1.82806 | 0.965772 |
| Pop7 | 39.99 | 21.1252 |
| Haus4 | 35.72346 | 18.86911 |
| Creg1 | 32.7244 | 17.285 |
| Timp1 | 55.37809 | 29.24812 |
| Hsd17b10 | 216.256 | 114.213 |
| Chp2 | 1.99448 | 1.053297 |
| B3galnt1 | 11.18200993 | 5.903827328 |
| Mblac1 | 6.45596 | 3.40791 |
| Gm13889 | 8.34376 | 4.40109 |
| Bcl7c | 101.447 | 53.4564 |
| Rpl36al | 884.497 | 466.048 |
| Gm826 | 2.95371 | 1.5562 |
| Spryd7 | 8.692798242 | 4.57983 |
| Mrps23 | 54.48846 | 28.703207 |
| Mrpl22 | 53.9507 | 28.4137 |
| Slc25a4 | 811.207 | 426.903 |
| Zfp942 | 5.048836 | 2.656887 |
| Snx14 | 16.33054 | 8.58936 |
| Fam134b | 2.250145512 | 1.183455517 |
| Fam57b | 1.465053107 | 0.769746391 |
| Naca | 1394.10701 | 732.456538 |
| Mrpl15 | 49.826141 | 26.1743177 |
| Prelid2 | 25.6909 | 13.4948 |
| Rab3d | 4.617604 | 2.425433753 |
| Il1f9 | 1.40993 | 0.739945 |
| Ppap2c | 36.60583815 | 19.21047 |
| Nup43 | 49.0079 | 25.7143 |
| Tpd52l1 | 28.22304 | 14.80732 |

| | | |
|---|---|---|
| Cisd1 | 61.6138 | 32.3206 |
| Fundc2 | 118.369567 | 62.0922 |
| Lonp2 | 27.97456 | 14.66976 |
| Gnat2 | 1.537759 | 0.806366 |
| Mrpl55 | 50.38206822 | 26.41575 |
| BC030336 | 16.55622116 | 8.6786 |
| Churc1 | 87.9814 | 46.1165 |
| Thg1l | 11.56666324 | 6.0621 |
| Erp29 | 182.107 | 95.436 |
| Ccdc167 | 12.785681 | 6.6973 |
| Chchd7 | 84.2562929 | 44.12406879 |
| Pla1a | 1.6297753 | 0.853489 |
| Tmem86a | 60.63383 | 31.74900263 |
| Tspyl5 | 3.57404 | 1.87073 |
| Zfp954 | 6.9755 | 3.64907 |
| Abhd17a | 215.282 | 112.614 |
| Atox1 | 289.7047 | 151.4804 |
| Mcee | 41.4812 | 21.68694 |
| Use1 | 257.2624 | 134.48721 |
| Stamos | 3.0677 | 1.60365 |
| Thoc7 | 97.76518 | 51.08878 |
| Jpx | 2.012127 | 1.050964 |
| Adra2c | 0.750873 | 0.392061 |
| Rps19 | 4907.706 | 2561.132 |
| Ddx25 | 11.97891 | 6.2508 |
| Psmg3 | 50.18529 | 26.18374 |
| Morn2 | 22.8581 | 11.9234 |
| Nsmce1 | 110.823 | 57.8029 |
| Mpv17l2 | 31.363 | 16.3461 |
| Tmem47 | 8.89886 | 4.63739 |
| Psmb6 | 455.706 | 237.395714 |
| Gm9 | 2.36908 | 1.23363 |
| Pcbd2 | 53.0921 | 27.6411 |
| Slc18a2 | 2.34639 | 1.22107 |
| Mrps35 | 77.2785 | 40.1747 |
| Coa4 | 25.7892 | 13.4049 |
| H3f3a | 1471.627 | 764.7930083 |
| Sdhaf3 | 22.47997 | 11.680934 |
| Alox12 | 0.678852 | 0.352732 |
| Dusp14 | 4.88437799 | 2.537920469 |
| Gins2 | 67.4477 | 35.0317 |
| Ydjc | 14.50031 | 7.530577 |
| Ppp1r11 | 34.0261 | 17.6642 |
| Tst | 116.054 | 60.2361 |
| Mpped1 | 2.04924469 | 1.063419245 |
| Apool | 47.25826 | 24.52316434 |
| Naa20 | 67.98663 | 35.2741 |

| | | |
|---|---|---|
| D8Ertd738e | 241.151 | 125.086 |
| Zfp758 | 2.1263237 | 1.10246921 |
| Nqo1 | 11.631293 | 6.03032 |
| Zfp748 | 5.1359 | 2.66187 |
| Cutc | 10.62649 | 5.502343 |
| Spcs1 | 172.549 | 89.3303 |
| Nudt14 | 78.837 | 40.78226 |
| Pantr1 | 2.77893 | 1.43701 |
| BC004004 | 25.34228083 | 13.09697729 |
| Gm5801 | 64.51344 | 33.33202 |
| S100a6 | 72.2196 | 37.3117 |
| Cisd3 | 23.99472 | 12.39174857 |
| Yipf5 | 32.6176 | 16.8394 |
| Bace2 | 8.77654 | 4.529731 |
| Cops2 | 50.89305 | 26.25205 |
| Ndufa3 | 561.622 | 289.648 |
| Rpl23 | 1840.43963 | 949.12092 |
| Eif3i | 427.226 | 220.237 |
| Slc1a5 | 5.86431 | 3.02277 |
| Cacybp | 75.265844 | 38.785 |
| Rps6 | 3689.76 | 1899.341 |
| Qdpr | 105.6019084 | 54.31849 |
| Gng8 | 4.123947 | 2.120841948 |
| Rsph9 | 20.30645 | 10.43744 |
| Zfp386 | 6.479089332 | 3.329224 |
| Cnpy2 | 238.2727 | 122.3699 |
| Thap4 | 67.1087 | 34.45994 |
| Stra13 | 98.8198 | 50.73455 |
| Bub1 | 27.1081 | 13.916895 |
| Timm8a1 | 93.3864 | 47.9353 |
| Rhoc | 222.42047 | 114.119 |
| Arhgap24 | 3.858829421 | 1.979529 |
| Fth1 | 1574.85467 | 807.875 |
| Map1lc3a | 109.988 | 56.4171 |
| Coa3 | 162.859 | 83.5206 |
| Tomm20 | 78.8808651 | 40.4485627 |
| Phospho2 | 11.31227 | 5.8000864 |
| Aurkaip1 | 205.6169 | 105.4103 |
| Ggnbp2os | 7.85667 | 4.02579 |
| Psmb3 | 382.21 | 195.612 |
| Ndufv2 | 128.417348 | 65.696792 |
| Adhfe1 | 0.73992856 | 0.378370454 |
| Ube2v2 | 9.292638 | 4.750748 |
| Rpl10 | 4610.4 | 2354.52 |
| Rplp2-ps1 | 137.966 | 70.3852 |
| Cript | 56.198 | 28.6637 |
| Psma2 | 281.621 | 143.58 |

| | | |
|---|---|---|
| Hspa4l | 2.290454 | 1.166992 |
| Rpl24 | 2820.79 | 1436.25 |
| Nudt17 | 2.035184 | 1.036067 |
| Zglp1 | 16.0179 | 8.15168 |
| Ubb | 757.229 | 385.259 |
| Clybl | 20.8980854 | 10.63054666 |
| Fam35a | 78.65326594 | 40.00649862 |
| Wbp1 | 147.4427 | 74.98148 |
| Polr2g | 204.761 | 104.092 |
| Mtmr7 | 5.1385 | 2.61189 |
| Dph3 | 27.05327702 | 13.75077382 |
| Gemin7 | 119.866 | 60.91813 |
| Trnau1ap | 51.7555 | 26.2973 |
| Fkbp6 | 45.09155 | 22.89641086 |
| Lpar4 | 3.2628 | 1.65662 |
| Adarb2 | 9.87978 | 5.01378636 |
| Tubb3 | 6.85989 | 3.47924 |
| Xbp1 | 64.12177 | 32.51675 |
| Nt5c | 56.4008 | 28.5967 |
| Itpa | 91.3852 | 46.3191 |
| Macrod2 | 2.2079527 | 1.1190888 |
| Zfp456 | 1.597993326 | 0.809777 |
| Mettl23 | 31.84884 | 16.12347 |
| Mrps28 | 92.5958 | 46.8465 |
| Mrps10 | 45.0825 | 22.795768 |
| Ndufa1 | 205.042 | 103.634 |
| Aimp1 | 126.891 | 64.1015 |
| Gmfg | 10.99985 | 5.555411 |
| Alkbh7 | 58.50709 | 29.542204 |
| Gsta3 | 3.528399908 | 1.781305896 |
| Tmem208 | 106.251 | 53.6146 |
| Lgals7 | 526.38388 | 265.569221 |
| Caml | 33.4448 | 16.8721 |
| Sdpr | 4.09309 | 2.06424 |
| Eva1a | 3.21571 | 1.621544 |
| Rpl3 | 3569.62 | 1799.85 |
| Rplp0 | 4916.03 | 2478.63 |
| Cox6b1 | 797.703 | 402.159 |
| Mrpl12 | 202.712 | 102.192 |
| Smim4 | 79.58774 | 40.114631 |
| Sdhd | 191.09 | 96.2561 |
| Gm5141 | 1.40317 | 0.706673 |
| Siah1b | 31.13345 | 15.67511 |
| Bri3 | 164.297141 | 82.699 |
| Pdcd10 | 31.0529 | 15.615589 |
| Mrpl34 | 199.225 | 100.162 |
| Cox17 | 142.276 | 71.5026 |

| | | |
|---|---|---|
| Hist1h2ac | 2.31453 | 1.16277 |
| Ddc | 6.8456 | 3.437888234 |
| Polr1d | 257.6864 | 129.3927 |
| C330013J2 | 3.24903 | 1.6311 |
| Defb19 | 10.3932 | 5.21766 |
| Klhl35 | 1.2779 | 0.641243 |
| Timm17b | 88.29788 | 44.26352 |
| Ddrgk1 | 61.4325 | 30.7678 |
| Gm16740 | 7.55298 | 3.78195 |
| Mrpl27 | 84.3947 | 42.2471 |
| Anxa9 | 1.4053211 | 0.703472 |
| Sdhb | 206.023 | 103.09 |
| Pts | 36.54027 | 18.27974 |
| Dgcr6 | 64.032341 | 32.01038 |
| Nme3 | 70.9646 | 35.4521 |
| Rasl10b | 4.751589787 | 2.373493135 |
| Eid1 | 172.319 | 86.0739 |
| Cited4 | 9.54938 | 4.76962 |
| Mrpl41 | 23.022631 | 11.49794 |
| Prorsd1 | 6.97402 | 3.482515 |
| Ostc | 228.046 | 113.874 |
| Tm2d1 | 35.99519 | 17.96219 |
| Wfdc13 | 0.988947 | 0.493421 |
| Htatip2 | 6.109326468 | 3.047379824 |
| Sec22a | 10.279586 | 5.125693 |
| Timm23 | 215.555 | 107.395 |
| Fkbp7 | 39.36983 | 19.60219 |
| Fam174a | 13.7 | 6.82091 |
| Zfp850 | 1.923021061 | 0.95623825 |
| Chac2 | 3.23535 | 1.607521 |
| Hddc3 | 71.8679 | 35.7075 |
| Prss35 | 4.731016 | 2.3498444 |
| Anapc11 | 41.4551 | 20.58344 |
| Ssu72 | 90.3006 | 44.8322 |
| Metrn | 92.0358 | 45.6807 |
| Chchd6 | 56.18765 | 27.88634 |
| Defb36 | 0.719117 | 0.356878 |
| Guk1 | 109.6969842 | 54.4379861 |
| Pafah1b3 | 283.556 | 140.6514 |
| Gata1 | 1.3053 | 0.647387 |
| Dguok | 37.6621 | 18.67462 |
| Esco1 | 10.367795 | 5.138687 |
| Psenen | 210.502 | 104.321 |
| Btf3 | 1187.88604 | 588.64774 |
| Psma6 | 277.147 | 137.255 |
| Hrsp12 | 31.9365 | 15.8122 |
| Mia | 2.4078 | 1.19211 |

| | | |
|---|---|---|
| Wfdc6a | 3.223451485 | 1.595093828 |
| Hist1h4j | 2.442175 | 1.208006 |
| Ccdc61 | 53.61574 | 26.515325 |
| Grcc10 | 619.643 | 306.418 |
| Ptpn18 | 6.330378 | 3.130315 |
| Mrpl13 | 137.878 | 68.1656 |
| Pxmp2 | 16.8086 | 8.30918 |
| Lrp11 | 3.192201353 | 1.576851061 |
| Ppib | 329.75 | 162.851 |
| Chchd3 | 131.5328 | 64.9403 |
| Eif3h | 678.28 | 334.7992 |
| Zfp133-ps | 0.764345 | 0.37728 |
| Atp5c1 | 471.94424 | 232.943961 |
| Swi5 | 338.63198 | 167.128169 |
| Psmg1 | 103.065 | 50.8564 |
| Fam131c | 1.07348 | 0.529668 |
| Zdhhc2 | 15.217 | 7.50781 |
| Zfp820 | 1.8454844 | 0.91049822 |
| Sin3b | 326.31585 | 160.93992 |
| Aunip | 11.201262 | 5.52394 |
| Pilra | 4.82668 | 2.3791 |
| Dok5 | 1.986157553 | 0.97866935 |
| Amn | 4.8061 | 2.3678 |
| Tmem27 | 2.68219 | 1.32132 |
| Glt8d2 | 1.15468 | 0.568621 |
| Perp | 2.11505 | 1.04135 |
| Dhfr | 11.633559 | 5.7236123 |
| Pomc | 5.308520693 | 2.610814079 |
| Cdkn2aipnl | 55.37415 | 27.217995 |
| Clec2l | 23.6219 | 11.6095 |
| Ppih | 95.972938 | 47.159785 |
| Ppil3 | 48.409155 | 23.775353 |
| Mov10l1 | 3.6509676 | 1.793092425 |
| Gstp1 | 205.771 | 100.937 |
| Aven | 44.539562 | 21.84345 |
| Icam2 | 23.0093 | 11.2739 |
| Ccdc172 | 3.432541 | 1.681126 |
| Rpl5 | 1486.509822 | 728.0020017 |
| Vstm2l | 2.63677 | 1.290447465 |
| Klf1 | 1.67631 | 0.820284 |
| Snf8 | 113.649 | 55.6116 |
| B3galt6 | 10.7724 | 5.2703 |
| Sumo2 | 707.131 | 345.912 |
| Apitd1 | 21.73099 | 10.629188 |
| Rpsa | 3876.011498 | 1894.502215 |
| Apip | 50.5771 | 24.7068 |
| Fut8 | 16.79555904 | 8.2039117 |

| | | |
|---|---|---|
| Abracl | 76.8813 | 37.5294 |
| Ndufb10 | 199.547 | 97.4035 |
| Folr1 | 29.34111 | 14.32084778 |
| Lcn2 | 1.72335 | 0.839765 |
| Ephx2 | 12.47463 | 6.07864 |
| Rab26 | 2.416421 | 1.17722 |
| E030024N2 | 1.98651 | 0.967386 |
| Ptpmt1 | 41.8967 | 20.3979 |
| Cetn3 | 111.089 | 54.0792 |
| Psmb4 | 605.242 | 294.462 |
| Casp6 | 52.31285 | 25.444555 |
| Nat2 | 3.740024271 | 1.81843 |
| Rpl13a | 4831.14 | 2348.35 |
| Zfp946 | 5.075658 | 2.46666872 |
| Rtn1 | 2.466481 | 1.19859662 |
| Omd | 2.86454 | 1.39178 |
| Tmem120a | 59.5037 | 28.9038 |
| Psmg4 | 149.1198 | 72.4219 |
| Zfp948 | 4.07437 | 1.97821 |
| Cela3b | 1.91662 | 0.930548 |
| C1qtnf2 | 6.06079 | 2.94254 |
| Tefm | 19.67 | 9.54939 |
| Mrpl52 | 431.705 | 209.578 |
| Cox20 | 114.09106 | 55.3736 |
| Tmem80 | 24.7128 | 11.99243 |
| Cox16 | 59.632251 | 28.925133 |
| Dnase1l1 | 8.3520034 | 4.04878616 |
| Smagp | 16.5175 | 8.005527 |
| Dynll1 | 292.705649 | 141.847339 |
| H2-Ke6 | 51.7956 | 25.0995 |
| Blvrb | 64.8874 | 31.4354 |
| Bex1 | 80.4743 | 38.9862 |
| Fam120aos | 17.9761 | 8.70768 |
| Sod1 | 1183.93 | 573.285 |
| Gm10653 | 4.58654 | 2.22081 |
| Ppp1r14a | 36.2576 | 17.5523 |
| Dppa5a | 459.112 | 222.195 |
| Tmem37 | 23.8832 | 11.557 |
| B9d1 | 68.8266 | 33.2961 |
| Pard6a | 16.117144 | 7.79619037 |
| Atp6v1f | 266.884 | 129.095 |
| Grpel1 | 36.0826 | 17.4535 |
| Slc25a43 | 1.795483206 | 0.868426699 |
| Mrpl47 | 37.7632 | 18.2604 |
| Smdt1 | 263.51 | 127.305 |
| Atp5f1 | 365.383 | 176.5122283 |
| Aifm3 | 2.767266734 | 1.33663686 |

| | | |
|---|---|---|
| Mrpl17 | 24.303515 | 11.736908 |
| Nkain4 | 25.38063 | 12.252727 |
| Hoxb9 | 1.93017 | 0.93161 |
| Cdkn2a | 6.56208 | 3.16673 |
| Enho | 20.8374 | 10.0553 |
| Toporsos | 78.4173 | 37.8344 |
| Gm715 | 1.47488 | 0.711475 |
| Tmem258 | 534.73295 | 257.82265 |
| Trpc3 | 1.25564 | 0.604766 |
| Cox7a2l | 646.82109 | 311.48883 |
| Tspan32 | 3.6455632 | 1.755555 |
| Trp53rkb | 2.19748 | 1.05807 |
| Abcb6 | 24.986795 | 12.02802 |
| Acyp2 | 15.40612 | 7.41544 |
| Phlda2 | 88.0267 | 42.3679 |
| Znhit1 | 63.2533 | 30.4433 |
| Fam57a | 17.611349 | 8.47151847 |
| Nxnl2 | 2.55405 | 1.22851 |
| Mydgf | 116.507 | 55.9855 |
| Srgn | 4.74664622 | 2.28079714 |
| Hdhd3 | 16.5816 | 7.96491 |
| Phox2b | 1.147081 | 0.550931 |
| Ggh | 20.8902 | 10.0309 |
| Klk1 | 2.48685 | 1.19406 |
| Plek2 | 1.4915855 | 0.716079807 |
| Glrx | 61.1238 | 29.3385 |
| Tmem5 | 39.27949 | 18.85118 |
| Pdcd2 | 53.5629 | 25.703 |
| Gm8801 | 3.76456 | 1.806 |
| Snrnp25 | 64.6956 | 31.0145 |
| Ppapdc3 | 6.585347143 | 3.155341477 |
| Tmem147 | 189.23 | 90.5625 |
| Wfdc15a | 1.15688 | 0.553505 |
| Rpl18 | 6351.32 | 3038.48 |
| Rilpl2 | 35.4998 | 16.9761 |
| Krtcap3 | 15.06357 | 7.20335 |
| Rps21 | 7315.4786 | 3498.02 |
| Gm14325 | 7.971168 | 3.810969 |
| Zfp738 | 2.13034 | 1.01844 |
| Sla2 | 34.5815 | 16.5268 |
| Rhox9 | 146.68 | 70.095 |
| Postn | 27.01667 | 12.90356 |
| Smc1b | 6.39039625 | 3.051105 |
| Sf3b5 | 288.298 | 137.582 |
| Zfp54 | 1.061101681 | 0.505736942 |
| Stmn2 | 13.45973 | 6.41503 |
| Airn | 17.08588 | 8.142748 |

| | | |
|---|---|---|
| H2afz | 891.23418 | 424.583 |
| Pigw | 2.325296 | 1.10767 |
| Gm12504 | 1.02469 | 0.488096 |
| Rnf17 | 7.32536118 | 3.488406936 |
| Prdx1 | 627.848862 | 298.84794 |
| Rpl13 | 5709.51 | 2717.14 |
| Htra1 | 34.722 | 16.51844 |
| Zbtb8os | 57.026 | 27.1188 |
| Zfp874a | 3.16749 | 1.5052307 |
| Fam162a | 196.831 | 93.498 |
| Ntan1 | 74.0449 | 35.1294 |
| Fkbp1b | 9.25382 | 4.39022 |
| S100a1 | 45.3709 | 21.512 |
| Bvht | 9.74443 | 4.61954 |
| Mall | 1.055830561 | 0.500459704 |
| Eif3k | 727.7342375 | 344.9262838 |
| Qpct | 1.402635247 | 0.664804507 |
| Dtymk | 265.595 | 125.8809 |
| Anapc15 | 47.42197 | 22.46949 |
| Fhl2 | 8.41559 | 3.981694612 |
| Klk8 | 48.62279 | 22.97707 |
| Rpl21 | 805.62906 | 380.56014 |
| Pam16 | 132.091 | 62.3696 |
| Mcts2 | 38.4614 | 18.143 |
| Ormdl2 | 23.7906 | 11.2204 |
| Tstd3 | 9.33899 | 4.40452 |
| Fdx1l | 83.5848 | 39.3868 |
| Dnajc24 | 21.70609 | 10.22643 |
| Rpl14 | 2329.5722 | 1097.3877 |
| Trnp1 | 6.33543 | 2.98429 |
| Rps15a-ps6 | 118.566 | 55.8465 |
| Ninj1 | 59.2604 | 27.9123 |
| Npw | 7.14756 | 3.36654 |
| l7Rn6 | 78.258226 | 36.8561427 |
| Ccl24 | 1.33568 | 0.628932 |
| Ccdc152 | 2.696943 | 1.268581 |
| Lsm1 | 23.23783 | 10.92588 |
| Gm773 | 1.03726 | 0.487535 |
| Acot9 | 25.950637 | 12.1954 |
| Tmem60 | 37.60564 | 17.66206154 |
| Tma7 | 366.919 | 172.26893 |
| Gng10 | 104.809 | 49.1781 |
| H2-Oa | 1.46969 | 0.689541 |
| Minos1 | 57.690705 | 27.048524 |
| Gm13157 | 8.863031125 | 4.155102 |
| Psmb1 | 451.234 | 211.397 |
| Ptrhd1 | 5.31108 | 2.48612 |

| | | |
|---|---|---|
| Coprs | 85.1636 | 39.8613 |
| Gtf2a2 | 85.453329 | 39.987085 |
| Ap3s1 | 53.762 | 25.1561 |
| Rbm24 | 1.086027 | 0.507898892 |
| Sncg | 4.3911 | 2.05319 |
| Nsa2 | 128.486 | 60.0294 |
| Fam159a | 2.30915 | 1.07819 |
| Ccdc53 | 49.33707 | 23.02027 |
| Cox14 | 157.203 | 73.3132 |
| Hnmt | 1.0453 | 0.487434 |
| Calcoco2 | 6.82292 | 3.18031 |
| Efcab12 | 0.768462 | 0.357988318 |
| Gm3258 | 3.84067 | 1.7889 |
| Bola3 | 73.8696 | 34.3941 |
| Mnd1 | 25.48538138 | 11.86091 |
| Scarna13 | 7.2109 | 3.35452 |
| Cebpzos | 18.36672 | 8.54289 |
| Zfp930 | 6.93869516 | 3.22473296 |
| Mogat2 | 16.41228235 | 7.626131344 |
| Ndufs8 | 189.92331 | 88.22999 |
| Nup62cl | 3.7327 | 1.733981147 |
| Snca | 5.24178 | 2.434538231 |
| Rbm3 | 512.03486 | 237.58906 |
| Smim11 | 82.3662 | 38.2159 |
| Hist1h2ae | 13.7387 | 6.37413 |
| Fam92a | 57.1254 | 26.4971 |
| Kcnc4 | 1.064268 | 0.493519439 |
| Malsu1 | 60.5143 | 28.0602 |
| Sar1b | 57.0719 | 26.4618 |
| Atp5o | 958.484 | 444.237 |
| Rplp2 | 9548.61 | 4425.07 |
| Zfp52 | 1.47989 | 0.685787 |
| Fis1 | 416.3338 | 192.8818 |
| Sdhaf1 | 46.4201 | 21.5056 |
| Med30 | 57.462 | 26.5987 |
| Cox7a2 | 456.74 | 211.095 |
| Rpl7 | 4041.07 | 1866.43 |
| Ech1 | 163.464 | 75.4463 |
| Cbr4 | 17.8172 | 8.22178 |
| Gfer | 37.5508 | 17.3272 |
| Zfp943 | 5.15068 | 2.37528 |
| Mgst3 | 94.7201 | 43.6647 |
| Eef1b2 | 1137.419838 | 524.139375 |
| C1qtnf4 | 37.0841 | 17.0846 |
| Sec11c | 18.01341 | 8.29738 |
| Ndufa11 | 54.79787 | 25.231428 |
| H2afj | 30.4982 | 14.0426 |

| | | |
|---|---|---|
| Rpl22 | 545.994749 | 251.389012 |
| Spats1 | 1.804193 | 0.830395817 |
| Syce2 | 77.77637 | 35.79567 |
| Etfb | 291.7609 | 134.21169 |
| Gm38426 | 5.71749 | 2.62965 |
| Pfdn4 | 55.47528 | 25.50896 |
| Rnaseh2c | 236.322 | 108.627 |
| Snrnp27 | 82.5674 | 37.9489 |
| Atp5k | 1389.51 | 638.473 |
| Acta2 | 15.7055 | 7.21649 |
| Foxd1 | 11.1892 | 5.13925 |
| Lsm7 | 468.23955 | 214.988 |
| Mapk4 | 2.542639 | 1.16708 |
| Nedd8 | 399.237 | 182.978 |
| Car3 | 3.19712 | 1.46517 |
| N6amt2 | 78.1094 | 35.7832 |
| Rpl15 | 1569.56 | 718.845 |
| Glrx5 | 178.0219 | 81.52883 |
| Gin1 | 7.25696 | 3.32247 |
| Sct | 18.76166 | 8.588384 |
| Ndufaf2 | 43.0208 | 19.6744 |
| Tff2 | 7.14197 | 3.2658981 |
| Gm11744 | 48.04317 | 21.96729 |
| Gm10638 | 8.41711 | 3.84483 |
| Gpx1 | 542.415 | 247.749 |
| Emp3 | 168.7102 | 77.02236 |
| B230118H( | 18.069564 | 8.247851 |
| Gpbp1 | 58.71842842 | 26.77534466 |
| Anapc10 | 28.549958 | 13.01519 |
| Ppp1r14b | 471.069 | 214.726 |
| Polr2k | 107.1205 | 48.81343 |
| Gm4890 | 1.925169 | 0.876863 |
| Defb42 | 5.54238 | 2.52421 |
| Ppia | 5434.95 | 2474.45 |
| Hemk1 | 11.4792 | 5.225018 |
| Morc1 | 5.458503 | 2.484511 |
| Mrps33 | 124.0618 | 56.46219 |
| Ube2a | 48.82098 | 22.21795 |
| Pon3 | 5.0359461 | 2.2915755 |
| Shbg | 5.52776 | 2.514965 |
| Ctrl | 1.53106 | 0.696558 |
| Tm4sf1 | 6.54654 | 2.97694 |
| Gm128 | 0.818915 | 0.372176 |
| Tmem74b | 2.18709 | 0.993872 |
| Elof1 | 153.02 | 69.500178 |
| Scaper | 8.116234 | 3.685105 |
| Mrps24 | 107.51564 | 48.8085 |

| | | |
|---|---|---|
| Proca1 | 2.63608 | 1.196205 |
| Sys1 | 71.27526 | 32.33955 |
| Rpl30 | 230.7616 | 104.5838 |
| Ropn1l | 2.08394 | 0.94353 |
| Ecscr | 18.475 | 8.361186 |
| Gm6880 | 2.08601 | 0.944013 |
| Snrpd2 | 1049.913 | 475.087 |
| Spata24 | 33.5742 | 15.18656 |
| Atp5j | 602.89914 | 272.469434 |
| Tmem203 | 40.9012 | 18.4726 |
| Rps8 | 5478.7 | 2474.29 |
| Srp19 | 69.384 | 31.3262 |
| Nhej1 | 14.9444 | 6.74712 |
| AU041133 | 2.3594 | 1.0651 |
| Rpl38 | 5514.21 | 2485.526 |
| Gemin6 | 24.48893 | 11.03721 |
| Triqk | 6.00071 | 2.70405 |
| Cetn2 | 48.5898 | 21.895251 |
| Lipt2 | 29.0335 | 13.0702 |
| Mthfsl | 22.77703 | 10.25305 |
| Hmbs | 99.47305 | 44.76061 |
| Ube2t | 31.6147 | 14.21227 |
| Commd3 | 109.651 | 49.2917 |
| Isx | 2.343912449 | 1.0535346 |
| Derl3 | 10.71597319 | 4.81573 |
| Mfsd2a | 1.46941 | 0.659923 |
| Mrpl30 | 223.808 | 100.508 |
| Ndufa7 | 595.808163 | 267.560951 |
| Srp14 | 246.39 | 110.433 |
| Zfp944 | 4.74498 | 2.126643367 |
| Rpl31 | 1817.38992 | 814.31931 |
| Zfp931 | 4.1517248 | 1.859938 |
| C2cd4b | 1.27001 | 0.568632 |
| Rgn | 1.39401625 | 0.62391023 |
| Pcbd1 | 48.1902 | 21.5664 |
| Trem2 | 4.015475 | 1.796627704 |
| Cend1 | 0.800212504 | 0.357975978 |
| Emc2 | 44.22047727 | 19.77345978 |
| D330023K1 | 11.545277 | 5.161444 |
| Zfp868 | 12.84306 | 5.74038 |
| Ebpl | 24.19243 | 10.8094 |
| Dcun1d5 | 96.7408 | 43.21073 |
| Ppa2 | 39.07597 | 17.452678 |
| Ly6c1 | 2.58573899 | 1.154632309 |
| Rgs10 | 31.6434 | 14.1161 |
| Scand1 | 283.977 | 126.462 |
| Gm2694 | 8.19522 | 3.649035 |

| | | |
|---|---|---|
| Rpp38 | 19.4537 | 8.66145 |
| Atp5h | 687.203 | 305.811 |
| Thyn1 | 78.128 | 34.72207 |
| Slc25a31 | 7.34042 | 3.26194 |
| Ctla2b | 3.32355 | 1.47678 |
| Mtl5 | 1.509562 | 0.670561 |
| Aamdc | 35.89526 | 15.9392 |
| Atpif1 | 673.267554 | 298.918886 |
| Mrpl40 | 58.84 | 26.1044 |
| Psma7 | 416.727894 | 184.87313 |
| Ndufb7 | 359.723 | 159.534 |
| Hspb11 | 45.7498 | 20.26716 |
| Spata33 | 3.0409388 | 1.347003 |
| Trappc2 | 13.503927 | 5.97836324 |
| Cela1 | 8.222151688 | 3.638900513 |
| Lage3 | 119.084 | 52.7 |
| Gm12191 | 3893.07 | 1722.09 |
| Sfxn4 | 3.389672 | 1.499261925 |
| Tmem243 | 15.8764 | 7.01957 |
| Hspe1 | 833.793 | 368.048 |
| Tfec | 1.076878388 | 0.474621711 |
| Cbln1 | 3.36163 | 1.48113 |
| Serpinb1a | 4.62018 | 2.03219 |
| Gm10033 | 2.495323058 | 1.096991 |
| Hspb8 | 24.15534 | 10.618938 |
| Atp6v1g1 | 207.213 | 91.0762 |
| Mrps21 | 262.416 | 115.321 |
| Tex14 | 10.385519 | 4.563961 |
| Afp | 1.86217 | 0.818145 |
| Uqcrq | 478.723 | 210.197 |
| Abcc3 | 4.29969 | 1.886543 |
| Tmem261 | 136.238 | 59.7646 |
| Gabarap | 570.351 | 250.058 |
| Gnb2l1 | 5369.77 | 2353.39 |
| Efhd1 | 1.951219 | 0.854811 |
| Csrp2 | 106.324 | 46.5772 |
| Fut10 | 10.512695 | 4.601663285 |
| Colec11 | 2.204147 | 0.964789698 |
| Pdcd5 | 225.974 | 98.7732 |
| Ccdc23 | 150.1601 | 65.6128 |
| Pfdn5 | 640.133 | 279.359 |
| Pradc1 | 11.80548 | 5.142539 |
| Gstm5 | 166.499 | 72.5213 |
| Tesc | 2.48807 | 1.08264 |
| Mustn1 | 1.41086 | 0.61375 |
| Prr19 | 2.97736 | 1.29374 |
| Zfp677 | 2.0075015 | 0.8718854 |

| | | |
|---|---|---|
| Hscb | 51.8688 | 22.527 |
| Rpl37 | 1955.854844 | 848.967 |
| Rpl36 | 7239.52 | 3142.38 |
| Rpl35 | 5115.44 | 2220.27 |
| Emc4 | 76.7653 | 33.306 |
| Uqcrh | 1013.76 | 439.745 |
| Zfp273 | 2.176002 | 0.943799806 |
| Rps11 | 5845.5 | 2534.08 |
| Uba52 | 7350.78 | 3185.41 |
| Akr1c18 | 0.963539542 | 0.417481196 |
| Apela | 1.19186 | 0.515829 |
| Dmrta2 | 1.454542018 | 0.629362874 |
| Pigyl | 96.5369 | 41.767 |
| Iah1 | 97.0813 | 41.9695 |
| Atp8a2 | 3.934431 | 1.700594 |
| Slc25a5 | 497.749 | 215.091 |
| Hao2 | 1.176192884 | 0.508264694 |
| Ctla2a | 9.914856 | 4.284061 |
| Smim6 | 0.892393 | 0.38528 |
| Ccz1 | 68.3105 | 29.4897 |
| Tomm22 | 201.366 | 86.8453 |
| Mogat1 | 1.0661 | 0.459729 |
| Crip1 | 65.3094 | 28.1376 |
| Rpl10a | 4582.62 | 1974.18 |
| Nudt2 | 35.2343 | 15.1743 |
| Atf7ip2 | 2.88451 | 1.24222 |
| Ndufa12 | 337.101 | 145.052 |
| Lgals1 | 1006.7 | 433.114 |
| Rps15a-ps4 | 49.5614 | 21.3213 |
| Glo1 | 135.9398 | 58.44533 |
| Cst3 | 718.637 | 308.96 |
| Tph2 | 3.96 | 1.70155 |
| Mrpl14 | 197.951 | 85.0488 |
| Nenf | 301.787 | 129.652 |
| Siva1 | 418.18912 | 179.64918 |
| Serp2 | 9.47177487 | 4.0689354 |
| Rpl9 | 5982.37 | 2567.98 |
| Hist1h2bf | 1.69335 | 0.726249 |
| Pigx | 79.1722 | 33.95364 |
| Snx22 | 116.4718 | 49.94005692 |
| Pabpc1l | 1.29218 | 0.5538459 |
| Kcns3 | 4.394707856 | 1.88360239 |
| Vamp8 | 171.81897 | 73.62915 |
| Gm3604 | 2.371746858 | 1.016278 |
| Akr1c14 | 5.41454 | 2.31561 |
| Rbm44 | 4.68047 | 2.000826226 |
| Crp | 2.15404 | 0.920593 |

| | | |
|---|---|---|
| Rhox5 | 214.068 | 91.3769 |
| Dmrtc1a | 1.356493703 | 0.5786205 |
| Hsf2bp | 5.724438 | 2.4413 |
| Tuba1c | 66.9213 | 28.5357 |
| Hist1h4i | 48.5145 | 20.68229 |
| Dmrt2 | 1.85957 | 0.792754 |
| Ooep | 5.83705 | 2.48751 |
| Irak1bp1 | 7.23224 | 3.08186 |
| Rps16 | 7118.35 | 3032.91 |
| Rpl23a | 6615.52 | 2818.05 |
| Cdx2 | 3.21357 | 1.36836 |
| Cyb561d2 | 13.34837 | 5.68361 |
| Zfp97 | 0.955787 | 0.406933 |
| Adamts13 | 17.88169207 | 7.60913288 |
| Deb1 | 41.7919 | 17.7739 |
| Ttc30b | 4.59715 | 1.95454 |
| Dpm3 | 216.079 | 91.8069 |
| Cycs | 60.1156 | 25.5335 |
| Cilp2 | 2.17198 | 0.922324 |
| Rpl7a | 5078.07 | 2156.28 |
| Ms4a7 | 2.299574 | 0.9761514 |
| Ptcd2 | 55.2413 | 23.4425 |
| Dmc1 | 2.076373 | 0.8810392 |
| Eif1ax | 215.887 | 91.5334 |
| Fibin | 1.23373 | 0.522726 |
| Fuom | 11.031725 | 4.670661 |
| Kirrel3 | 2.59905057 | 1.10026842 |
| Gtsf1l | 4.16769 | 1.7643 |
| Pgls | 354.33493 | 149.99329 |
| Nsg2 | 2.6694825 | 1.130015612 |
| Bad | 105.6009 | 44.66481 |
| Gm5148 | 84.3451 | 35.6742 |
| Ndufs5 | 182.477 | 77.15623265 |
| Hist1h2bh | 3.49083 | 1.47565 |
| Atp6v0e | 145.453 | 61.4774 |
| Tuba3a | 53.0306 | 22.4127 |
| Cryge | 8.282791107 | 3.49978 |
| Fabp5 | 133.4008 | 56.311318 |
| Plekhb1 | 4.539895568 | 1.914566515 |
| M1ap | 5.516481 | 2.32360146 |
| Fau | 5288.841 | 2227.55672 |
| Gm19461 | 1.660892 | 0.699308 |
| Magoh | 260.466 | 109.653 |
| Pdf | 24.4099 | 10.2595 |
| Atp5e | 1738.47 | 730.45 |
| Rps9 | 9076.42 | 3812.23 |
| Rab29 | 8.24766 | 3.45824 |

| | | |
|---|---|---|
| Nxt1 | 97.788 | 40.99283 |
| Uqcr10 | 589.303 | 246.823 |
| Bbip1 | 14.33234 | 6.00144 |
| Mageb16 | 11.40887 | 4.77618 |
| Rpl17 | 4158.63 | 1739.13 |
| Zfp119a | 2.64433 | 1.10513 |
| Cdkn3 | 18.86233 | 7.87726 |
| Dazl | 67.47065 | 28.15296 |
| Rps20 | 10264.3 | 4282.03 |
| Pstk | 39.85283 | 16.622852 |
| Lsm3 | 221.475 | 92.3636 |
| Pdlim2 | 45.68219 | 19.04909 |
| Tmsb15l | 12.6872 | 5.28979 |
| Bambi-ps1 | 0.983554 | 0.409868 |
| Nhp2 | 203.179 | 84.6627 |
| Gm4944 | 3.14111624 | 1.308513 |
| Steap1 | 2.15138 | 0.896027 |
| Cib1 | 71.93999 | 29.953393 |
| Rps7 | 3821.02 | 1590.89 |
| Gm13051 | 3.61063 | 1.50223 |
| Chrac1 | 74.0802 | 30.8181 |
| Gm9112 | 9.35944 | 3.89055 |
| Cox7c | 1188.87 | 493.707 |
| Mrpl54 | 226.213 | 93.9303 |
| Gm5512 | 1.243980338 | 0.515971 |
| Gal3st1 | 3.614549791 | 1.498817 |
| Rpl19 | 5516.8409 | 2285.989 |
| Prg4 | 1.08595 | 0.4494561 |
| Tbca | 432.629 | 178.785 |
| Hist1h3c | 30.0848 | 12.4321 |
| Rpl27 | 3841.58 | 1586.62 |
| Ndufa6 | 635.961 | 262.237 |
| Jmjd7 | 8.62843 | 3.55782 |
| C030013C2 | 1.26593 | 0.521988 |
| Gm15441 | 1.39445 | 0.574951 |
| Slirp | 291.628 | 120.23 |
| Sec61b | 586.193 | 241.569 |
| Mrps22 | 46.899 | 19.2893 |
| Bcap29 | 21.39602 | 8.795934 |
| Ndufb6 | 352.953 | 145.0865 |
| Rps10 | 7571.66 | 3111.92 |
| Spaca1 | 2.85134 | 1.171657 |
| Fkbp3 | 262.883 | 107.922 |
| Defb25 | 1.80011 | 0.737936 |
| Snord15a | 88.0422 | 36.0504 |
| Polr2f | 403.73 | 165.244 |
| Ndufa13 | 505.788 | 207.0081 |

| | | |
|---|---|---|
| Cmbl | 13.84587 | 5.66433 |
| Hist1h2bg | 7.32157 | 2.99422 |
| Arhgdig | 36.8008 | 15.0398 |
| Klhl32 | 4.201043544 | 1.716539674 |
| Foxr1 | 22.12645 | 9.039025 |
| Cenpq | 17.5125 | 7.15231 |
| Gm5617 | 132.582 | 54.1013 |
| Mien1 | 64.1639 | 26.1753 |
| Zfp708 | 1.49982229 | 0.611608 |
| Cyp17a1 | 1.37117 | 0.558982 |
| Cox7b | 213.782007 | 87.0748 |
| Romo1 | 399.6187 | 162.64583 |
| Gm1673 | 293.882 | 119.5861 |
| Agpat9 | 2.548310148 | 1.036741 |
| Arxes1 | 17.7463 | 7.2163 |
| Ndufab1 | 306.0587 | 124.3156 |
| Rpl35a | 5176.4059 | 2102.3778 |
| Adh1 | 14.9979 | 6.08459 |
| Sdhaf4 | 87.616 | 35.5299 |
| Anxa13 | 1.24915 | 0.506431 |
| Lurap1l | 3.07955 | 1.2484 |
| Dnajb3 | 16.286 | 6.6014 |
| Rhox13 | 1.96444 | 0.795821 |
| Myeov2 | 456.13058 | 184.740196 |
| Nme2 | 1454.104 | 588.489 |
| BC028528 | 22.50702 | 9.10639 |
| Hsbp1l1 | 2.80597 | 1.13411 |
| Pla2g5 | 6.771368 | 2.735496424 |
| Cox4i1 | 1740.4986 | 702.693556 |
| Dbil5 | 4.77255 | 1.92301 |
| Krt79 | 1.04368 | 0.420512 |
| Etohi1 | 11.740843 | 4.72955 |
| Smim8 | 27.87817 | 11.20402 |
| Fbxo6 | 43.083111 | 17.293122 |
| Pgp | 113.0497 | 45.3765 |
| Gm20594 | 9.94004 | 3.98859 |
| Cdx1 | 1.952093 | 0.783214086 |
| Pfdn1 | 148.15074 | 59.37334 |
| Spa17 | 9.529881 | 3.818056 |
| Fam228b | 1.4073118 | 0.563586 |
| Ngp | 2.39938 | 0.960634 |
| Nanos1 | 1.19747 | 0.479326 |
| Tcfl5 | 4.90115 | 1.96178 |
| Tmem121 | 17.3033 | 6.92347 |
| Rps5 | 8854.77 | 3540.48 |
| Rps23 | 6288.35 | 2511.33 |
| Emx1 | 1.47134 | 0.58753 |

| | | |
|---|---|---|
| Gimap9 | 5.4772 | 2.186178 |
| Ndufs6 | 352.17671 | 140.46619 |
| Aqp1 | 5.73549 | 2.28444 |
| Hist1h2bl | 6.22155 | 2.47796 |
| Rps15a | 558.365773 | 222.2899141 |
| Mpc1 | 59.1691 | 23.53932 |
| Ppp1r36 | 2.38134 | 0.947091 |
| Nr2e3 | 0.994989 | 0.395656 |
| Hint1 | 1185.81 | 470.583 |
| Zfp729b | 4.640509881 | 1.841460709 |
| Zfp958 | 7.79559 | 3.09326 |
| Ccdc28a | 13.0056 | 5.15963 |
| Zswim7 | 51.676 | 20.4641 |
| D17Ertd64{ | 1.12103 | 0.443524 |
| Mrpl23 | 314.54 | 124.388 |
| Gsto2 | 2.167089777 | 0.855732429 |
| Tmsb4x | 2521.61 | 994.938 |
| Cited1 | 11.535299 | 4.54860954 |
| Tmem134 | 125.9362 | 49.61794 |
| Zfp936 | 1.362950642 | 0.536473793 |
| Rhox6 | 124.887 | 49.1458 |
| Rpl34 | 6571.7206 | 2585.6049 |
| Mettl5 | 17.94185 | 7.05748 |
| Chchd2 | 817.0019 | 321.3188 |
| Pet100 | 25.67862 | 10.09772 |
| Ufsp1 | 6.03492 | 2.37220483 |
| Fbxo47 | 1.290549 | 0.506990891 |
| Dnajc19 | 104.422177 | 40.98782 |
| Tctex1d2 | 33.08776 | 12.98733 |
| Jazf1 | 1.345703862 | 0.52798956 |
| Hddc2 | 77.66 | 30.428991 |
| Commd1 | 187.53 | 73.4658 |
| Lamtor4 | 93.0463 | 36.4473 |
| Dynlt1f | 60.5576 | 23.68378483 |
| Ndufb2 | 422.772 | 164.8669 |
| Cst8 | 39.73454 | 15.4845 |
| Batf3 | 9.404479 | 3.662267 |
| Cntf | 1.12132 | 0.436557 |
| Rpl32 | 9619.51 | 3743.7 |
| Dppa3 | 205.168 | 79.8255 |
| BC048679 | 26.99931521 | 10.49960096 |
| Rps18 | 9507.1 | 3696.31 |
| Gml2 | 1.271884 | 0.49421 |
| Dctpp1 | 537.331 | 208.68 |
| Hist1h2bj | 11.04805 | 4.28931 |
| Sf3b6 | 122.94289 | 47.72499 |
| Znhit3 | 37.1444 | 14.4159 |

| | | |
|---|---|---|
| Mgmt | 47.3846 | 18.3854 |
| Gng11 | 64.7673 | 25.1244 |
| Rpl11 | 5814.44 | 2255.5 |
| Uqcr11 | 719.322 | 278.814 |
| Rpa3 | 135.992 | 52.6981 |
| Rp9 | 76.5123 | 29.6327 |
| Rhox1 | 10.7506 | 4.16143 |
| Ccdc107 | 55.4032 | 21.4268 |
| Kcnk12 | 1.34869 | 0.52153 |
| Uchl4 | 1.0828 | 0.41829 |
| Cenpv | 50.627608 | 19.551213 |
| Mrpl33 | 283.965 | 109.557 |
| Xrcc6 | 86.5931 | 33.40264 |
| Lsm5 | 180.738 | 69.6985 |
| Pdzk1ip1 | 13.433264 | 5.176451 |
| Bloc1s1 | 198.942 | 76.5992 |
| Aard | 15.4849 | 5.94385 |
| Ttpa | 1.29894 | 0.497357 |
| Rps27a | 5851.527 | 2240.322 |
| Zfp951 | 2.595865 | 0.99264142 |
| Rfc4 | 94.1633 | 35.991 |
| Rps15 | 7895.19 | 3017.62 |
| Nol7 | 57.49837 | 21.96069 |
| Rps24 | 4001.2055 | 1528.0606 |
| Cox6c | 1573.01 | 600.643 |
| Bola1 | 54.286 | 20.72714 |
| Ndufa5 | 179.017 | 68.2714 |
| Tmem14a | 4.202151 | 1.602272 |
| Cst9 | 1.74129 | 0.663319 |
| Hint2 | 116.208 | 44.2381 |
| Pcolce2 | 2.08691 | 0.794331 |
| Ulbp1 | 1.506531099 | 0.573403883 |
| Akr1cl | 140.369 | 53.4171 |
| Tmem100 | 1.120969373 | 0.426452552 |
| Cenpk | 13.195225 | 5.01523 |
| Arxes2 | 46.4647 | 17.6558 |
| Prdx4 | 327.602 | 124.32 |
| Rps27l | 805.044 | 305.223 |
| Ndufb9 | 514.567 | 194.997 |
| Batf | 1.33292 | 0.504542 |
| Rpl12 | 6532.1 | 2472.55 |
| Etaa1os | 11.0566 | 4.18409 |
| Rpl26 | 8589.446 | 3248.767 |
| Gm14326 | 4.231192 | 1.600262 |
| LOC10050[illegible] | 10.5052 | 3.97152 |
| Fabp4 | 25.051 | 9.453 |
| Rpl39l | 73.495 | 27.6818 |

| | | |
|---|---|---|
| BC002163 | 58.5424 | 22.0477 |
| Rhox7a | 13.329367 | 5.017369 |
| Serpinb6b | 66.2523 | 24.9367 |
| Rpl14-ps1 | 336.319 | 126.5266 |
| Hist1h2bm | 10.8314 | 4.07488 |
| S100a16 | 111.203 | 41.82 |
| Mt3 | 4.21671 | 1.58201 |
| Pemt | 16.64621 | 6.242506712 |
| Ndufc2 | 413.846 | 155.151 |
| Gpx4 | 796.780264 | 298.675608 |
| Bik | 9.81019 | 3.664898139 |
| Mrpl53 | 128.735 | 48.0823 |
| Slc51a | 0.957696 | 0.357412 |
| AU021092 | 2.058874955 | 0.767653 |
| Saxo2 | 2.8331684 | 1.056300133 |
| Iqcd | 1.6888858 | 0.629409 |
| Angpt2 | 1.07524 | 0.400697 |
| Fam92b | 1.486574 | 0.5532708 |
| Slc16a4 | 5.431127 | 2.02006861 |
| Aqp11 | 4.911138 | 1.825323 |
| Ssr4 | 282.0113 | 104.6668 |
| Nfe2 | 3.049626266 | 1.13183861 |
| Atp6ap1l | 1.78911 | 0.663539 |
| Foxp2 | 3.6240553 | 1.343297 |
| Tagln | 14.9675 | 5.54781 |
| Rsph3a | 6.21065 | 2.30095 |
| Gm4737 | 46.9046 | 17.3414 |
| Hist1h3i | 5.55712 | 2.05453 |
| Mlf1 | 7.46058 | 2.75533 |
| Serf1 | 234.575 | 86.6074 |
| Hist1h2bp | 1.291128 | 0.476660228 |
| Sat2 | 37.04304 | 13.62406 |
| Ndufb11 | 698.887 | 256.4615 |
| Cox7a1 | 64.9409 | 23.8169 |
| Mpc2 | 90.2634 | 33.0617 |
| Gm960 | 0.969761 | 0.3548394 |
| Cib2 | 105.055971 | 38.43523 |
| Oxld1 | 20.75871 | 7.591816 |
| Med31 | 43.1591 | 15.76915 |
| Khdc3 | 1.076601416 | 0.393078848 |
| Rplp1 | 14200.4 | 5183.45 |
| Rpl22l1 | 1669.708 | 609.339 |
| Snrpe | 1089.95 | 397.13 |
| Prn | 1.01514 | 0.369867 |
| Rpl36a | 4236.92 | 1543.26 |
| Fhl4 | 1.67353 | 0.608844 |
| Rpl41 | 13745.7 | 5000.54 |

| | | |
|---|---|---|
| Prkg1 | 1.622634 | 0.590089 |
| Efcab10 | 2.98338 | 1.08468 |
| Myl6 | 1026.742 | 371.7 |
| Zfp85 | 3.30723 | 1.19679 |
| Nox1 | 4.41582 | 1.59503 |
| Cav1 | 60.03166 | 21.662709 |
| Tmem116 | 1.864127911 | 0.67112343 |
| Fabp7 | 8.59449 | 3.0934 |
| Rps17 | 8758.59 | 3150.83 |
| Tmem174 | 1.51867 | 0.546076 |
| Psma8 | 3.76213 | 1.35144 |
| Rps2 | 9746.94 | 3499.85 |
| Gm12657 | 71.7117 | 25.6952 |
| Tuba3b | 8.93654 | 3.20145 |
| Rpl27a | 2745.58124 | 981.82909 |
| Eno1b | 82.1816 | 29.3526 |
| Zfp119b | 3.062650959 | 1.093611 |
| Stmn3 | 5.89649 | 2.10464 |
| S100a13 | 44.9532858 | 16.0103836 |
| Serpinb6a | 274.15452 | 97.590075 |
| Ndufa2 | 449.665 | 159.989 |
| Ogn | 2.54013 | 0.903313 |
| Usmg5 | 831.536 | 295.685 |
| Gm11837 | 3.941757 | 1.399083 |
| Fam229b | 10.54777 | 3.740678 |
| Tomm5 | 220.94005 | 78.27787 |
| Tex30 | 45.67717 | 16.17639 |
| Rpph1 | 186.251 | 65.9519 |
| Umodl1 | 3.415923 | 1.206726 |
| Ddx43 | 1.00189 | 0.353743 |
| Snora43 | 8.72744 | 3.07869 |
| Rps27 | 12290.0746 | 4329.521161 |
| Ndufc1 | 386.781 | 136.136 |
| Uchl3 | 121.717 | 42.8335 |
| Cetn4 | 2.923378 | 1.027693 |
| Tsacc | 14.3336 | 5.036013 |
| Ppp1r14d | 1.979195072 | 0.695116959 |
| Snapc5 | 19.1793 | 6.73235 |
| C1qtnf7 | 46.513173 | 16.3178 |
| Rpl37rt | 248.63161 | 87.1834 |
| Gm10046 | 3.16671 | 1.11041 |
| Cbr3 | 17.6363 | 6.16909 |
| Smpx | 18.570815 | 6.491235379 |
| Cmc4 | 14.212 | 4.9606 |
| Cenpp | 20.8232 | 7.26188 |
| Taf10 | 290.915 | 101.44 |
| Il13ra2 | 1.051807804 | 0.366452557 |

| | | |
|---|---|---|
| Mrps16 | 315.289 | 109.644 |
| Fcrls | 5.84612 | 2.03108 |
| Rpl39 | 11365.12 | 3944.583 |
| Cpsf2 | 243.69083 | 84.57629 |
| Atp5j2 | 1207.82 | 418.48 |
| Med21 | 105.681 | 36.5469 |
| Tmem141 | 42.0277577 | 14.53117429 |
| Fn3k | 1.15732 | 0.40007 |
| D630033O: | 2.084371065 | 0.719808135 |
| Ost4 | 438.5 | 151.3457 |
| Gdf15 | 1.48054 | 0.510889667 |
| Gm4013 | 8.49372 | 2.92213 |
| Frmd7 | 3.67467 | 1.26413 |
| Actg2 | 1.6332 | 0.561751 |
| Uqcc2 | 544.392 | 187.036 |
| Kiss1 | 4.46938 | 1.52654 |
| Tmem40 | 1.948818319 | 0.664516226 |
| Coa6 | 62.4435 | 21.283 |
| Mmd2 | 1.18851 | 0.404456 |
| S100a10 | 504.664 | 171.3 |
| Pop5 | 99.0578 | 33.5574 |
| Bola2 | 485.706 | 164.485 |
| Rps14 | 12867.2 | 4351.28 |
| Rps12 | 10495.6 | 3547.06 |
| Mylpf | 9.115215 | 3.079751 |
| Syce1 | 7.96893 | 2.68802 |
| Kcnip4 | 1.072793304 | 0.361596838 |
| Zfp51 | 5.875609178 | 1.980393186 |
| Lrif1 | 9.74185 | 3.280252 |
| Uxt | 49.76271 | 16.739899 |
| Cyp21a1 | 1.23076 | 0.41374 |
| Npb | 2.22205 | 0.746475 |
| Hist1h1d | 9.37065 | 3.13565 |
| Tmem126a | 74.3124 | 24.8462 |
| Adamts5 | 1.87099 | 0.625314 |
| Npm3-ps1 | 7.49781 | 2.49122 |
| Sec61g | 758.7977 | 251.76435 |
| Hist1h2ak | 4.65351 | 1.54192 |
| Scarletltr | 2.59472 | 0.858369 |
| Mrpl20 | 221.756 | 73.2653 |
| C8g | 6.631331 | 2.18718 |
| Atp5l | 917.175 | 301.558 |
| Ndufb4 | 533.668 | 175.432 |
| Trap1a | 16.3355 | 5.36042 |
| Cyb5a | 276.152 | 90.5642 |
| Gm6682 | 8.45745 | 2.77229 |
| Chrna7 | 3.8749 | 1.268530129 |

| | | |
|---|---|---|
| Mrps18c | 188.611 | 61.7439 |
| Vnn1 | 3.43693 | 1.12322 |
| Pigp | 132.563463 | 43.2909858 |
| Hint3 | 37.58138 | 12.265553 |
| Hist1h2bb | 9.09473 | 2.9672 |
| Kbtbd6 | 1.66114 | 0.540288 |
| Gp1bb | 3.869041 | 1.258119 |
| Thnsl1 | 4.99464 | 1.62388 |
| Mrap | 2.24542 | 0.728834 |
| Ndufb8 | 671.833 | 218.059 |
| Lrrc3b | 3.80708 | 1.23351 |
| Blnk | 5.643022 | 1.826684 |
| Cyp11b1 | 6.528059 | 2.1118014 |
| Meox1 | 2.1212 | 0.685662 |
| Tpt1 | 11290.4 | 3649.38 |
| Mc2r | 3.354741685 | 1.082039072 |
| Gcnt1 | 2.21126611 | 0.713181761 |
| Fam46b | 2.5103 | 0.809094 |
| Pla2g2f | 0.972811 | 0.3133 |
| Tmem205 | 75.60012733 | 24.229273 |
| Selenbp2 | 1.59295 | 0.509572 |
| Taf7l | 4.15757 | 1.329782924 |
| Mfap5 | 2.060991 | 0.658071 |
| Crygs | 6.689019318 | 2.135588837 |
| Nanos3 | 19.21157 | 6.13357 |
| Tmem256 | 453.859 | 144.808 |
| Hist1h2bn | 6.33309 | 2.01931 |
| Dnajc15 | 177.831 | 56.6958 |
| Hist1h2ag | 13.3821 | 4.26018 |
| Rpl31-ps12 | 788.951 | 250.892 |
| Tnni1 | 63.53789846 | 20.11440246 |
| Hist1h2bk | 10.0271 | 3.17267 |
| Gm13212 | 3.15992 | 0.997794 |
| Gm12338 | 116.834 | 36.8367 |
| Serpina3a | 69.07979 | 21.72773 |
| Fmr1nb | 15.92877 | 5.008536 |
| Hist1h3h | 12.5737 | 3.95117 |
| Rec114 | 14.762 | 4.63854 |
| Hist1h4f | 4.9592 | 1.55544 |
| Nudt8 | 61.7335 | 19.3179 |
| Hspb2 | 1.70188 | 0.532355 |
| Myl4 | 3.61747 | 1.12596 |
| Gm6251 | 2.43638 | 0.756337 |
| Nrtn | 53.0042 | 16.4089 |
| Gm4285 | 2.64606 | 0.817775 |
| Gm4984 | 1.06359 | 0.328379 |
| Chchd1 | 269.288 | 82.999 |

| | | |
|---|---|---|
| C130026L2 | 1.274149979 | 0.392161723 |
| Hotairm1 | 15.58269 | 4.786978 |
| Ndufa4 | 932.635 | 286.024 |
| Timm8b | 353.042 | 107.846 |
| Klhl10 | 1.5223 | 0.464956 |
| Slc52a3 | 3.097426 | 0.940965631 |
| Tex19.2 | 3.3759 | 1.02403 |
| Cryba4 | 3.07679 | 0.931765 |
| Gng5 | 1296.1 | 392.218 |
| Hist1h4h | 10.2325 | 3.09314 |
| Naa38 | 219.44 | 66.1844 |
| Gm15421 | 77.2149 | 23.045 |
| Sec1 | 1.73307 | 0.516183 |
| Slc4a8 | 2.639047 | 0.785700711 |
| Ccdc122 | 1.24507 | 0.369496 |
| Apoo | 44.468689 | 13.142537 |
| Rprml | 2.68664 | 0.793282 |
| Immp2l | 12.2662 | 3.61257 |
| C230072F1 | 1.130151651 | 0.331475569 |
| Aqp2 | 2.04587 | 0.599254 |
| Vwf | 3.73885 | 1.091158 |
| Boll | 1.699379002 | 0.49287901 |
| Lhcgr | 5.14639 | 1.492494 |
| Pf4 | 46.1355 | 13.3225 |
| Hormad1 | 8.875574 | 2.560221173 |
| Cyp2r1 | 1.339739172 | 0.384070151 |
| Sh3bgr | 4.4491 | 1.27127 |
| Lrguk | 3.907654 | 1.115452437 |
| Car2 | 15.5383 | 4.430572 |
| Apoa2 | 2.617945 | 0.746331 |
| Mir692-2 | 2.75948 | 0.786256 |
| Ppp1r1a | 6.87648 | 1.95675 |
| Zbtb11os1 | 8.76705 | 2.49262 |
| Zfp947 | 3.496355 | 0.9915955 |
| Itgb3bp | 3.70144 | 1.046536 |
| Tnni2 | 1.38025 | 0.387906 |
| Hist1h4d | 12.7304 | 3.57315 |
| Endog | 22.3646 | 6.26493 |
| Hist1h3f | 2.18689 | 0.611927 |
| Hist1h4k | 7.204247 | 2.010858 |
| Fam132a | 93.0954 | 25.9103 |
| Mup6 | 1.299883718 | 0.361365 |
| Hist1h1a | 10.1305 | 2.81413 |
| Sdcbp2 | 1.350448054 | 0.374317742 |
| Hist2h2bb | 22.5262 | 6.21228 |
| Hacd1 | 71.42837 | 19.677121 |
| Zfp345 | 1.23366 | 0.339247 |

| | | |
|---|---|---|
| Eif4ebp3 | 9.93104 | 2.72063 |
| Wfdc10 | 12.0335 | 3.29198 |
| Xlr3b | 4.2185 | 1.15212 |
| Mir8109 | 2.09034 | 0.569485 |
| Xlr3c | 4.9435 | 1.34611 |
| Gypa | 2.43306 | 0.661395 |
| Thoc2 | 9.86329288 | 2.67922 |
| Tmsb15b1 | 23.9879 | 6.48294 |
| Gm13154 | 12.01384 | 3.246147 |
| Ncmap | 1.952487416 | 0.526717697 |
| Hist1h4a | 4.70437 | 1.26701 |
| Rmrp | 741.841 | 199.693 |
| Sycp3 | 8.48527 | 2.26375 |
| Hist1h1e | 10.2841 | 2.73348 |
| Ankrd29 | 1.35514 | 0.35983 |
| Nupr1l | 4.83751 | 1.28278 |
| Rps19-ps3 | 2.54478 | 0.673334 |
| Gm17660 | 4.80138 | 1.26716 |
| Cxcl13 | 4.904316914 | 1.278371 |
| Hist1h1b | 8.26929 | 2.13987 |
| Ascl2 | 2.57967 | 0.664027 |
| SlcUbl4a | 44.397379 | 11.4243 |
| Cdkn2b | 1.21139 | 0.308711 |
| Inca1 | 2.615756 | 0.665552127 |
| Stra8 | 8.75557 | 2.22611 |
| Myh14 | 1.27447699 | 0.32140573 |
| Urah | 1.49557 | 0.377062 |
| Xlr4a | 2.11714 | 0.531058 |
| Cela2a | 1.40091 | 0.350271 |
| Tmem219 | 25.7835 | 6.43566 |
| Guca1a | 1.94186 | 0.484367 |
| Mir99ahg | 9.574514 | 2.3781935 |
| Hist1h2be | 9.69178146 | 2.406800393 |
| Ccl19 | 4.82931 | 1.19879 |
| C2cd4a | 1.02837 | 0.253682 |
| Hist1h2ab | 5.12469 | 1.2635 |
| Hsd3b1 | 6.67328 | 1.641281 |
| Acad12 | 3.093657 | 0.758192424 |
| Stard6 | 1.171222725 | 0.284714 |
| Hist3h2ba | 57.1629 | 13.8802 |
| Asb9 | 1.93911 | 0.470535 |
| Hist1h2ah | 1.79687 | 0.432583 |
| Retn | 1.38607 | 0.333179 |
| Hist4h4 | 7.2772 | 1.74344 |
| Gm13139 | 23.88498194 | 5.712896111 |
| Slc22a18 | 14.54941 | 3.470938501 |
| Gm10416 | 14.71819549 | 3.504567 |

| | | |
|---|---|---|
| Cyct | 56.9278 | 13.5406 |
| Cdh12 | 1.38736 | 0.329483 |
| Paqr9 | 1.28818 | 0.305586 |
| Hist1h3g | 3.80691 | 0.901865 |
| Ctxn3 | 7.35024997 | 1.726170829 |
| S100a4 | 4.7784 | 1.1152 |
| Dync1i1 | 1.62864114 | 0.37848723 |
| Apoc1 | 85.381 | 19.6784 |
| Spink1 | 14.5918 | 3.36134 |
| Mir5133 | 1.18787 | 0.272654 |
| Ugt8a | 1.60591 | 0.368487 |
| Rhd | 3.37807 | 0.773487 |
| Lmo1 | 37.012707 | 8.463582 |
| Slc6a17 | 1.535930948 | 0.3506418 |
| Actn2 | 19.50664 | 4.44855 |
| Resp18 | 1.1009 | 0.250412 |
| Hsd3b4 | 1.18574 | 0.268758 |
| Srd5a2 | 2.162893 | 0.490171687 |
| Scarna6 | 5.9171 | 1.33931 |
| Gm6402 | 9.26974 | 2.09008 |
| Pglyrp1 | 9.67893 | 2.18194 |
| Ndufaf6 | 28.6377 | 6.428943 |
| Hist1h3d | 5.10856 | 1.14627 |
| Tg | 0.96717226 | 0.216671 |
| Aurkc | 8.019610565 | 1.785231868 |
| Foxd2os | 4.782605481 | 1.063107275 |
| Gm5635 | 2.04854 | 0.451866 |
| Hist1h3a | 11.1795 | 2.46271 |
| Akr1b7 | 2.9675 | 0.648738 |
| Hba-x | 2177.38 | 472.793 |
| E030030I0( | 3.39169 | 0.731879 |
| Gm14305 | 1.14425 | 0.246044 |
| Acbd7 | 2.38282 | 0.511004384 |
| Fggy | 13.017906 | 2.788053422 |
| Cst12 | 22.2558 | 4.75486 |
| Raet1a | 3.076446 | 0.65707 |
| Ly6d | 4.22283 | 0.900745 |
| Gm41 | 2.239147 | 0.474804302 |
| Gpha2 | 3.93162 | 0.821827 |
| Tfdp1 | 40.63346236 | 8.4613 |
| Pp2d1 | 2.41517 | 0.50015 |
| Casc1 | 7.989426 | 1.6528969 |
| Hist1h3e | 18.7432 | 3.86986 |
| Stfa2 | 3.77061 | 0.777683 |
| Tex11 | 1.854375 | 0.379608 |
| Tex40 | 2.04597 | 0.412581 |
| Ppbp | 6.05706 | 1.21438 |

| | | |
|---|---|---|
| Gdpd3 | 3.9127 | 0.778939 |
| Agt | 1.075 | 0.211708 |
| Tex101 | 12.6436 | 2.48834 |
| Prap1 | 2.49843 | 0.489055 |
| Trim10 | 1.593382655 | 0.29990843 |
| LOC100049 | 1.76431 | 0.328307 |
| Mael | 49.8445 | 9.18367 |
| Crygc | 2.626078 | 0.48375215 |
| Dydc2 | 1.15199 | 0.2105 |
| Cyp4b1-ps2 | 2.73874 | 0.495816 |
| Kel | 1.51532 | 0.272775 |
| Tubb1 | 1.443701 | 0.2592515 |
| Islr | 65.92492 | 11.8208 |
| Gm5595 | 119.08132 | 20.37088 |
| Cox7b2 | 3.85385 | 0.654338 |
| Raet1d | 7.725 | 1.30821 |
| Snord17 | 85.5261 | 14.4187 |
| Lars2 | 5282.63144 | 880.27656 |
| Aqp12 | 1.101702 | 0.182861913 |
| Med9os | 3.81289 | 0.628823 |
| Lgals2 | 1.31454 | 0.216388 |
| Upp2 | 11.79995068 | 1.935446966 |
| Thrsp | 1.91053 | 0.310802 |
| Gtsf1 | 3.59068 | 0.581158 |
| Lect2 | 1.25949 | 0.20362 |
| Gm14295 | 52.5291 | 8.30401 |
| Hbb-b2 | 628.728 | 98.4233 |
| Tmsb15b2 | 24.7791 | 3.8657 |
| Cldn13 | 1.08747 | 0.169523 |
| Xlr | 4.165522607 | 0.644225 |
| Hbb-y | 11000.4 | 1697.51 |
| Fbp1 | 2.349093 | 0.360508515 |
| C920006O1 | 1.82017 | 0.278795 |
| Usp26 | 2.257233 | 0.344961651 |
| Spink4 | 9.93621 | 1.51732 |
| Mir703 | 4152.14 | 630.266 |
| Ocel1 | 8.0604 | 1.21721 |
| Alas2 | 17.53201 | 2.643811882 |
| Apoc4 | 3.32988 | 0.493908 |
| Serpina1b | 1.760578613 | 0.257444971 |
| Coch | 3.51359 | 0.512478372 |
| Gjd3 | 1.07044 | 0.155888 |
| Usp9x | 22.74073 | 3.26425 |
| Gsc2 | 1.26925 | 0.175844 |
| Gm10012 | 2.09838 | 0.287038 |
| Tmed5 | 3.12413 | 0.416622 |
| Crxos | 6.3804791 | 0.850116686 |

| | | |
|---|---|---|
| Hist1h2ai | 7.34684 | 0.969563 |
| Jarid2 | 9708.73347 | 1264.96278 |
| Slc4a1 | 6.03676 | 0.780232 |
| BC100530 | 17.1591 | 2.16869 |
| Treml1 | 2.397054519 | 0.294864712 |
| BC049762 | 1.584441759 | 0.192205373 |
| Gm1564 | 2.41728 | 0.292623284 |
| Hbb-bh1 | 706.022 | 83.3092 |
| Hba-a1 | 1579.474 | 185.9253 |
| Xlr5a | 0.993692 | 0.113261 |
| Hbb-b1 | 529.409 | 60.2052 |
| Tspo2 | 2.04128 | 0.232130725 |
| Ccdc120 | 5.37323 | 0.608231 |
| Filip1l | 54438.22854 | 6022.36657 |
| Gt(ROSA)2( | 11.17788 | 1.234618 |
| Camp | 11.1273 | 1.22657 |
| Sprr2a3 | 1.47741 | 0.162848 |
| S100a9 | 35.0325 | 3.8526 |
| Atp6v0c-ps | 61.3178 | 6.66896 |
| Hist1h4c | 8.25718 | 0.89072 |
| Xlr4c | 2.00772 | 0.213156 |
| Gm8013 | 1.1437 | 0.120876 |
| Apoc2 | 3.36885 | 0.352615 |
| S100a8 | 65.7171 | 6.6435 |
| Rnase12 | 8.283049 | 0.801251468 |
| Gm14499 | 1.60169 | 0.152872 |
| Gm5483 | 9.34572 | 0.849423 |
| Stfa2l1 | 2.43883 | 0.198888 |
| Slc7a2 | 13.89492156 | 1.127888806 |
| Xlr4b | 3.904753256 | 0.311820486 |
| Rhox10 | 1.07679 | 0.0779682 |
| Tex12 | 4.96231 | 0.353645 |
| Gp9 | 3.34362 | 0.218673622 |
| Crygb | 1.59294 | 0.103631 |
| Hist2h3b | 3.94826 | 0.228673 |
| Hist1h3b | 4.50655 | 0.247092 |
| Hba-a2 | 1905.956 | 101.3926 |
| Gm6213 | 3.440826 | 0.142289601 |
| Stfa3 | 5.42074 | 0.202379 |
| Abhd1 | 1.30737 | 0.0442796 |
| Ereg | 2.839239 | 0.0936741 |
| Insl3 | 2.46645 | 0.0581884 |
| BC049730 | 7.70868 | 0.095365237 |
| Snx31 | 19.56441 | 0.159168709 |
| Gm17066 | 1.975 | 0.0150899 |
| Rsc1a1 | 4.04052 | 0.00382482 |
| Hbb-bs | 33.249 | 0.0166093 |

| | | |
|---|---|---|
| Lgals3 | 2.41141 | 0.000471906 |
| Ccdc84 | 3.82274 | 0 |
| Crh | 1.35086 | 0 |
| Cst11 | 1.3013 | 0 |
| Ddx17 | 99.42959 | 0 |
| Defb7 | 2.00935 | 0 |
| Figla | 1.50297 | 0 |
| Gm10336 | 2.85091 | 0 |
| Gm12409 | 1.244262901 | 0 |
| Gm3219 | 4.37084 | 0 |
| Hist1h2af | 1.30014 | 0 |
| Hist1h2an | 1.41019 | 0 |
| Hist1h2ba | 2.19001 | 0 |
| Hist1h4m | 3.66151 | 0 |
| Hist2h4 | 2.44884 | 0 |
| Hnrnpab | 308.1841262 | 0 |
| Kdelr3 | 28.0741 | 0 |
| Klk1b22 | 1.4444 | 0 |
| Mir344 | 1252.7 | 0 |
| Mir6236 | 75.579 | 0 |
| Phykpl | 3.93386 | 0 |
| Rnu12 | 14.632 | 0 |
| Rps25 | 3126.6616 | 0 |
| S100z | 1.50824 | 0 |
| Snora21 | 26.6594 | 0 |
| Snora33 | 288.963 | 0 |
| Snora52 | 13.5342 | 0 |
| Snora68 | 1679.57 | 0 |
| Snora7a | 48.3847 | 0 |
| Sprr2i | 1.45218 | 0 |
| Stfa1 | 15.135 | 0 |
| Terc | 1.47135 | 0 |
| Tmem265 | 4.83974 | 0 |
| Trappc4 | 58.5998 | 0 |
| Wfdc21 | 1.67987 | 0 |
| Xist | 24.44977 | 0 |

**lox XX gonads at E12.5**
