## Supplementary material for "*Lhx2* in germ cells suppresses endothelial cell migration in the developing ovary": Suppl Table 5

**Supplymentery Table 5: FPKM values of endothelial cell enriched genes in Wild type vs Lhx2 flox/flox XX gonads at E12.5**

| Gene Sym | XX WT | XY WT | XX *Lhx2* flox flox |
|---|---|---|---|
| 1810011O | 13.29481 | 9.72253 | 14.02731 |
| 2810025M | 106.853 | 49.0413 | 50.4024 |
| 4930578C1 | 0.101135 | 0.320888 | 0.169577 |
| Abca1 | 5.066784 | 3.12435 | 4.42565 |
| Abcb1a | 0.036761 | 0.051725 | 0.051725 |
| Abcb1b | 0.129217 | 0.107029 | 0.107029 |
| Abi3 | 1.09745 | 1.690949 | 2.017971 |
| Acer2 | 3.63432 | 4.683245 | 5.228245 |
| Ackr3 | 9.49995 | 8.10457 | 5.9597 |
| Acsl5 | 7.81513 | 6.83076 | 6.83076 |
| Acvrl1 | 3.729748 | 7.137177 | 5.552723 |
| Adam15 | 9.7577 | 5.76328 | 10.9161 |
| Adcy4 | 3.62605 | 2.813666 | 3.438187 |
| Adgre1 | 2.270947 | 2.040694 | 2.89318 |
| Adgre5 | 9.441267 | 7.086203 | 10.84602 |
| Adgrg3 | 1.201515 | 0.330483 | 0.718971 |
| Adgrl4 | 2.6002 | 1.89009 | 1.89009 |
| Adora2a | 4.290485 | 2.057654 | 3.745998 |
| Afap1l1 | 5.85667 | 5.27825 | 5.27825 |
| Ahr | 2.71956 | 2.18367 | 2.18367 |
| Ammecr1 | 2.86919 | 6.22188 | 4.10798 |
| Amotl1 | 5.360598 | 5.819742 | 8.218949 |
| Anxa3 | 15.701 | 11.3163 | 12.8334 |
| Ap1s2 | 8.7453 | 7.41605 | 5.817678 |
| Apbb2 | 2.527791 | 3.347134 | 3.249256 |
| Aplnr | 26.5198 | 9.47354 | 19.3718 |
| Aqp1 | 5.73549 | 3.42364 | 2.28444 |
| Arap3 | 4.021686 | 4.35786 | 7.351859 |
| Arhgap18 | 7.09904 | 5.62273 | 6.0107 |
| Arhgap25 | 0.605528 | 0.497636 | 0.476773 |
| Arhgap27 | 3.07528 | 2.425081 | 3.590719 |
| Arhgap28 | 9.60157 | 7.685628 | 13.79664 |
| Arhgap31 | 2.460639 | 2.260736 | 3.196038 |
| Arhgdib | 20.02854 | 12.60126 | 16.00267 |
| Arhgef15 | 3.64059 | 2.40531 | 4.572768 |
| Arhgef2 | 8.646879 | 15.91431 | 19.02594 |
| Arhgef28 | 2.100258 | 4.436873 | 3.332865 |
| Arhgef5 | 4.09519 | 4.00958 | 5.40409 |
| Arhgef7 | 11.76751 | 12.99645 | 17.34643 |
| Arl6ip5 | 37.5549 | 30.5033 | 31.5254 |
| Arpc3 | 159.0959 | 128.8822 | 111.9584 |
| Arpc5 | 66.6126 | 71.9818 | 47.9634 |
| Asah2 | 0.540695 | 1.128987 | 1.13959 |

| | | | |
|---|---|---|---|
| Asb4 | 18.90051 | 10.91477 | 12.29039 |
| Atp8b1 | 0.066951 | 0.156203 | 0.091596 |
| Atxn7l1 | 3.715822 | 3.881463 | 2.785367 |
| B3gnt2 | 8.68258 | 9.220048 | 5.4075 |
| Bach1 | 6.960884 | 7.707922 | 9.947757 |
| BC028528 | 22.50702 | 6.14844 | 9.10639 |
| Bcl2l11 | 5.397833 | 5.578928 | 7.451149 |
| Bcl6b | 2.68845 | 1.63087 | 3.40605 |
| Bmp2k | 4.872272 | 4.68688 | 4.462938 |
| C1qb | 20.4201 | 21.208 | 28.8643 |
| C3ar1 | 1.34559 | 1.14436 | 2.41116 |
| Cables1 | 16.18801 | 9.947477 | 11.33432 |
| Calcrl | 3.469652 | 1.701692 | 2.214685 |
| Camk2n1 | 6.028202 | 4.26256 | 8.28004 |
| Camk4 | 0.231652 | 0.253753 | 0.2319 |
| Casp7 | 6.8665 | 14.5085 | 8.60465 |
| Casp8 | 3.133248 | 9.10735 | 3.82838 |
| Cav1 | 60.03166 | 8.963838 | 21.66271 |
| Cav2 | 2.154039 | 1.44317 | 1.554254 |
| Ccdc149 | 0.484259 | 0.502551 | 0.578111 |
| Ccdc85a | 0.122045 | 0.143363 | 0.220606 |
| Ccl12 | 1.15152 | 15.5287 | 2.25301 |
| Ccm2l | 2.643722 | 1.124713 | 2.602633 |
| Ccnd1 | 22.5992 | 25.0476 | 23.2302 |
| Ccser1 | 0.5055 | 0.815659 | 0.428221 |
| Cd109 | 0.80141 | 0.740285 | 0.706217 |
| Cd34 | 36.61106 | 14.50237 | 23.42958 |
| Cd38 | 3.21255 | 2.0838 | 4.12802 |
| Cd40 | 1.936684 | 2.741361 | 2.250716 |
| Cd47 | 7.97362 | 9.707294 | 6.787961 |
| Cd93 | 4.41209 | 3.09493 | 4.63857 |
| Cdc42ep1 | 18.0835 | 14.2559 | 16.0728 |
| Cdc42ep3 | 6.59419 | 10.03021 | 6.69577 |
| Cdh5 | 11.7071 | 8.73642 | 13.1557 |
| Cdipt | 63.3342 | 37.75427 | 51.06334 |
| Cdk17 | 8.31251 | 8.60018 | 7.41832 |
| Cds2 | 11.09423 | 9.08508 | 12.25905 |
| Chd7 | 1.059689 | 1.635365 | 1.519328 |
| Cldn5 | 54.6133 | 20.792 | 29.8885 |
| Clec14a | 4.14926 | 2.59783 | 2.47243 |
| Clec1a | 0.259598 | 0.110595 | 0.212309 |
| Clec1b | 3.607919 | 0.54216 | 2.289415 |
| Clic1 | 200.3887 | 130.344 | 147.5512 |
| Clic4 | 31.9684 | 45.6115 | 33.5358 |
| Cmklr1 | 0.958365 | 1.588533 | 1.459653 |
| Cmtm3 | 124.978 | 57.7045 | 97.0489 |
| Cmtm8 | 6.06924 | 6.05377 | 5.73009 |

| | | | |
|---|---|---|---|
| Cnn2 | 71.6428 | 73.5528 | 51.2459 |
| Col15a1 | 6.594112 | 6.01963 | 10.89471 |
| Coro2b | 2.68364 | 3.15769 | 4.75174 |
| Cotl1 | 145.107 | 68.8075 | 93.2587 |
| Creb5 | 0.088812 | 0.459066 | 0.191981 |
| Crim1 | 1.89645 | 4.28454 | 3.18221 |
| Crip2 | 47.6333 | 28.42755 | 45.1231 |
| Crmp1 | 3.420483 | 4.10144 | 3.793378 |
| Crybg3 | 2.833698 | 1.806825 | 2.583966 |
| Ctla2a | 9.914856 | 2.72881 | 4.284061 |
| Ctla2b | 3.32355 | 2.78996 | 1.47678 |
| Cxcr4 | 7.672301 | 9.674134 | 9.248091 |
| Cyth4 | 2.403442 | 2.015758 | 3.296372 |
| Cyyr1 | 0.421962 | 0.236199 | 0.571483 |
| Dach1 | 3.004264 | 1.345783 | 4.667352 |
| Dazap2 | 72.0165 | 61.4117 | 68.7026 |
| Dennd4a | 0.570669 | 0.714304 | 1.010431 |
| Dgkh | 0.47255 | 0.566094 | 0.582902 |
| Dkk2 | 1.99039 | 0.683088 | 1.55809 |
| Dlc1 | 7.469452 | 7.791433 | 10.81768 |
| Dock11 | 5.501649 | 4.544742 | 4.321138 |
| Dock6 | 21.51183 | 16.54862 | 23.87943 |
| Dock8 | 0.472684 | 0.684693 | 0.932684 |
| Dok4 | 14.591 | 10.9194 | 11.5335 |
| Dpysl3 | 7.97098 | 8.14108 | 11.29311 |
| Dusp3 | 3.486271 | 2.973411 | 3.048761 |
| Dysf | 1.772245 | 1.979444 | 2.22256 |
| Ebf3 | 1.24689 | 2.636161 | 0.902612 |
| Ecm2 | 0.109681 | 0.153656 | 0.110869 |
| Ecscr | 18.475 | 8.9056 | 8.361186 |
| Ednrb | 4.637853 | 1.956884 | 3.078749 |
| Efnb2 | 7.95229 | 11.6876 | 16.4318 |
| Egfl7 | 44.54343 | 28.47455 | 29.6254 |
| Ehd2 | 20.57042 | 15.04848 | 18.42775 |
| Eng | 13.1205 | 9.940647 | 22.25509 |
| Eogt | 2.46837 | 2.729 | 2.6055 |
| Epas1 | 3.30358 | 4.64763 | 2.47338 |
| Erf | 13.45345 | 22.27068 | 24.94402 |
| Erg | 0.719202 | 1.441057 | 1.084145 |
| Esam | 23.52533 | 9.434918 | 14.95174 |
| Esm1 | 0.513326 | 1.20447 | 1.07573 |
| Ets2 | 17.628 | 11.9323 | 16.143 |
| Exoc3l | 3.11829 | 5.93098 | 5.3995 |
| Exoc5 | 9.19097 | 10.3014 | 5.90247 |
| Exoc6 | 2.31827 | 2.99363 | 1.690235 |
| F2r | 24.66798 | 20.75242 | 26.09663 |
| Fabp4 | 25.051 | 2.81839 | 9.453 |

| | | | |
|---|---|---|---|
| Fam102b | 2.91391 | 3.83213 | 3.59634 |
| Fam107b | 9.82258 | 8.244951 | 10.5738 |
| Fam124b | 0.112481 | 0.144885 | 0.423797 |
| Fam129b | 19.50969 | 17.02183 | 22.71296 |
| Fam13c | 0.480866 | 0.590249 | 1.100341 |
| Fam167b | 4.51651 | 2.46885 | 4.40027 |
| Fam43a | 9.62889 | 5.53505 | 6.08455 |
| Fam57a | 17.61135 | 6.427969 | 8.471518 |
| Fcer1g | 17.2447 | 19.3906 | 23.0528 |
| Fhl3 | 7.51022 | 10.221 | 9.888267 |
| Filip1 | 0.512428 | 0.700858 | 0.573054 |
| Fli1 | 3.118378 | 4.289524 | 2.781496 |
| Flnb | 5.921965 | 15.23642 | 13.93601 |
| Flt1 | 1.585876 | 1.055205 | 2.120527 |
| Flt4 | 1.05699 | 0.81938 | 2.31958 |
| Fmnl3 | 12.48627 | 18.01045 | 16.95849 |
| Frmd4a | 3.746996 | 6.127726 | 6.392767 |
| Fscn1 | 100.1126 | 133.828 | 148.0224 |
| Fxyd5 | 8.342206 | 5.121009 | 8.087551 |
| Fyb | 0.379173 | 0.493465 | 0.649016 |
| Fzd4 | 4.23678 | 2.57462 | 4.63269 |
| Fzd6 | 1.377874 | 1.485468 | 3.125889 |
| Gap43 | 13.38064 | 7.84934 | 7.8627 |
| Gbp9 | 0.396984 | 3.123713 | 3.196112 |
| Gdpd3 | 3.9127 | 0.339156 | 0.778939 |
| Gdpd5 | 4.0639 | 3.394136 | 5.040402 |
| Ggta1 | 8.190944 | 4.309195 | 7.821373 |
| Gimap1 | 5.59276 | 3.262123 | 4.90582 |
| Gimap4 | 13.92184 | 2.746022 | 8.32348 |
| Gimap6 | 12.5447 | 4.74949 | 7.12101 |
| Gimap8 | 0.516752 | 0.251091 | 0.545902 |
| Gimap9 | 5.4772 | 3.12916 | 2.186178 |
| Git2 | 4.311217 | 9.006612 | 7.874043 |
| Gja5 | 2.126759 | 4.013853 | 1.790431 |
| Gm10791 | 0.250011 | 0 | 0.267947 |
| Gm14005 | 0.40362 | 1.835274 | 0.123763 |
| Gnai2 | 120.565 | 129.276 | 154.15 |
| Gnb4 | 12.4503 | 16.0828 | 11.5605 |
| Gng11 | 64.7673 | 27.7069 | 25.1244 |
| Gng2 | 19.21589 | 12.68719 | 17.3621 |
| Gngt2 | 7.79141 | 2.92233 | 4.50018 |
| Gpihbp1 | 7.11261 | 0.281536 | 4.58587 |
| Gpr183 | 0.929489 | 0.761648 | 0.829368 |
| Gramd1a | 14.67832 | 19.60459 | 27.31636 |
| Gria2 | 0.122557 | 0.077677 | 0.371197 |
| Gulp1 | 4.80833 | 6.8396 | 4.10824 |
| H2-K1 | 5.541667 | 20.82565 | 21.30717 |

| | | | |
|---|---|---|---|
| Hapln1 | 4.38742 | 2.07939 | 3.11976 |
| Hbegf | 2.636042 | 1.745192 | 2.252869 |
| Hcls1 | 1.48 | 2.60483 | 2.68835 |
| Hdac7 | 14.31337 | 10.82165 | 22.01538 |
| Heg1 | 4.274636 | 4.37243 | 7.43946 |
| Hey1 | 15.9035 | 26.6009 | 15.7561 |
| Hhex | 3.680602 | 2.35656 | 2.631177 |
| Hivep2 | 0.491645 | 0.417075 | 0.814062 |
| Hlx | 5.26008 | 4.87224 | 6.15926 |
| Hoxb4 | 1.42533 | 1.13458 | 2.41867 |
| Hoxb5 | 1.80606 | 1.47606 | 4.28471 |
| Hpgd | 9.021346 | 36.8603 | 8.28631 |
| Hspg2 | 6.685616 | 9.045662 | 16.57563 |
| Ica1 | 4.007208 | 3.080242 | 3.406477 |
| Icam2 | 23.0093 | 11.3083 | 11.2739 |
| Ifi203 | 0.165351 | 1.041889 | 0.816157 |
| Ifngr1 | 7.47656 | 6.97198 | 5.42408 |
| Igfbp3 | 105.259 | 100.595 | 231.551 |
| Il1r1 | 1.59924 | 1.336917 | 2.245366 |
| Il2rg | 1.604552 | 1.882714 | 2.44628 |
| Inpp4b | 0.391292 | 0.256216 | 0.21951 |
| Inpp5d | 1.25163 | 1.790786 | 2.116874 |
| Itga1 | 1.304439 | 7.991965 | 2.559781 |
| Itga2 | 0.625162 | 0.226524 | 0.617467 |
| Itga5 | 7.1132 | 9.94644 | 12.9532 |
| Itpripl2 | 2.73773 | 3.93032 | 3.23834 |
| Itsn2 | 2.831494 | 4.457267 | 3.63116 |
| Jmjd1c | 2.58405 | 3.55142 | 3.586863 |
| Kbtbd11 | 0.836196 | 0.772841 | 1.526 |
| Kcna5 | 2.05355 | 2.31125 | 2.18641 |
| Kcne3 | 4.219001 | 3.133354 | 4.048828 |
| Kctd12b | 2.872369 | 1.34073 | 2.275541 |
| Kdm6b | 1.903523 | 8.71313 | 11.39861 |
| Kdr | 6.3867 | 5.23343 | 11.8142 |
| Kif1c | 7.700859 | 10.90314 | 13.87171 |
| Klf3 | 7.276212 | 7.190758 | 7.097545 |
| Klf6 | 7.66667 | 7.15123 | 5.19132 |
| Klhl4 | 5.806263 | 3.598857 | 3.432549 |
| Klhl6 | 5.52913 | 1.63055 | 5.32038 |
| Lama4 | 29.82326 | 23.64917 | 16.9278 |
| Lcp1 | 6.06607 | 5.031267 | 6.46802 |
| Ldb2 | 15.79655 | 14.42213 | 17.35416 |
| Lepr | 0.40866 | 0.25744 | 0.301011 |
| Limch1 | 2.315977 | 2.609353 | 2.435715 |
| Lmbr1 | 2.881919 | 1.74119 | 2.46147 |
| Lmo2 | 19.40891 | 7.882167 | 12.49255 |
| Lpar4 | 3.2628 | 2.78562 | 1.65662 |

| | | | |
|---|---|---|---|
| Lpar6 | 5.25415 | 6.5995 | 7.75045 |
| Lrch1 | 1.610253 | 2.702223 | 2.51578 |
| Lrrc8c | 2.07966 | 3.454199 | 2.573423 |
| Lxn | 22.6973 | 16.4261 | 14.6457 |
| Lyn | 3.286254 | 4.127091 | 2.760413 |
| Madcam1 | 5.18375 | 1.215048 | 2.876168 |
| Magi3 | 2.432463 | 2.767442 | 2.902438 |
| Mal | 2.28058 | 1.06022 | 2.30686 |
| Maml2 | 1.711284 | 1.037368 | 1.790042 |
| Map2k6 | 9.03822 | 7.89087 | 9.65591 |
| Map3k11 | 9.09852 | 11.9971 | 14.98029 |
| Map4k2 | 3.28621 | 5.366369 | 6.904743 |
| Map4k5 | 7.75545 | 10.4022 | 9.56662 |
| Mcam | 14.42999 | 17.5221 | 11.5451 |
| Mecom | 1.579077 | 1.217422 | 2.159714 |
| Mef2c | 2.884243 | 5.456533 | 2.738995 |
| Mef2d | 5.17781 | 13.06205 | 12.72751 |
| Mest | 214.2032 | 219.5295 | 286.9528 |
| Mfng | 14.3816 | 5.7828 | 9.1765 |
| Mmrn1 | 1.527159 | 0.962958 | 0.956829 |
| Mmrn2 | 3.64973 | 3.13159 | 4.00679 |
| Mospd1 | 8.518491 | 6.467008 | 5.62754 |
| Mpeg1 | 1.16636 | 3.619961 | 2.771999 |
| Mpp1 | 19.8591 | 14.294 | 16.3587 |
| Mpzl1 | 66.8882 | 46.08344 | 65.01902 |
| Ms4a6d | 3.33038 | 2.06419 | 2.24918 |
| Msn | 20.6054 | 30.3016 | 28.0248 |
| Msrb3 | 3.164367 | 5.085902 | 4.094179 |
| Myc | 7.492998 | 8.57711 | 11.25986 |
| Myct1 | 3.00775 | 1.11627 | 2.13483 |
| Myo1b | 11.35014 | 14.09378 | 12.42548 |
| Myo1e | 3.393469 | 5.44769 | 6.37905 |
| Myo6 | 7.59045 | 7.14095 | 4.28965 |
| Myzap | 2.75983 | 4.800261 | 2.645181 |
| N4bp3 | 6.56407 | 9.516449 | 8.867002 |
| Nav3 | 0.497663 | 0.210693 | 0.393976 |
| Nckap5 | 0.365252 | 0.514605 | 0.256557 |
| Ndnf | 1.419215 | 1.523327 | 2.21876 |
| Nfkbia | 14.6053 | 15.7028 | 14.7065 |
| Nmi | 3.19305 | 12.89033 | 5.195546 |
| Notch1 | 2.21477 | 2.02689 | 3.83002 |
| Notch4 | 1.33308 | 1.00007 | 1.97346 |
| Nox4 | 1.348494 | 1.347242 | 1.416228 |
| Npr3 | 1.468107 | 0.962835 | 1.567985 |
| Nr4a2 | 0.236563 | 1.228181 | 0.595759 |
| Nrarp | 7.28277 | 5.49942 | 6.19639 |
| Nrp1 | 14.15906 | 3.48635 | 7.894089 |

| | | | |
|---|---|---|---|
| Nrp2 | 2.515868 | 3.567945 | 3.005242 |
| Nrros | 7.083534 | 5.824243 | 4.952463 |
| Oit3 | 2.46158 | 0.96401 | 2.02487 |
| Pak2 | 11.3774 | 17.552 | 15.2323 |
| Palmd | 0.613034 | 1.89039 | 1.08893 |
| Pamr1 | 3.34031 | 0.176591 | 3.25055 |
| Pcdh1 | 4.505196 | 2.313779 | 4.351023 |
| Pcdh12 | 2.997023 | 1.568065 | 2.54421 |
| Pcdh17 | 0.565599 | 0.505966 | 0.557972 |
| Pcdhb16 | 0.435571 | 0.324012 | 0.627531 |
| Pcdhb17 | 1.77563 | 1.29217 | 1.928 |
| Pde4b | 1.32269 | 2.156023 | 1.562435 |
| Pde4d | 1.139163 | 1.08132 | 1.208689 |
| Pdgfb | 4.17988 | 4.746913 | 4.6153 |
| Pdlim5 | 3.57519 | 6.371064 | 3.661436 |
| Pea15a | 29.2823 | 18.7426 | 18.1484 |
| Pgm1 | 11.1419 | 11.1021 | 9.8457 |
| Phactr2 | 2.831698 | 2.214898 | 4.860992 |
| Piezo2 | 0.835231 | 0.84095 | 1.515975 |
| Pik3cg | 0.327105 | 0.265991 | 0.357391 |
| Pitpna | 23.88206 | 26.61501 | 22.18749 |
| Pitpnc1 | 2.044385 | 3.81261 | 4.103718 |
| Plagl1 | 21.05175 | 10.42737 | 22.3546 |
| Plekhg1 | 2.587 | 1.484427 | 3.529435 |
| Plekho1 | 40.34271 | 41.5502 | 36.80821 |
| Plk2 | 16.4708 | 11.4042 | 18.6949 |
| Plvap | 34.8496 | 16.2662 | 28.31955 |
| Plxnd1 | 50.34295 | 35.3704 | 49.94918 |
| Pnp | 18.4183 | 6.78558 | 10.9583 |
| Pon2 | 7.32945 | 8.199981 | 5.790817 |
| Pon3 | 5.035946 | 1.714644 | 2.291576 |
| Ppap2a | 12.73827 | 11.05376 | 9.19356 |
| Ppdpf | 209.932 | 157.004 | 163.829 |
| Ppp1r2 | 18.9767 | 18.2034 | 12.9376 |
| Prcp | 13.2414 | 8.23009 | 11.25699 |
| Prex2 | 0.591481 | 0.646924 | 0.537531 |
| Prkcdbp | 49.348 | 38.2829 | 33.8317 |
| Prkch | 4.509042 | 2.155601 | 2.847202 |
| Prkd2 | 4.237672 | 5.126028 | 6.82826 |
| Prkx | 2.3696 | 2.72209 | 3.40632 |
| Procr | 10.0005 | 5.1483 | 7.26318 |
| Pros1 | 4.27803 | 5.53433 | 6.28796 |
| Ptp4a3 | 20.16726 | 19.21517 | 18.2341 |
| Ptprb | 0.715956 | 0.43608 | 0.610688 |
| Ptprm | 2.933792 | 2.205465 | 2.840075 |
| Ptrf | 28.4506 | 43.4219 | 29.5815 |
| Pxdc1 | 2.779403 | 4.850707 | 2.744467 |

| | | | |
|---|---|---|---|
| Ralb | 58.10238 | 58.38115 | 43.83564 |
| Rapgef5 | 1.777045 | 1.275271 | 1.705257 |
| Rasgrp2 | 8.418459 | 3.179746 | 5.012972 |
| Rasgrp3 | 2.141886 | 2.292244 | 2.468067 |
| Rasip1 | 11.831 | 7.07824 | 12.3446 |
| Rassf2 | 1.521894 | 1.981569 | 2.724761 |
| Rbms1 | 16.00788 | 21.59218 | 16.1358 |
| Rbpms | 35.3132 | 72.9166 | 49.60349 |
| Rell1 | 4.44759 | 6.13035 | 5.53819 |
| Rftn2 | 4.410585 | 2.609962 | 4.354933 |
| Rgl1 | 9.23138 | 9.848038 | 12.37473 |
| Rgs16 | 1.7872 | 2.50785 | 1.5966 |
| Rhoc | 222.4205 | 77.16207 | 114.119 |
| Rhoj | 20.5063 | 5.33012 | 14.9547 |
| Rnasek | 82.4541 | 109.187 | 91.9535 |
| Robo4 | 4.48698 | 2.468073 | 5.25816 |
| Rsad2 | 0.957578 | 24.4968 | 4.88569 |
| Rtp3 | 0.129511 | 0.081049 | 0.086818 |
| Rtp4 | 0.934538 | 49.6558 | 15.8761 |
| S100a13 | 44.95329 | 13.90699 | 16.01038 |
| S100a16 | 111.203 | 16.6523 | 41.82 |
| Samsn1 | 0.416509 | 0.597205 | 0.453939 |
| Sdpr | 4.09309 | 3.33295 | 2.06424 |
| Sema4c | 7.748543 | 8.34773 | 14.8072 |
| Sema6a | 2.27692 | 2.84964 | 5.99957 |
| Sepp1 | 147.349 | 79.61613 | 114.6221 |
| Sgk1 | 8.287923 | 7.379387 | 10.52719 |
| Sh2d3c | 6.31293 | 6.303371 | 11.89167 |
| Sh3bgrl3 | 148.175 | 72.922 | 116.109 |
| Sh3glb1 | 16.69354 | 20.2636 | 13.69778 |
| Sh3pxd2b | 6.26844 | 5.84485 | 10.9216 |
| Sh3tc1 | 1.453398 | 1.708648 | 2.294875 |
| Shank3 | 2.285808 | 2.282245 | 4.181054 |
| She | 2.4396 | 1.67105 | 2.05483 |
| Shroom2 | 6.205413 | 5.612492 | 5.60261 |
| Sipa1 | 6.969076 | 16.63296 | 16.02164 |
| Slc15a2 | 1.235265 | 0.644064 | 1.033493 |
| Slc26a10 | 0.991967 | 0.136919 | 0.852597 |
| Slc2a1 | 34.9578 | 39.05985 | 41.85432 |
| Slc43a3 | 16.10032 | 13.25858 | 16.31327 |
| Slco2a1 | 0.706065 | 0.287147 | 0.958705 |
| Slfn5 | 1.154724 | 3.023935 | 2.025632 |
| Smad1 | 5.67585 | 9.19421 | 8.20715 |
| Smagp | 16.5175 | 8.55678 | 8.005527 |
| Snap29 | 7.87482 | 5.34274 | 6.24256 |
| Sncg | 4.3911 | 3.64254 | 2.05319 |
| Snrk | 7.675194 | 7.052494 | 8.865999 |

| | | | |
|---|---|---|---|
| Sntb1 | 0.503723 | 1.231704 | 0.50478 |
| Sox17 | 3.296642 | 2.853231 | 2.708534 |
| Sox18 | 23.1446 | 16.7694 | 19.9235 |
| Sox7 | 2.710294 | 2.69361 | 3.571959 |
| Spag9 | 7.794803 | 8.548869 | 8.942294 |
| Sparcl1 | 64.78211 | 26.00068 | 42.17828 |
| Spata6 | 8.41641 | 5.11474 | 5.769875 |
| Sqrdl | 5.421986 | 1.213201 | 3.286485 |
| Srgn | 4.746646 | 4.172991 | 2.280797 |
| Ssh2 | 1.803942 | 1.975109 | 1.927559 |
| St7 | 7.633171 | 6.495813 | 7.103354 |
| St8sia4 | 1.57721 | 0.70599 | 1.345766 |
| St8sia6 | 0.073376 | 0.030184 | 0.114174 |
| Stab1 | 4.33237 | 1.80445 | 5.20382 |
| Stab2 | 0.610211 | 0.054117 | 0.638231 |
| Stap2 | 4.346816 | 1.910425 | 3.296565 |
| Stard13 | 0.75924 | 0.98234 | 1.254277 |
| Stard4 | 5.54379 | 6.947047 | 6.211987 |
| Stim2 | 3.96333 | 4.73965 | 3.24719 |
| Stk10 | 2.549844 | 4.068358 | 2.83489 |
| Stmn2 | 13.45973 | 6.25756 | 6.41503 |
| Stx2 | 8.35588 | 6.976436 | 9.31936 |
| Stx6 | 32.5749 | 25.5335 | 28.8984 |
| Tagln2 | 59.5154 | 61.6213 | 51.6215 |
| Tcf4 | 10.98233 | 12.07387 | 11.65709 |
| Tead4 | 0.637863 | 1.941268 | 1.19127 |
| Tek | 1.989579 | 1.128582 | 1.644254 |
| Tfpi | 29.19014 | 16.78953 | 22.22455 |
| Tgfb1 | 5.95289 | 12.8351 | 12.7058 |
| Tgfbr2 | 6.870971 | 4.82648 | 8.76653 |
| Thsd1 | 2.884702 | 1.770943 | 3.284103 |
| Thsd7a | 0.236665 | 0.220072 | 0.409239 |
| Tie1 | 6.9344 | 6.11916 | 11.5482 |
| Tifa | 1.565835 | 3.068703 | 3.41991 |
| Tjp1 | 27.60455 | 30.20392 | 20.20821 |
| Tlr3 | 0.349692 | 1.3896 | 0.334786 |
| Tlr4 | 0.32428 | 0.165933 | 0.326532 |
| Tm4sf1 | 6.54654 | 3.31968 | 2.97694 |
| Tm6sf1 | 1.37616 | 1.607219 | 1.78 |
| Tmem204 | 8.13993 | 3.64282 | 6.00724 |
| Tmem88 | 18.2958 | 5.98499 | 13.4039 |
| Tnfaip2 | 1.18399 | 1.34225 | 1.5039 |
| Tnfaip3 | 0.459432 | 0.755746 | 0.636188 |
| Tnfrsf22 | 0.774829 | 0.309443 | 1.1251 |
| Tram2 | 1.169327 | 5.07741 | 3.52025 |
| Trf | 5.775126 | 43.46379 | 5.406827 |
| Trim16 | 1.08949 | 1.20437 | 1.2525 |

| | | | |
|---|---|---|---|
| Trim3 | 6.464234 | 8.5086 | 9.653942 |
| Trp53i11 | 8.934122 | 14.83704 | 14.6159 |
| Trpc3 | 1.25564 | 2.78202 | 0.604766 |
| Tspan14 | 34.03284 | 29.60672 | 33.26423 |
| Tspan18 | 7.367083 | 7.475062 | 7.331954 |
| Tspan6 | 78.2564 | 44.6246 | 49.0477 |
| Ttc28 | 1.694656 | 3.95515 | 5.111675 |
| Tubb6 | 21.9448 | 30.701 | 25.0803 |
| Tyrobp | 32.4809 | 18.9386 | 27.7532 |
| Uaca | 6.526075 | 9.777314 | 8.659178 |
| Ufm1 | 9.57863 | 7.954296 | 5.89122 |
| Unc5b | 5.144755 | 7.63515 | 9.889095 |
| Upp1 | 6.72107 | 2.869608 | 4.17783 |
| Ushbp1 | 2.90919 | 0.634103 | 1.712023 |
| Uvrag | 9.03185 | 7.053161 | 6.748743 |
| Vamp5 | 8.610141 | 6.24279 | 4.72591 |
| Vash1 | 4.60255 | 5.4404 | 8.03833 |
| Vav3 | 4.627598 | 0.924201 | 3.304845 |
| Vegfc | 4.767813 | 2.471878 | 2.956046 |
| Wipf1 | 3.698525 | 5.148517 | 3.839377 |
| Yes1 | 6.23572 | 6.783776 | 4.81282 |
| Zc3h7a | 4.0418 | 6.798734 | 6.34806 |
| Zdhhc20 | 16.45941 | 12.5562 | 11.50137 |
| Zfp608 | 1.782622 | 2.298425 | 3.722827 |
| Zfp69 | 2.099773 | 0.913612 | 1.289417 |
| Zfp711 | 2.640754 | 1.432523 | 2.188617 |
